# DNA Encoded Glycan Libraries as a next-generation tool for the study of glycan-protein interactions

**DOI:** 10.1101/2020.03.30.017012

**Authors:** Shukkoor M. Kondengaden, Jiabin Zhang, Huajie Zhang, Aishwarya Parameswaran, Shameer M. Kondengadan, Shrikant Pawar, Akhila Puthengot, Rajshekhar Sunderraman, Jing Song, Samuel J. Polizzi, Liuqing Wen, Peng George Wang

**Affiliations:** Department of Chemistry and Center for Diagnostics and Therapeutics, Georgia State University, Atlanta, GA 30303, USA; Department of Biology, Georgia State University, Atlanta, GA 30303, USA; Robinson College of Business, Georgia State University, Atlanta, GA 30303, USA; Department of Computer Science, Georgia State University, Atlanta, GA 30303, USA; Chemily LLC, Peachtree Corners, GA 30092, USA; Department of Chemistry, Southern University of Science and Technology, Shenzhen, Guangdong 518055, P. R. China

**Keywords:** DNA encoded glycan library, Multivalent glycan-DNA, Multiplex glycan assay, DNA glycan conjugates

## Abstract

Interactions between glycans and glycan-binding proteins (GBPs) mediate diverse cellular functions, and therefore are of diagnostic and therapeutic significance. Current leading strategies for studying glycan-GBP interactions require specialized knowledge and instrumentation. In this study, we report a strategy for studying glycan-GBP interactions that uses PCR, qPCR and next-generation sequencing (NGS) technologies that are more routinely accessible. Our headpiece conjugation-code ligation (HCCL) strategy couples glycans with unique DNA codes that specify glycan sugar moieties and glycosidic linkages when sequenced. We demonstrate the technology by synthesizing a DNA encoded glycan library of 50 biologically relevant glycans (DEGL-50) and probing interactions against 25 target proteins including lectins and antibodies. Data show glycan-GPB interactions in solution that are consistent with lower content, lower throughput ELISA assays. Data further demonstrate how monovalent and multivalent headpieces can be used to increase glycan-GPB interactions and enrich signals while using smaller sample sizes. The flexibility of our modular HCCL strategy has potential for producing large glycan libraries, facilitating high content-high throughput glycan binding studies, and increasing access to lower cost glyco-analyses.

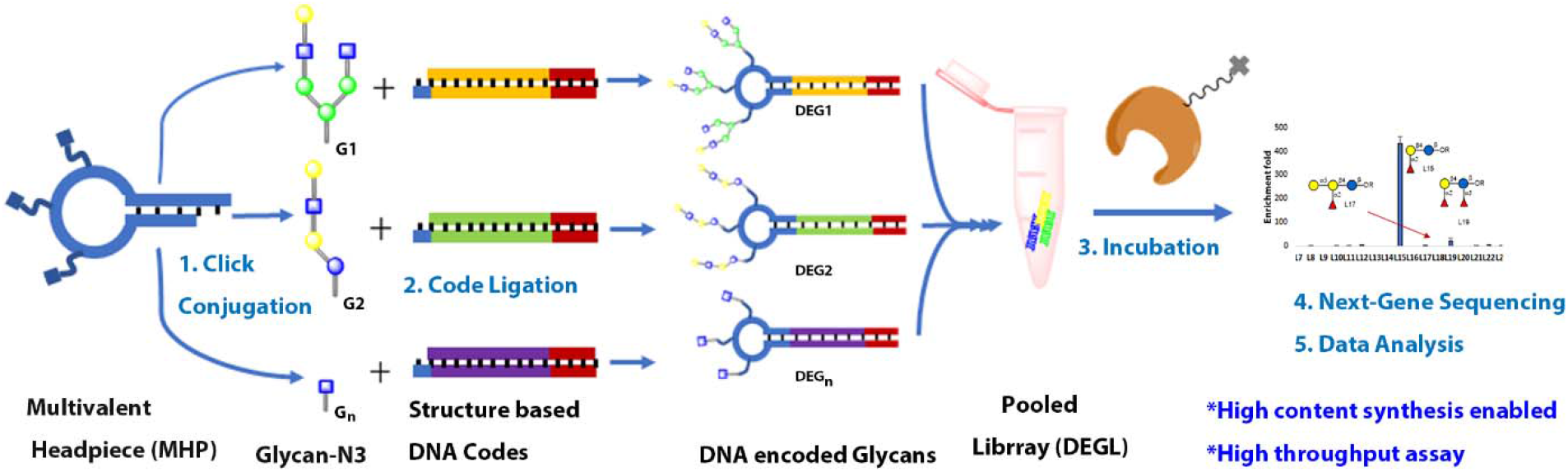

## Introduction

Glycans are abundant biomolecules that interact with glycan-binding proteins (GBPs) in diverse cellular functions such as molecular recognition, adhesion, pathogenesis, and inflammation.[1, 2] Glycan expression patterns are widely altered in cancer, retrovirus infection, atherosclerosis, thrombosis, diabetes, neurodegeneration, arthritis and other diseases.[3] Therefore, the interactions between glycans and GBPs are of particular interest for life science and pharmaceutical applications.[3, 4] Historically, the study of glycan-GBP interactions has been slowed by the weak nature of the interactions, and compounded by difficulties synthesizing glycans in appropriate quantities and purities. Contemporary methods to study glycan-GBP interactions with high sensitivity and low sample consumption are desirable to advance the field.

Currently, ELISA and glycan microarray are the prominent methods for glycan-GBP studies [4–11]. In ELISA studies, single types of glycans can be immobilized, allowing individual GBPs to be screened against glycans of known identity. While ELISA is a highly reliable solution phase assay used in many labs, it is less amenable to high-content analysis. Similarly, microarrays use immobilized glycans on a flat surface that can be incubated with GBPs. The microarray format allows a higher number of known glycans to be screened in one experiment, but requires specialized printing/reading instrumentation and sometimes cannot reflect the real binding conditions.[2, 6, 12, 13] More recently, a method called multiplex glycan bead array was reported to use colored luminex beads for high-throughput and high-content detection of GBP interactions. [9] However, this strategy also required special instrumentation, and detection was limited by low commercial availability of differently colored beads. The above methods have facilitated the study of limited glycan-GBP interactions, but leave room for improvements in the areas of high-content and high-throughput analyses.

In other related fields, high-content and high-throughput analyses have been aided by libraries of DNA encoded molecules. For example, DNA encoded libraries (DEL) of small molecules are gaining importance in drug screening.[14] By linking each small molecule with a unique piece of DNA, low quantities of many small molecules can be screened for binding in a single experiment, and the detection range of the small molecules can be enhanced many-fold by polymerase chain reaction (PCR) using standard molecular biology equipment.[15] Further, next generation sequencing (NGS) technologies are available for scaling to high-throughput and quantitative detection of 10^6^ to 10^9^ small molecules.[16–18]

DNA-glycan conjugates have previously been synthesized and utilized for high-sensitivity detection of glycans in vaccine design.[19–23] Kwon and co-workers also showed the feasibility of using Glyco-PCR to detect GBPs with high sensitivity, though the method was limited to singleplex assay.[24, 25] More recently, Song’s group demonstrated the application of NGS in glycan interaction studies.[26] Song’s work established the feasibility of DNA encoding and NGS in multiplex glycan interaction studies, while the glycan-DNA conjugation method used may limit the accessibility to limited glycan content and thereby compromise the true potential DNA encoding. Advances in the synthesis of carbohydrates via organic, enzymatic, chemo-enzymatic and machine-driven synthesis [27] have potential to eliminate the bottleneck in glycan interaction studies, i.e., availability of glycan probes. These new developments necessitate novel approaches for high-content and high-throughput methods for glycan interaction studies.

Here we developed a DNA encoded glycan library (DEGL) demonstrating higher content and throughput potential. This strategy uses routine molecular biology lab technologies to probe the interactions of glycans with target proteins. Contrary to libraries exploring undefined chemical spaces, the goal of our DEGL was to make a library of known glycans. Therefore, we employed new synthetic methodologies to define and encode the diversity of glycan structures. First, we developed a unique DNA coding system for glycans. Next, we devised a protocol for attaching the unique DNA codes to monovalent or multivalent headpieces, respectively containing single or multiple copies of each encoded glycan. This modular headpiece conjugation-code ligation (HCCL) system can be applied to picomolar amounts of each glycan and has the capacity to encode thousands of unique glycans using a single protocol. We synthesized a DEGL of 50 glycan epitopes (DEGL-50) and screened the library against 25 GBPs including lectins and antibodies in a single NGS experiment. Contributions to the field include a publicly available program for encoding and decoding DNA sequences corresponding to distinct glycans (http://tinman.cs.gsu.edu/~raj/carb/carb.html), and a method for glycan-GBP analysis that is low in sample consumption and high in content, throughput and sensitivity.

## RESULTS

### Unique DNA coding of glycans

Unlike proteins and nucleic acids, carbohydrates are not naturally defined by a template or synthesized by a coding scheme. As a result, glycans can display a wide diversity of sugar and non-sugar residue combinations, linkages, stereochemistry and positions. In order to create a template or coding scheme that would account for such a diversity of glycan representations, we developed a computer program GLYCO DNA Encoder and Reader (GLYCODER) to transform IUPAC names [28] into single-stranded DNA codes (available at http://tinman.cs.gsu.edu/~raj/carb/carb.html). This program utilized definition lists corresponding to the different parts of an IUPAC name. Separate lists were created to define glycan monomers (e.g. galactose), 46 glycan linkages (e.g. alpha 1→3), and over 100 non-sugar residues (Supporting Info, Table S4, S5, S6). Together, the definition lists were used to iteratively encode glycans using a common format of sugar residue (i.e. 4 DNA bases), linkage (i.e. 3 bases), and the next sugar or non-sugar residue (i.e. 4 bases), etc., until the IUPAC name terminated. The resulting string of DNA bases was unique to the glycan but also based in definitions and a general formula that can be used to build more complex branching glycans (Figure 1).

**Figure 1.**
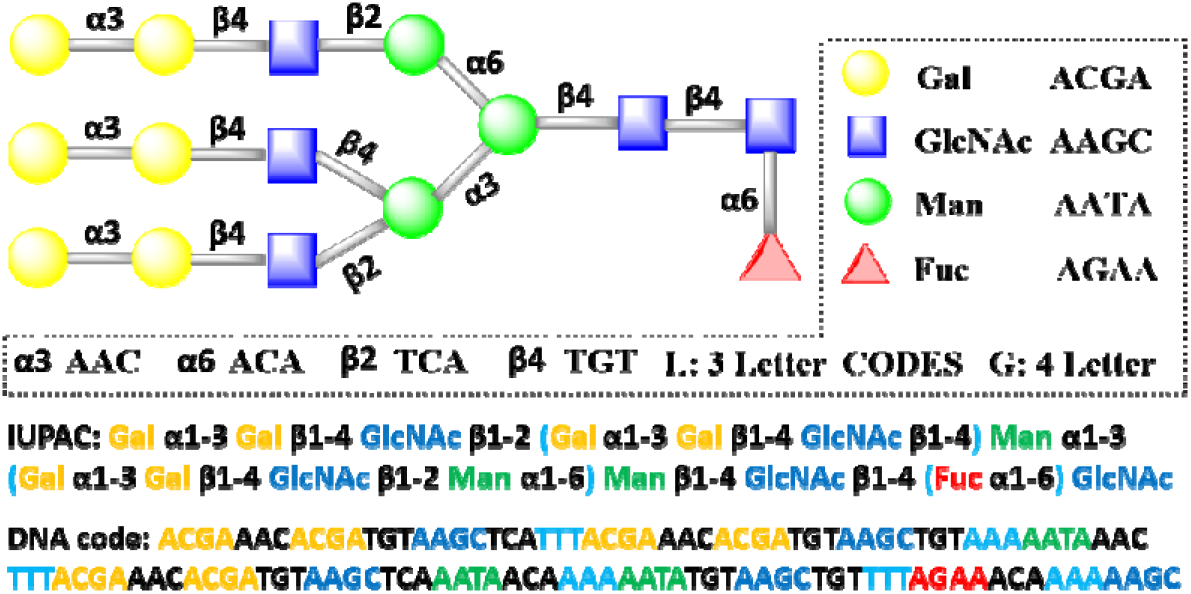
Assignment of a branched glycan using DNA code. Glycan structure is divided into sugar residues or linkages, which are coded with specific DNA ‘codons’ of length four or three, respectively. A full length DNA code is generated from the IUPAC name.

Recognizing that a template length of 40-120 bases can be ideal for PCR-based applications, we built length requirements into the GLYCODER. Short glycans encoded by <20 bases were automatically assigned a 20 base primer sequence that was constant and distinct from glycan codes. Longer glycans encoded by >120 bases were automatically condensed using an additional list of definitions based on common glycan structures, such as the hexasaccharide core structure of N-glycans (Supporting Info, Table S7), and thus limited the overall base length. In addition to converting a user supplied IUPAC sugar name into a unique DNA code, the publicly available program can also convert DNA sequences into sugar names after NGS analysis of DEGL samples.

### Conjugation and display strategy for DNA encoded glycans

Recent advances in synthetic carbohydrate chemistry have resulted in a rapid increase in the number and diversity of glycans available for study. However, the quantity of each glycan often remains low due to synthesis occurring at an analytical scale. Therefore, methods for creating DEGL should ideally be adaptable, high yielding, and suitable for sub-nanomole scale synthesis. Methods for direct conjugation of DNA to glycans has been achieved via click chemistry, amine-reactive (NHS) crosslinking, and reductive amination [3, 19, 24, 29], but often requires unique DNA codes to be custom synthesized with specific chemical linkers (Figure 2a). Direct conjugation strategies are limited in adaptability and cost-effectiveness and represent barriers to creating large DEGL.

**Figure 2.**
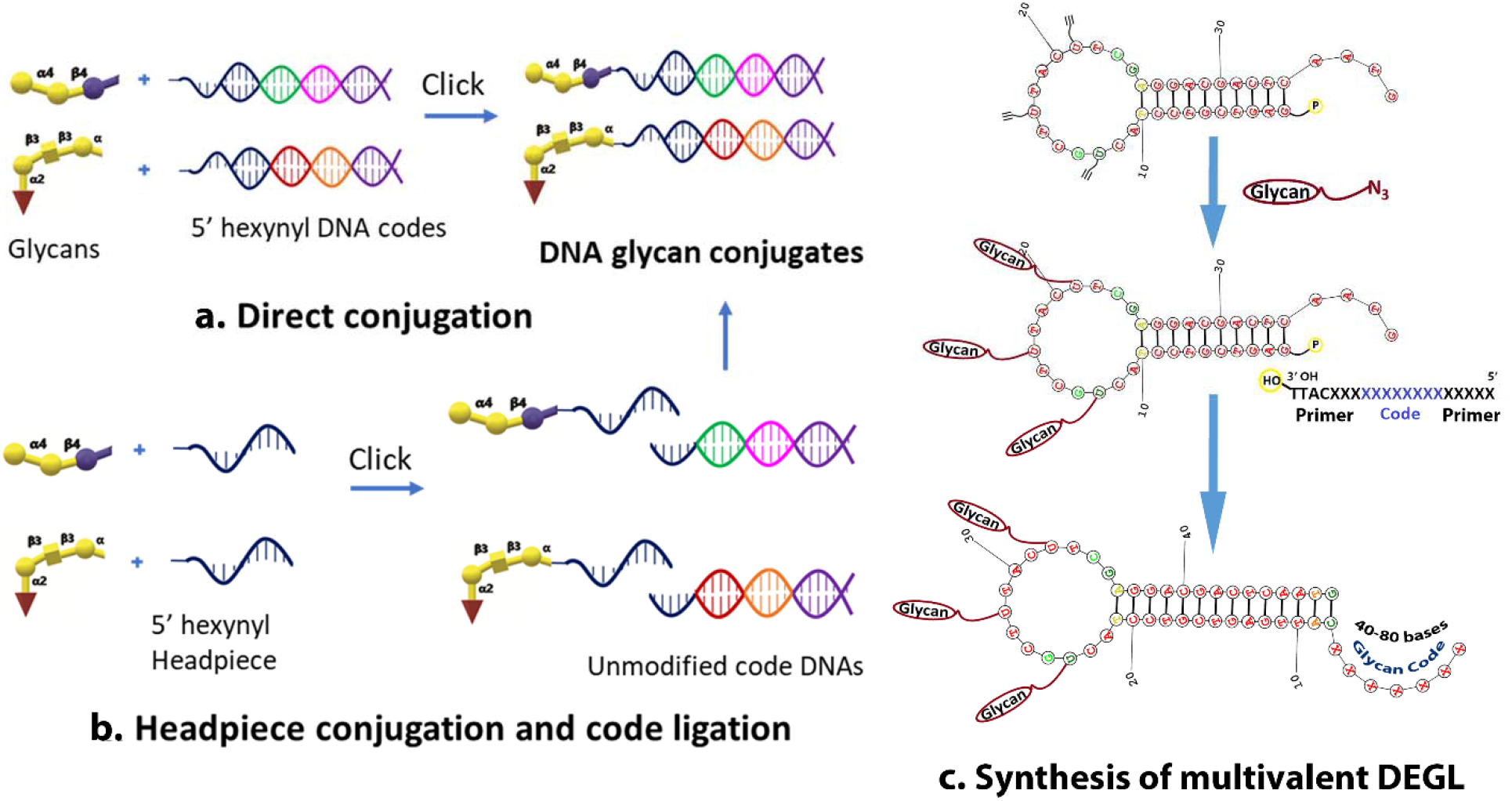
Different methods for synthesis of DNA encoded glycans. a) Glycans ‘click’ conjugated to the DNA sequences featuring DNA codes and primers for amplification. b, c) Glycans initially conjugated to the 5’ hexynyl headpiece or multi headpiece loop, then DNA codes are ligated enzymatically to the tail end of the headpiece DNA.

We report the use of an altered version of split and pool synthesis,[30] headpiece conjugation and code ligation (HCCL) approach to DNA-glycan production that is more adaptable and suited to high content DEGL (Figure 2b, 2c). HCCL is a two-step modular approach to synthesizing glycan conjugates. In the first step, a DNA headpiece is directly conjugated to the glycan using the approaches described above. In the second step, the desired DNA is ligated to the glycan via the headpiece. The HCCL method represents several advantages over direct DNA-glycan conjugation. HCCL does not require that each unique DNA sequence to be custom modified with a linker. Instead, only the headpiece needs to be chemically modified, meaning it can be obtained in bulk and applied to diverse glycans in DEGL. One terminus of the headpiece attaches to the glycan via click conjugation or NHS chemistry. The other terminus of the headpiece accepts the DNA coding fragment, which is introduced via routine enzymatic ligation and requires no special chemical modifications. HCCL can be modified by any unique DNA coding fragment, such as our IUPAC code or another the user prefers.

We also note the adaptable HCCL method provides opportunities to tune glycan presentation. Interactions of glycan-GBPs are usually weak in nature, and often they form multivalent interactions to result in tighter binding.[13, 31–34] By substituting the simple headpiece described above with a multivalent headpiece, both the number of glycans presented on each headpiece and the spatial display of glycans can be controlled (Figure 2c).

### Generation of headpiece glycan conjugates

We began by synthesizing a monovalent DNA headpiece (HP) that could be conjugated to glycans. HP consisted of the short single-stranded DNA ‘/5Hexynyl/AAT GAT ACG GCG A’. The hexynyl moiety of HP was directly conjugated to a sample of blood group A glycan (Figure 3) containing a propyl azido linker. Copper assisted azido alkyne (CuAAC) click conjugation proceeded with slight modification from previous reports.[19] Briefly, HPLC purified HP was mixed with different concentrations of glycan and click reagents, then allowed to react for two hours. Products were desalted using gel filtration columns (Clarion™ MINI Spin Columns), and the reaction was monitored using MALDI-TOF. Approximately ten equivalents of glycan were sufficient to entirely convert the alkyne moiety present in HP to corresponding glycan conjugate, as little to no HP was observed via MALDI TOF after the reaction. We successfully conjugated 2.5 nmol of HP with glycan, indicating the reaction was well suited for nanomole scale synthesis. It is possible to carry out the reaction at an even lower scale, but monitoring via MALDI-TOF becomes harder at low concentration, requiring further concentration steps before spotting MALDI plates.

**Figure 3.**
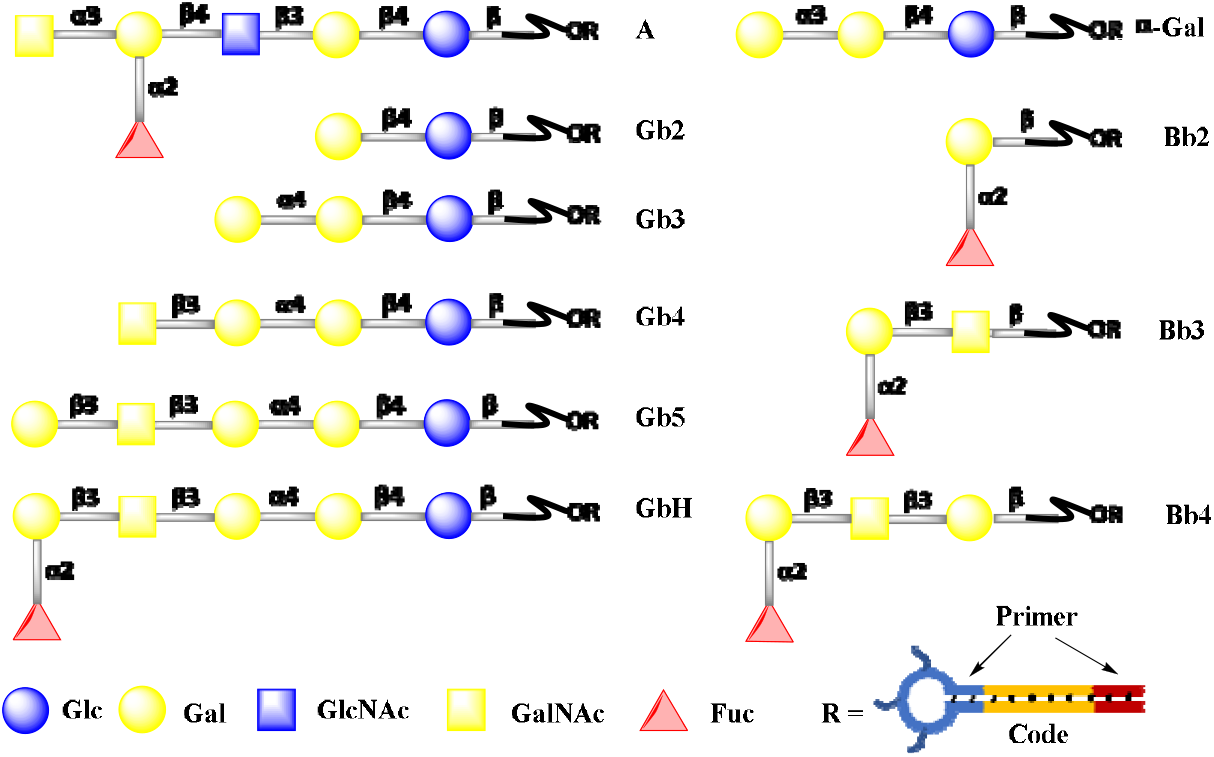
Glycans used in DEGL-10. Globo-series (GbH, Gb5, Gb4, Gb3, Gb2, Bb4, Bb3 and Bb2), blood group A glycan (A antigen) and α-Gal.

We also explored a remodeled headpiece that forms a looped DNA structure with three alkyne modifications (Octadiynyl dU) intended to attach three glycans to a single DNA code (Figure 2c). The multivalent DNA headpiece (MHP) consisted of ‘/5Phos/GAG TCG TCC TAC /i5OctdU/GC T/i5OctdU/T AC/i5OctdU/ TCG AGG ACG ACT CAA TG −3’. Conjugation proceeded as with HP. Headpiece conjugated glycans (Glycan+HP and Glycan+MHP) were quantified via nanodrop and stored for later encoding procedures.

### Generation of DNA-encoded glycans

Following the HCCL approach, we considered the unique DNA code to be ligated to the headpiece. During this step, it is possible to adopt different DNA codes based on instrument- or technique-specific requirements, such as NGS sequencing, qPCR, probe-based qPCR, or ligation PCR. In this study, we focused on creating a DEGL for NGS-based multiplex detection. Hence, the DNA possessed two distinct parts: primer binding sites and a unique coding region. The primer sites ensured a uniform condition for PCR amplification, while the unique coding region established the identity of each glycan.

Since HP and MHP were both conjugated with blood group A glycan, we procured the specific DNA codes for those glycans from the commercial source (Thermofisher, USA). DNA codes were ligated to the sticky ends of HP and MHP, respectively, using the T4 ligase (Figure 2b&2c, S1). Synthesis of monovalent conjugate was achieved by introducing the complimentary code strand, 5’-Primer-glycan code-TCGCCGTATCATT-3’ to form the dsDNA via hybridization to the headpiece, and subsequent ligation of the DNA code. For the multivalent conjugate, a tail DNA with the general sequence 5’-Primer-glycan code-primer-TTAC-OH-3’ was added, along with the rapid ligation buffer and T4 ligase to get the final DNA encoded multivalent glycan. DNA encoded glycan was then tested via PCR to confirm the amplification of the full-length code and to determine the optimum Tm value for later experiments (details are provided in Supporting material).

### Generation of DEGL

After optimizing the synthetic protocol above, we began synthesizing a first-generation DNA encoded library (DEGL) for the validation of selection and sequencing methods. We chose globo-series glycans, which exist as a part of glycosphingolipids, since their expression patterns are related to cancer metastasis and progression[35]. Structural features, binding specificities, and the clinical significance of these glycans are well studied, [35–37] thus providing a comparison for our new method. Globo H (GbH) and its truncated analogs Gb5, Gb4, Gb3 were synthesized from chemically synthesized Gb2 followed by enzymatic extension (Figure 3). To test the terminal epitope specificity, Bb4, Bb3 and Bb2 were also synthesized (Figure 3). All these glycans were then subjected to HCCL to obtain the corresponding DNA encoded glycans.

### Monovalent-displayed DEGL for qPCR (singleplex) and NGS (multiplex) detection

Initial validation was carried out using VK9, a mouse IgG monoclonal antibody which has reported specificity towards GbH and some of its truncated analogs. This validation step served as both a positive control for our synthesized reagents, and a baseline for binding that might be compared to our PCR based detection strategy. We used ELISA to test interactions between VK9 and our Globo-series glycans conjugated to BSA. Results confirmed binding of GbH and Bb4 to VK9, with comparatively lower binding towards Bb3 (Which is compared in Figure 7d. Partial binding of Bb3 may be attributed to the terminal epitope Fuc α1-2 Gal β1-3 GalNAc, which is recognized by VK9 antibodies.[38]

After confirming the specific binding of glycans with the target antibody, we tested the interactions of our GbH DNA conjugate (GbH+DNA) and VK9 using qPCR. After incubating GbH+DNA (25 pmol or 50 pmol) with the VK9 antibody, biotinylated anti-mouse secondary antibody was added to the mixture, followed by streptavidin magnetic beads. GbH+DNA-VK9-bead complexes were precipitated by magnet, and unbound DNA in the solution was eliminated by several wash cycles. Finally, bound DNA was eluted by heating the beads to 80°C for 10 minutes in 20 μL pure water and used as a template in qPCR assay. A control experiment omitting the VK9 antibody was used for comparison. As expected, the amplification and Ct values were correlated with the concentration of the antigens present in the solution. A low Ct value of 7 and 19 was observed for 50 pmol conjugates, and comparatively higher Ct value of 14 and 21 were obtained for 25 pmol conjugates. (Figure 4a).

**Figure 4.**
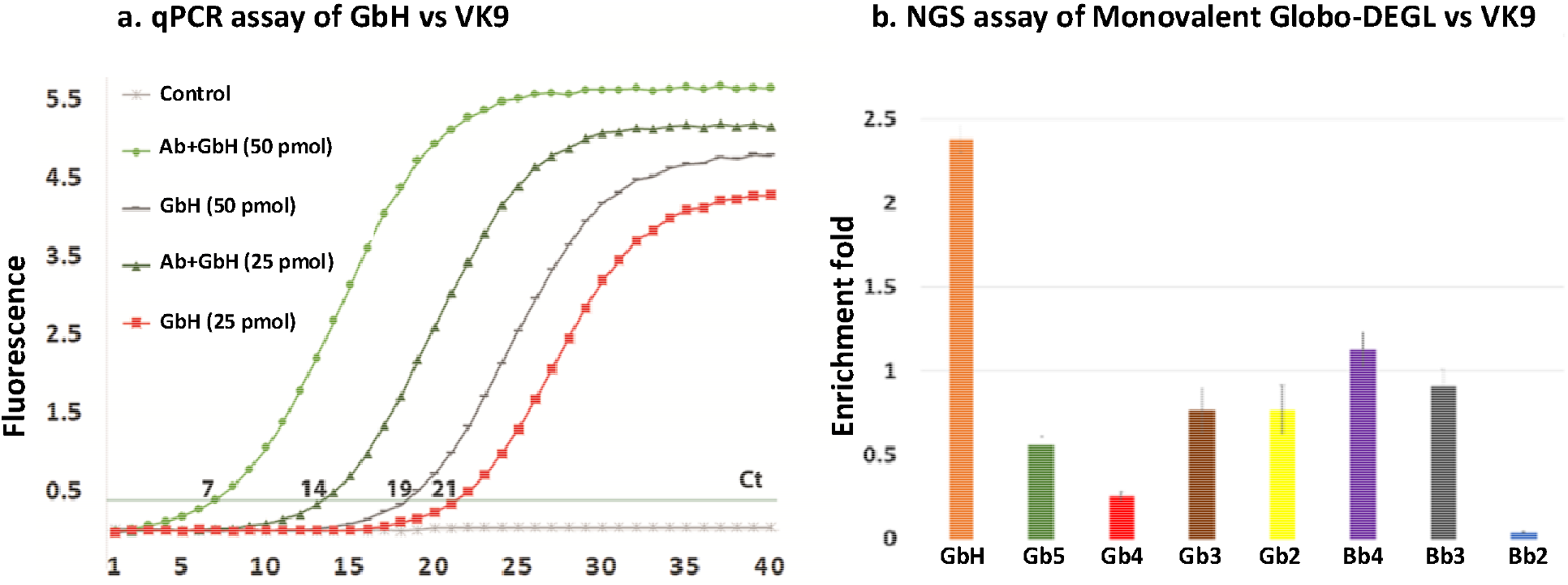
Singleplex qPCR assay of GbH glycan and a multiplex assay of globo series glycans against VK9 antibody. a) qPCR assay of GbH vs VK9: 50 pmol and 25 pmol of GbH+DNA conjugate incubated against VK9 antibody and Ct value with and without antibody are compared. b) NGS assay of Monovalent Globo DEGL vs VK9: 1 pmol of DEGL incubated with the VK9 antibody and the DNA selected were subjected to NGS. RPM count of each of the glycan codes was compared against the no antibody control selection. Data are the mean of two replicates. Error bars indicate ± standard error of the mean.

To demonstrate the multiplex assay using an NGS platform, we then pooled all the globo series glycans in an equimolar ratio to get the globo DEGL library. Varying quantity of the DEGL (50, 25, 10, 1, 0.1 pmol) were incubated with the VK9 antibody. The selection protocol was repeated as described above to get the DNA template for PCR (Figure 5). The NGS library was prepared in two steps. Initially, eluted DNA was amplified by PCR (18 cycles) followed by NGS barcoded fusion PCR (30 cycles) to incorporate the index codes and ion-torrent NGS adapters. NGS barcoded PCR products were recovered using AMPure beads, and the concentrations were measured using a bioanalyzer and qubit dsDNA assay kit. Sample concentrations were normalized and pooled to produce the NGS library, which was further diluted to 26 pM, and 25 μL was used in an ion-PGM semiconductor sequencer. Sequencing data were analyzed using an in-house developed program (uploaded in electronic supplementary information) for analysis of count and read of each sequence and each barcoded sample.[39] Read per million (RPM) sequence count of each DNA was then compared against the RPM count of the same DNA sequence in the no target control. When 1 pmol DEGL was used, GbH and its analog Bb4 were enriched compared to the remaining glycans, confirming the validity of the NGS method in detecting appropriate glycans from the mixture (Figure 4b). Although the NGS results matched the ELISA results, the enrichment folds (~2.4 for GbH and ~1.2 for Bb4) were not significantly high for the monovalent glycan-DNA conjugates, suggesting additional improvements could be made.

**Figure 5.**
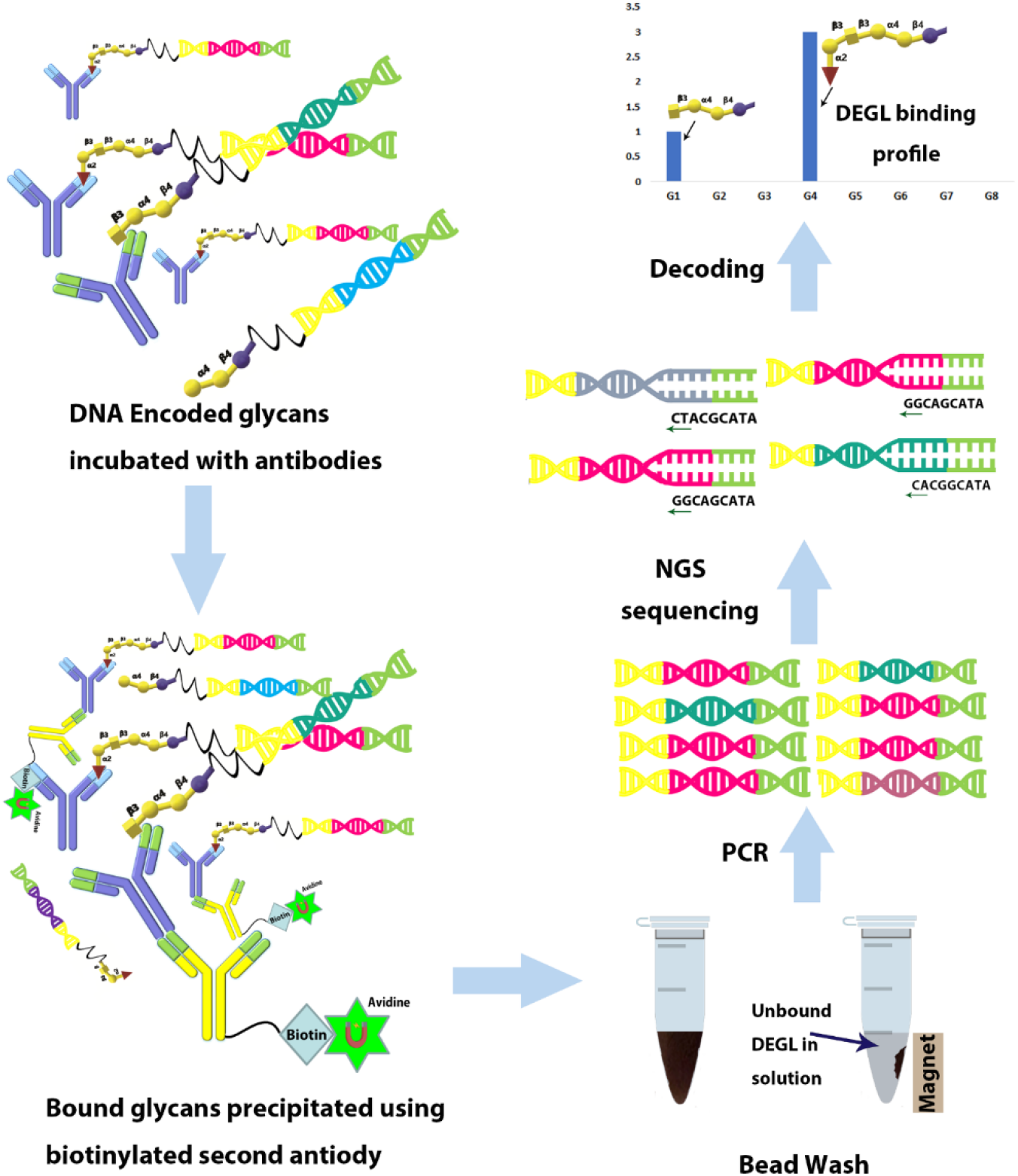
Protocol for the multiplex detection of glycan binding to the target. A DNA encoded glycan library (DEGL) was formulated by mixing equimolar concentrations of glycan-DNA conjugates. DEGL was then incubated with the target protein followed by the biotinylated secondary antibodies. The resulting complex was precipitated by magnet and unbound glycan conjugates were washed out in several cycles. Finally, the beads were eluted to 20 μL of pure water by heating to 80°C and used as a template in NGS library preparation. The data plotted is the ratio of the sequence count in RPM with and without target proteins.

### Multivalently-displayed DEGL-10 for qPCR (singleplex) and NGS (multiplex) detection

Glycans generally form multivalent interactions with target proteins for improved binding. To explore this route to improved binding in our conjugates, we synthesized a multivalent DEGL that contained 10 glycan structures (Figure 3), including the eight globo series glycans, blood A antigen and α-Gal. For cross comparison studies with VK9, we select two additional binding proteins.

The lectin *Helix pomatia* agglutinin (HPA) has specificity towards the terminal αGalNAc of A glycan [40–42], while *Erythrina Cristagalli* Lectin (ECA) has specificity towards the terminal lactose of Gb2.[43] Three multivalent glycan-DNA conjugates and their specific GBPs (GbH vs VK9, blood A antigen vs HPA, and Gb2 vs ECA) were subjected to singleplex selection with different quantity (10, 5, 1, 0.1 pmol of conjugates), followed by qPCR assay. A significant difference in Ct values between the DNA-glycan vs target and no target control were observed, even at a lower quantity (1 pmol) of DNA-glycans (Figure 6, S7).

**Figure 6.**
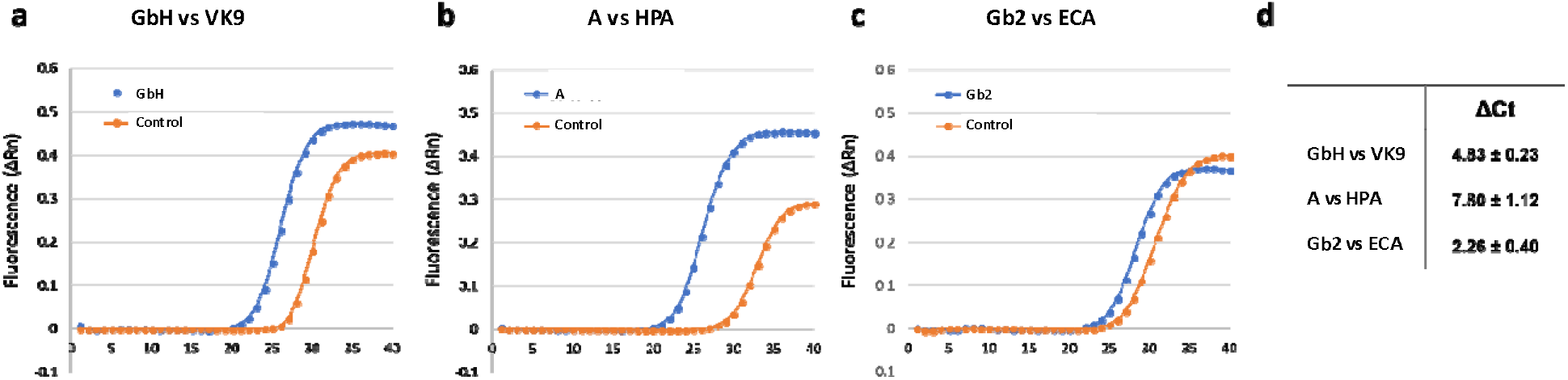
qPCR assay for singleplex selection of multivalent glycan conjugates (MHP+glycan). **a.** qPCR assay for MHP+GbH selection with VK9 antibody which gave the lower mean Ct value of 22.8 while control gave the mean Ct value of 27.7. **b.** qPCR assay for MHP+A vs HPA. Mean Ct value of A is 22.9 while control for A is 30.7. **c.** qPCR assay for MHP+Gb2 vs ECA. Mean Ct value of Gb2 is 25.5 while control for Gb2 is 27.8. **d.** Summary of the Ct value differences with and without glycan. Data are the average of two replicates ± SD.

**Figure 7.**
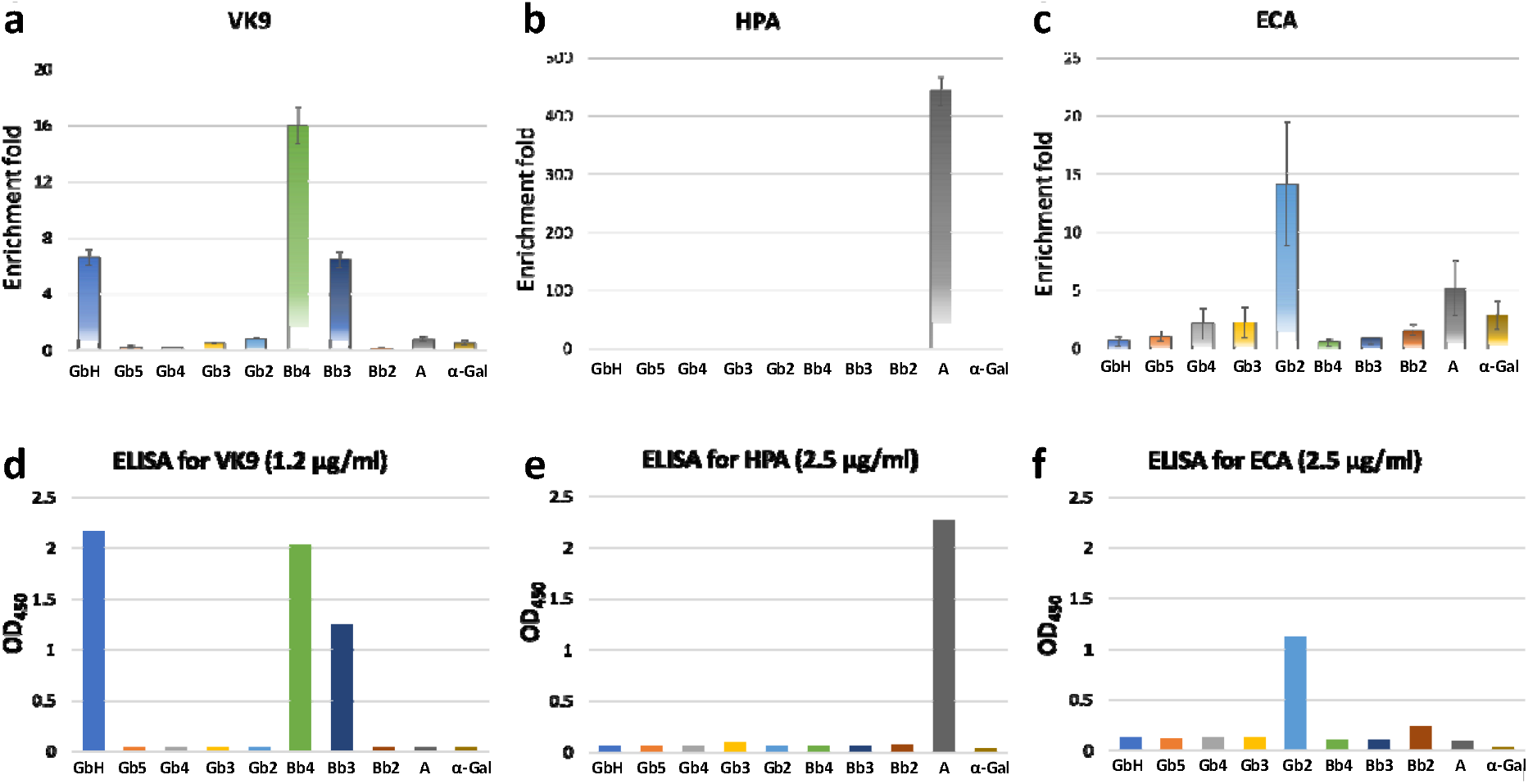
Multiplex assay of multivalent DEGL-10 against VK9, HPA and ECA (**a-c**), ELISA results of same glycans against the three GBPs (Glycan presented as BSA conjugates, **d-f**). NGS results of multivalent-DEGL towards the target proteins VK9, HPA and ECA showing enrichment over control with no GBP. VK9 showed enrichment of GbH (~7-fold), BB4 (~16-fold) and BB3 (~7-fold). HPA showed enrichment of blood group A antigen (~450-fold). ECA had good enrichment of Gb2 (~15-fold).

Following the singleplex study, we carried out a separate multiplex selection of multivalent DEGL for NGS assay. We assayed all three targets (VK9, HPA, ECA) in parallel, with triplicates of selection. 1 pmol of DEGL was incubated with the target proteins, and selection, washing, and elution were performed as described previously. Each selection was barcoded twice to obtain replicates.

Glycan binding was calculated using the read per million (RPM) of each glycan-code sequence in the DEGL with GBP and without GBP control. The result in Figure 7a-c shows the enrichment fold of each glycan in the library, and a binding pattern that was consistent with ELISA assays (Figure 7d-f). Compared to the NGS assay of monovalent glycans vs VK9 antibody shown in Figure 4b, enrichment fold difference for the GbH vs VK9 over GbH vs no antibody control was improved from ~2 fold to ~7 fold in this multivalent glycans. GbH, Bb4 and Bb3 glycans found to be binding to the VK9 antibody in both the ELISA and NGS, while Gb2 showed binding with the ECA lectin and A glycan showed a strong binding with the HPA lectin in both ELISA and NGS. HPA has been identified as a hexamer with a total molecular mass of 79 kDa[44] and six separate carbohydrate binding sites. Hence, it showed the greatest enrichment of blood group A antigen (more than 400-fold).

### High content and throughput assay using DEGL-50

To further validate the promising high throughput and high content aspect of the DEGL assay, we synthesized more than 50 glycans. In this DEGL-50, glycans included several important classes of glycans with biological significance, such as gangliosides, blood group glycans, and both 2-3 and 2-6 linked sialic acids. A full list of DEGL-50 glycans provided in the supporting information Figure S5. For the synthesis of DEGL-50, MHP was added to a 50-well plate and subjected to HCCL to generate the desired DNA-glycan conjugates (Figure S8, Glycan codes are listed in the SI table S1 and S3). Ligation was confirmed using gel electrophoresis. Concentrations of the Glycan+MHP-DNA conjugates were then normalized and pooled to make the final DEGL-50. Next, we chose a panel of target proteins that are known to bind glycans, including 20 plant lectins, VK9 (Mouse, IgG), anti-SSEA3 (Rat, IgM), anti-GM1 (Rabbit, IgG) and IgM-A (Mouse, IgM). We then probed each of the target proteins against DEGL-50. Selection and sequencing were repeated as described previously using 1 pmol of DEGL-50. Figure 8 shows representative results from four of the target proteins described in greater detail below.

**Figure 8.**
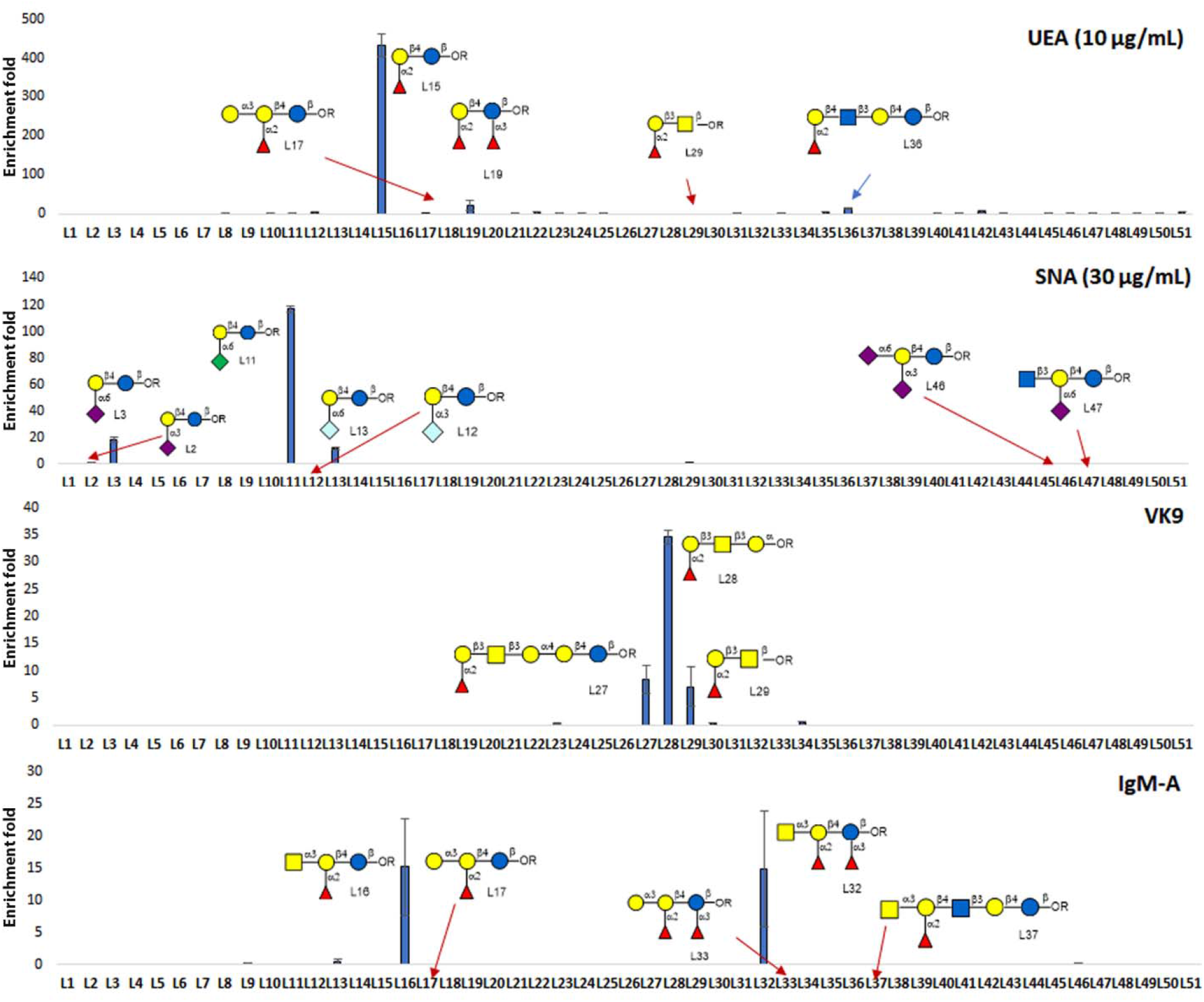
Representative examples for the high throughput-high content assay of DEGL-50. Binding profiles of two lectins (UEA and SNA), IgG antibody VK9, and IgM antibody IgM-A are provided. Each panel shows structures of glycans that are predicted to bind the GBP, as well as structurally similar glycans that were present in the library and did not show binding. Results shown are the mean of two replicates ± standard error of the mean. All experiments were repeated three times.

*Ulex europaeus* agglutinin (UEA) is reported to bind with blood group O cells, with preference for the blood group H type 2 (BGH2) and type 5 (BGH5) glycan motif featuring Fuc α1-2 Gal β1-4 GlcNAc or Fuc α1-2 Gal β1-4 Glc, respectively.[9, 45] As expected, UEA showed a higher signal for the L15 (BGH5) glycan, followed by L19. Some closely related structures which did not show any enrichment signal, like L17 and L29 also given in figures to depict the specificity of the binding. The signal intensity for L36, which has a similar motif Fuc α1-2 Gal β1-4 GlcNAc but in a long chain was also less, consistent with the reported literature.[9] Hence, the binding of UEA was specific to Fuc α1-2 Gal β1-4 Glc, and any addition to the core structure has reduced the affinity as indicated by the low intensity signal for L19, L36, and L49.

*Sambucus nigra* lectin (SNA), binds preferentially to sialic acid attached to terminal galactose in α2-6, and to a lesser degree, α2-3 linkages.[46] Using 30 μg/mL SNA lectin vs DEGL-50, we observed specific binding of the lectin to glycans bearing α2-6 linked sialic acid. A strong affinity was observed for Kdn α2-6 Gal (L11), with similar glycans Neu5Ac α2-6 Gal and Neu5Gc α2-6 Gal (L3 and L13, respectively) also showing affinity. Glycans with the α2-3 linkage (L2, L10, L12) did not show affinity in our assay, confirming the earlier reported preference of SNA lectin for α2-6 linkages.

In parallel, we repeated the selection of VK9 against DEGL-50. Results were consistent with the earlier assay of DEGL-10, even at the lower glycan levels used in the DEGL-50 assay. In the DEGL-10 experiment, the amount of each glycan was ~100 fmol in the final solution. In the DEGL-50 experiment, 50 different glycans, each at 1 pmol, were combined such that 20 fmol of each glycan was in the final solution. The result shows enrichment fold of 10, 35 and 8 for GbH, Bb4 and Bb3 glycans, respectively, compared to enrichment folds of 7, 16 and 7 showed earlier. This suggests that expansion of the library to higher contents may not have adverse effects on sensitivity.

The fourth GBP test we discuss was mouse IgM-A, representing the IgM class of antibodies in the assay. Mouse IgM-A showed higher enrichment (~16 fold) for the A-type glycans as seen for L16 and L32, with structural motifs GalNAc α1-3(Fuc α1-2)Gal β1-4 Glc β and GalNAc α1-3(Fuc α1-2)Gal β1-4(Fuc α1-3)Glc β, respectively. There was no binding observed for the blood group B glycans L17 and L33 which have Gal α1-3(Fuc α1-2)Gal β1-4 Glc β and Gal α1-3(Fuc α1-2)Gal β1-4(Fuc α1-3)Glc β motifs, respectively, suggesting the specificity of the detection method for the correct terminal GalNAc over Gal. We did not observe binding with blood group A glycan L37, GalNAc α1-3(Fuc α1-2)Gal β1-4 GlcNAc β1-3 Gal β1-4 Glc β, despite the presence of a terminal GalNAc, but a similar result was also reported while using MGBA method.[9]

Overall, 11 of 20 lectins tested (DBA, RCA, WGA, UEA, SNA, STL, VVL, Jacalin, ECA, HPA, RSA) showed binding consistent with prior studies. Binding intensity graphs are provided in the supporting information (Figure S9. Lectins that did not show any binding included SBA, ConA, MAL-I, PNA, LEL, DSL, GSL-II, CS-II, and GNA. Lack of binding to these lectins are likely correlated with the absence of lectin specific glycans in the library. For example, ideal epitopes recognized by lectins ConA, GNA, LEL, GSL-II, and DSL were not included in DEGL-50. Alternatively, the lack of signal from the assay of several lectins (SBA, MAL-II, PNA, CS-II) and antibodies (SSEA3 and GM1) may be due to unoptimized buffer conditions and resulting low affinity interactions between target proteins and binding partners. A similar diversity of results was observed in a related MGBA assay as well.[9] Results may be improved if higher concentrations of lectins were used. For instance, SNA lectin did not show significant binding when used at a concentration of 10 μg/mL, but at 30 μg/mL, the signal was much improved without compromising the specificity of the binding (Figure 8).

## DISCUSSION

Detection of the interactions between glycans and GBPs is critically important for biomedical and clinical research.[4, 9, 11, 47–51] Recognizing the importance of glycans, research has focused on synthesis and isolation of glycans from different sources, including animals and plants.[12, 27, 52–54] Synthetic efforts have led to the availability of a large number of glycan structures for study, and the number is increasing quickly. Despite these advances, high throughput methods to investigate the functions of diverse glycan structures have lagged behind. Here we report a DNA encoded glycan library (DEGL) platform that can address these challenges. DEGL uses customizable DNA barcodes to identify unique glycan structures. The methods utilize wide spread molecular biology technologies like qPCR and NGS for signal amplification and reading of unique glycan codes. Since DEGL relies on PCR, qPCR and NGS, it offers an affordable and accessible method for binding studies of glycans. Advantages over existing methods include multiplex analysis, high sensitivity, low cost and flexibility to tune the glycan presentation.

Advancements in DNA synthesis and sequencing have extended beyond genetics, and applications in glycoscience are a logical next step. DNA encoded libraries (DEL) of small molecules [30] in drug screening and lead optimization have already been applied to research in major pharmaceutical companies.[14] In this study, we applied DEL technologies to glycans, which are restricted by low sensitivity detection techniques. This technology draws upon an almost limitless availability of DNA barcodes for both the encoding of the glycans and the index barcoding of samples. With virtually unlimited DNA barcodes, we can finally imagine all the synthetic and natural glycan epitopes available for study in a single tube. However, the synthesis of DEGL is not as straight forward as the DEL of small molecules. DEL synthesis is achieved by careful selection of the DNA compatible building blocks and DNA enabling chemistries for the coupling.[30] Realizing the difficulties in attaining broad substrate scope for DNA enabled chemistry, we developed a modular HCCL approach as an alternative. This approach also requires careful design and optimization of headpiece DNA and glycan chemical coupling procedures, but removes the need for custom chemical modeling of each unique DNA code.

Competing methods for glycan analysis lack uniformity, even within a similar platform of technology, thereby making it difficult for researchers to compare results from different labs.[13] For this reason, we believe developing a simple general protocol for DEGL is very important. A major challenge to achieve uniformity is in the selection of codes. Our structure-based DNA code is a way to introduce a systematic method for the coding of glycans. The program works based on a list of different structural motifs of glycans, and can be updated as new natural and synthetic motifs are realized. We have generated DNA codes for many existing glycan structures, and in general, the base length of most of them fit a bell curve spanning approximately 40-120 bases (Supporting information Table S8). Following a general protocol, we can maintain this approximate range by adding additional sequences for small DNA codes, or by introducing abbreviated core structure sequences for overly long DNA codes. Additional modifications can be made within the coding program if a different base pair range, or an identical base pair length, is desired for PCR amplification. The proposed modular library design and modular HCCL strategy are well suited for those cases.

Copper assisted azide-alkyne cyclo condensation (CuAAC), or ‘click’ chemistry, is one of the most used attachment linkers in DNA conjugation reactions.[55] A direct approach for DNA encoding attaches the whole amplifiable DNA code to the glycan in a single click reaction. Although direct conjugation is the most straightforward approach, it also has some limitations. Despite the growing availability of DNA sequences on demand, the direct conjugation method is better suited for custom synthesis and low content DEGLs. Incorporating a chemical attachment tag into a large library of DNA molecules is not straightforward, as yields are relatively low, products often must be HPLC purified before NGS, and commercial alternatives may not be available from on demand DNA suppliers. These factors contribute to escalated prices for modified DNA, and are therefore cost-prohibitive for direct conjugation at the scale of 100-1000 glycans.

Alternatively, DNA without any attachment chemistry (5’, 3’ or internal) has many advantages. Naked DNA is low in cost, HPLC purification is not necessary, and nanomole scale synthesis is feasible. Hence, our modular HCCL approach draws upon existing commercial infrastructure for DNA synthesis and lower cost reagents. In HCCL, a single headpiece is used for all the glycans, then DNA code (unique for each glycan) is ligated by T4 ligase enzyme, thus requiring no chemical modification to either end of the DNA code. A small headpiece allows efficient coupling in water, which reduces purification steps and costs. Furthermore, the reactions can be tailored to the picomole scale to minimize consumption of the headpiece DNA and precious glycan materials. Ultimately, HCCL has the potential to remove cost barriers to the construction of comprehensive libraries for glycan analyses.

In this study, we also examined functional advantages of the modular HCCL method for using different headpiece DNAs. By careful design, it is possible to investigate multivalent presentation, the spatial orientation of glycans, and attachment chemistries. We demonstrated the synthesis of monovalent and trivalent glycan conjugates with two different headpieces. A 5’-modified headpiece was used in first generation library, followed by a loop forming DNA with alkyne modified internal bases in subsequent versions. Internal bases are fine-tunable for number, and spatial distribution of glycans, suggesting that additional studies could examine a diversity of glycan presentations.

In our hands, the effect of glycan presentation was most evident in the case of comparatively large glycans. We observed a general trend of reduced affinity as the glycan chain length increased. For instance, L16, L32, and L37 all have blood group A terminal epitope, but only L16 and L32 glycans showed binding to the IgM-A antibody. Similarly, in monovalent DEGL, GbH showed the highest affinity towards the VK9 antibody, while the truncated analogs Bb4 and Bb3 showed a moderate binding. But a different affinity pattern was observed with multivalent DEGL. In this case, Bb4 showed the highest affinity, indicating possible steric interactions, especially for the longer polysaccharides. More work needs to be done to understand the multivalency fully and to identify the best possible headpiece. Many details, including changes in the secondary structure of DNA when glycans are conjugated, may need to be considered before creating an even higher content DEGL.

Using our current DEGL, we detected glycan-target interactions in solution that required minimal amounts of sample and were comparable with ELISA assays. For instance, the ELISA procedure required 10 pmol of BSA-glycan for the experiment, while the DEGL procedure only required 20 fmol of each glycan (1 pmol of DEGL), resulting in a 500-fold reduction in sample consumption. Even at this low amount of sample, the enrichment signal from the DEGL method was as high as ~400 fold for the hexameric HPA lectin. While not every target contains six binding sites, we observed moderate binding and enrichment for ECA and other targets as well. Our results indicate that binding signal (enrichment fold) can be varied according to the glycans and lectins used in the study. This suggests the general method we have proposed can be further fine-tuned for specific glycan-GBP interactions. Still, our overall results indicate that the signals of glycan-GBP interactions are geometrically enhanced by multiple grade amplification involved in the NGS method.

Comparing our results from the DEGL-10 and DEGL-50 trials suggests that even larger libraries are viable. We note that the method did not lose its sensitivity when more glycans were added to the library. Compared to the low content library, sensitivity was actually higher in the high content library analyses. Therefore, we plan to rapidly expand the library size by 10-to 20-fold. Beyond our own work, DEGL from multiple research groups with diverse glycan sources, content, and intended applications could be pooled into a unified library to be assayed in a single NGS run, which would accelerate the cost-effectiveness and robustness of the method. In this study, we processed 72 samples in a single assay using 36 DNA barcodes, and each barcode received ~80,000 reads (ion 520 chip, 3-6 million reads). When bigger chips are used to their full capacity, 5 million reads of NGS can be currently obtained for $200 [56], driving down costs. Moreover, since our method uses a modular approach for the DEGL, library synthesis can be remodeled to fit any current and future gene sequencers. We believe that DEGL technology can fulfill the critical need for high content and high throughput studies of glycan interactions, which will help to advance the glycobiology field.

## METHODS

### Synthesis of glycan DNA conjugates

All DEGL-50 glycans were prepared by chemoenzymatic synthesis.[57–61] All oligosaccharides contained an azido propyl linker at the reducing end for conjugation with DNA via click chemistry. DEGL-50 contained important glycan epitopes, including α2-3/6-sialic acid epitopes, Blood ABO antigen epitopes, globo series glycans, and many ganglioside oligosaccharides. Glycan (Antigen)-DNA Conjugates were synthesized using Azido-Alkyne cycloaddition click reactions. The 5’-/5Hexynyl-terminated DNA was procured from the commercial suppliers (IDT and Thermofisher). Desalted DNA was HPLC purified before click conjugation.

### Headpiece-glycan click reaction procedure

Conjugation was conducted based on previous reports.[19] Briefly, water soluble THPTA (10 mM), CuSO_4_ (10 mM) and Ascorbic acid (250 mM) were prepared. CuSO_4_ and THPTA were premixed in a 1:1 ratio. DNA (10 μL, 500 μM), Azido sugar (10 μL, 5 mM), and THPTA:CuSO_4_ (5 μL, 1:1 ratio) were combined. 24 μL of H_2_O was added, followed by 1 μL of ascorbic acid. Reactions were kept at room temperature for 2 hrs. 12.5 μL of THPTA was added to the mixture to quench the reaction. Reactions were desalted with Clarion™ N25 Columns according to the manufacturer’s protocol. DNA glycan conjugates were analyzed by nanodrop, HPLC and MALDI-TOF.

### Analytical HPLC method

Agilent 2100, Eclipse Plus C18, 3.5 μM, 4.6 × 100 mm column; Solvent A: pH 7.0 TEAA buffer 0.1 M, Solvent B: 20% ACN in solvent A; 5% B to 95% B over 30 minutes, 30-40 minutes 95% B with a flow rate of 0.6 mL/minute.

### MALDI-TOF MS analysis

For matrix preparation, 10 mg of 3-Hydroxypicolinic Acid (HPA) was dissolved in 200 μL of 50% ACN in water, and separately 10 mg of dibasic ammonium citrate (DAC) was dissolved in 200 μL of H_2_O. HPA and DAC were mixed 8:1 for the final matrix solution. Matrix solution (1 μL) was spotted to the MALDI target plate and allowed to air dry. DNA glycan conjugate (1 μL) was added to the dried matrix, allowed to completely dry, and analyzed using a Bruker MALDI-TOF instrument.

### DNA code ligation

Glycan-HP conjugates (10 μL, 20 μM) were mixed with the reverse complement DNA (2 μL, 100 μM) and 5’-phosphorylated tail DNA (2 μL, 100 μM). The mixture was heated to 70°C for 5 min and allowed to cool to room temperature. 5X rapid ligation buffer (3.75 μL) and T4 ligase (1 μL) were added, and the mixture was kept at room temperature for 30 minutes. Then, T4 ligase was denatured by heating the mixture at 70°C for 10 minutes. After cooling the DNA-HP-Code was directly used in further studies. (Rapid DNA Ligation Kit, Thermo Scientific™, Catalog number: K1422).

### Immunoprecipitation of GBP-Glycan-DNA Conjugates with Streptavidin Magnetic Beads

10 μL of glycan(antigen)-DNA Conjugate was incubated with 1 μL (50 μg/mL) VK9 antibody (Globo H Monoclonal Antibody, eBioscience; Catalog Number: 14-9700-82) for 2 hr at room temperature. After 2 hr, 0.5 μL of Biotin Conjugated Goat Anti-Mouse IgG Secondary Antibody (Thermofischer Scientific; Catalog Number:31800) and 90 μL of TBST buffer (0.1 % Tween 20, 0.1% BSA) was added and incubated at room temperature for 1 hr. The same protocol without the secondary antibody was applied for biotinylated lectins. Incubation without a target antibody or lectin was performed as a control. After 1 hr the solution was incubated with 15 μL prewashed Streptavidin Magnetic beads (Dynabeads Myone Streptavidin T1; Catalog Number: 65601) for 30 mins at room temperature. Then beads were washed 7 times using 100 μL TBST buffer and 2 times using pure water. The washed beads were then diluted in 20 μL of pure water and eluted at 80°C for 10 minutes. 1 μL of eluted samples were used for PCR, qPCR, or NGS detection.

### NGS Library Preparation

1 μL of the eluted sample was amplified using universal template primers with Taq polymerase for 18 cycles. PCR products were purified using Agencourt AMPure XP magnetic beads (30% PEG) as per manufacturer instructions. Purified PCR products were amplified again using NGS adapters (30 Cycles) and re-purified using Agencourt AMPure XP magnetic beads. Purified product concentrations were measured using an Agilent 2100 Bioanalyzer/Qubit dsDNA HS assay kit. An equimolar solution of 26 pM (100 pM for ion S5) was prepared, and 25 μL was loaded in ion Chef for Chip preparation and sequencing by Ion PGM or Ion S5.

### Data analysis

Sequencing data were converted into FASTAQ files for each of the barcodes employed. We used an inhouse program to count the DNA code sequence present in each of the files (code uploaded as electronic supporting information). Read count was normalized to read per million (RPM) taking account of the number of reads for each index barcode. Enrichment fold was then calculated by diving the RPM count of each glycan in the sample with the average count of two no target control experiment.

## Supporting information

GLYCODER

Raw data

Raw data

## Acknowledgments

This work was partially supported by Georgia Research Alliance, Atlanta, GA. We thank Mrs. Ping Jiang and the GSU core facility for sequencing services.

## Author Contribution

S.M.K. designed, synthesized, characterized and assayed DEGL-50; L.W. synthesized the glycans; H.Z. developed selection and sequencing methods, conducted ELISA assay; A.P. validated primers, PCR and qPCR assays, and wrote the coding program; S.P., A.P., and J.S. wrote and webhosted of the programs. P.G.W conceived the study, coordinated research and edited the manuscript; All authors contributed in manuscript preparation.

## Material request & Correspondence

Should be addressed to P.G.W.

## Computer program and Data availability

The computer programs used for encoding glycans or counting sequences, and raw data generated and analyzed during the study are available as a part of electronic supporting information. Further queries can be directed to the corresponding author.

