## Supplemental Info for "DNA Encoded Glycan Libraries as a next-generation tool for the study of glycan-protein interactions"

| **Materials and methods** | **Page 3-11** |
| --- | --- |
| **Figure S1-S9** | **Page 12-20** |
| **MALDI TOF MS and HPLC results** | **Page 21-53** |
| **Table S1. DNA sequences used in HCCL library synthesis** | **Page 53-55** |
| **Table S2. MALDI Mass of DNA glycan conjugates** | **Page 55** |
| **Table S3: DNA Sequences used for the synthesis of DEGL-50** | **Page 56-61** |
| **Table S4-S6. Coding Dictionaries (Library A, B, C)** | **Page 62-67** |
| **Table S7. Core structure and their codes (Library D)** | **Page 68-70** |
| **Table S8. Examples of DNA encoding of glycans** | **Page 71-97** |
| **References** | **Page 98** |

### 1. Materials and Methods

All chemicals and biological reagents were purchased from Thermo Fisher unless otherwise mentioned. Maxima SYBR Green/ROX qPCR Master Mix, Platinum Pfx DNA Polymerase were purchased from Life Technologies (Carlsbad, CA). Tris[(1-Benzyl-1H-1,2,3-Triazol-4-yl) methyl] amine (TBTA) Click chemistry Ligand were purchased from TCl (Tokyo,Japan). MicroAmp 96 well Fast PCR Reaction Plate, MicroAmp Optical Adhesive Film, MicroAmp Fast Reaction Tubes, Strips were purchased from Applied Biosystems (Foster City, CA). Primers and headpiece oligonucleotides were purchased from Integrated DNA Technologies (Coralville, IA), DNA codes, NGS primers and adaptors were purchased from Invitrogen (Carlsbad, California).

Primary antibodies used are mouse anti-Globo H monoclonal antibody VK-9 (IgG; eBioscience™, Catalog #: 14-9700-82), Blood Group A Antigen Monoclonal Antibody (Mouse / IgM, HE-193, Thermo fisher scientific, Catalog # MA1-19693), SSEA3 Monoclonal Antibody (Rat / IgM, MC-631, Thermo fisher scientific, Catalog # MA1- 020) and Anti-Ganglioside GM1 antibody (IgG, Rabbit polyclonal to Ganglioside GM1, Abcam, ab23943). The secondary antibodies used were Biotin-conjugated Goat anti-Mouse IgG (H+L) (Invitrogen, Catalog # 31800), Goat anti-Mouse IgM Secondary Antibody, Biotin (Invitrogen, Catalog # 31804), Goat anti-Rat IgM Secondary Antibody, Biotin (Invitrogen, Catalog # 31832) and Goat anti-Rabbit IgG (H+L) Secondary Antibody, Biotin (Invitrogen, Catalog # 65-6140). Biotin conjugated lectins were purchased from vector labs (Burlingame, CA). Taq DNA Polymerase, Maxima SYBR Green/ ROX qPCR Master Mix (ThermoScientific, Catalog #: 10342020, Catalog #:  K0221). HRP conjugated streptavidin was purchased from Abcam (Catalog #: ab7403). Beads used were streptavidin conjugated magnetic beads (Dynabeads™ MyOne™ Streptavidin T1, Invitrogen, Catalog #: 65601), Agencourt AMPure Beads (Beckman Coulter). Qubit™ 1X dsDNA HS Assay Kit (Invitrogen, Catalog #: Q33230) and Agilent DNA 1000 Kit (Part no: 5067-1504, Agilent, Santa Clara, CA) were used for DNA quantitation in NGS application. Clarion^TM^ MINI Spin Columns, DesaltS-25N, was purchased from Sorbent Technologies.

**Synthesis of glycan DNA conjugates**

A glycan library (Figure S5) that contain 50 oligosaccharides was prepared by chemoenzymatic synthesis.[1-5] All oligosaccharides have an azido propyl linker at the reducing end for next conjugation step with DNA via click chemistry reaction. This glycan library covers most of important glycan epitopes, including a2-3/6-sialic acid epitopes, Blood ABO antigen epitopes, globo series glycans, and many of Ganglioside oligosaccharides. Glycan (Antigen)- DNA Conjugates were synthesized using Azido-Alkyne cycloaddition click reaction. The 5’-/5Hexynyl-terminated DNA was procured from the commercial suppliers (IDT) with standard desalting purified. Desalted DNA was HPLC purified before click conjugation.

**Click reaction procedure**

Headpiece-glycan conjugation: We chose water soluble THPTA ligand considering the following T4 ligation. Click procedure was adopted from a reported method with little modification.[6] Briefly, THPTA (10 mM), CuSO_4_ (10 mM) and Ascorbic acid (250 mM) were prepared. CuSO_4_ and THPTA is premixed in a 1:1 ratio. 10 µL of DNA (500 µM), Azido sugar (10 µL, 5 mM) were added to the 5 µL of THPTA: CuSO_4_ in the tube. 24 µL of H_2_O was added followed by 1 µL of ascorbic acid. Reaction were kept at room temperature for 2 hrs. 12.5 µL of THPTA were added to the mixture to quench the reaction. Next, the mixture is desalted using the Clarion™ N25 Columns according to the manufacturer’s protocol. Concentrations of the DNA glycan conjugates were checked by nanodrop and analyzed by HPLC and MALDI-TOF.

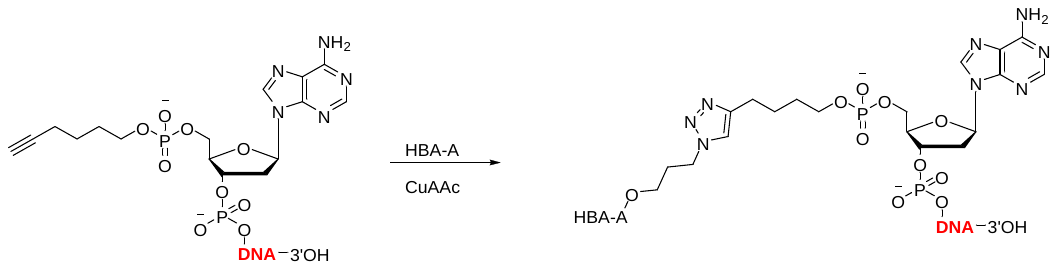

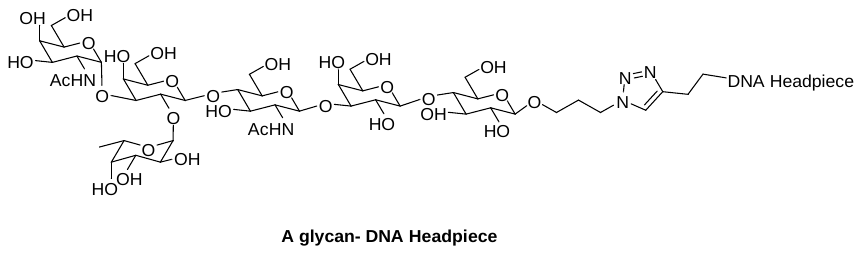

**Scheme S1.** Click conjugation of 5’-hexynyl DNA and azido modified glycans.

CuSO_4_ and THPTA is premixed in a 1:1 ratio. 10 µL of DNA (500 µM), Azido sugar (10 µL, 5 mM) were added to the 5 µL of THPTA: CuSO_4_ in the tube. 24 µL of H_2_O was added followed by 1 µL of ascorbic acid. Reaction were kept at room temperature for 2 hrs. 12.5 µL of THPTA were added to the mixture to quench the reaction. Next, the mixture is desalted using the Clarion™ N25 Columns according to the manufacturer’s protocol. Concentrations of the DNA glycan conjugates were checked by nanodrop and analyzed by HPLC and MALDI-TOF.

**Analytical HPLC method**

Agilent 2100, Eclipse Plus C18, 3.5 µM, 4.6 X 100 mm column; Solvent A: pH 7.0 TEAA buffer 0.1 M, Solvent B: 20% ACN in solvent A; 5% B to 95% B over 30 minutes, 30-40 minutes 95% B with a flow rate of 0.6 mL/minute.

**MALDI-TOF MS analysis**

Matrix preparation: 10 mg **3**-Hydroxypicolinic Acid (HPA) was dissolved in 200 µL of 50% can in water; in another tube, 10 mg of dibasic ammonium citrate (DAC) was dissolved in 200 µL of H_2_O. Then mixed HPA to DAC in 8:1 in a new tube for the final working matrix. Next, 1 µL of matrix solution was spotted to the MALDI target plate and allowed to air dry, on top of the dried matrix 1 µL of HPLC purified DNA glycan conjugates were spotted, allowed to complete dryness and analyzed using Bruker MALDI-TOF instrument.

**DNA code ligation**

10 µL of 20 µM glycan-HP conjugates were mixed with the reverse complement DNA (2 µL, 100 µM) and 5’- phosphorylated tail DNA (2 µL, 100 µM). Mixture was heated to 70^o^C for 5 mins and allowed to cool to room temperature. 3.75 µL of 5X rapid ligation buffer and 1 µL of T4 ligase was added and the mixture is kept at room temperature for 30 minutes. Then, T4 ligase was denatured by heating the mixture at 70^o^C for 10 minutes. After cooling the DNA-HP-Code was directly used in further studies. (Rapid DNA Ligation Kit, Thermo Scientific™, Catalog number: K1422).

**Gradient PCR and Tm Determination**

Initially a gradient PCR was performed to figure out the optimal melting temperature to be used in the further quantitation experiments. Platinum Pfx DNA Polymerase was used for the PCR, approximately 26 ng of template, 1 µL of primers (10 µM), 1mM dNTP, 10X amplifying buffer, MgSO_4_, Polymerase and nuclease free water was used for a typical 25 µL reaction. Standard thermocycling condition were used with different temperature in the melting phase (95°C for 10 min, X°C for 30 s, 95°C for 15 s, 13 cycles) wherein X represents the melting phase and temperature gradient from 50^o^C - 60^o^C were tested during this phase. The amplified PCR products were analyzed using 4% agarose gel electrophoresis. T_m_ determined using this method was used for qPCR.

| Component | 25 µl | 50 µl | Final Concentration |
| --- | --- | --- | --- |
| 10X Amplifying Buffer | 5 | 10 | - |
| MgSO_4_ | 1 | 2 | - |
| dNTP (10 mM) | 0.75 | 1.5 | 0.3 mM |
| Forward Primer (10 µM) | 0.75 | 1.5 | 0.3 µM |
| Reverse Primer (10 µM) | 0.75 | 1.5 | 0.3 µM |
| Template (1 µM) | 1 | 2 | 0.04 |
| Polymerase | 0.3 | 0.5 | - |
| Nuclease Free Water | 15.4 | 31 | - |

**PCR, qPCR validation of DNA glycan conjugates and limit of detection analysis**

The success of the DEGL lies in the utility of applying the DNA codes to the standard PCR and qPCR protocols. It is crucial to have the glycan-coupled DNA (G+DNA) achieve similar PCR efficiency to the native DNA, so we compared the G+DNA conjugates with the corresponding native DNA strands to verify the efficiency of amplification in both PCR and qPCR. PCR was performed on DNA and G+DNA templates using standard thermal cycles with varying annealing temperatures. Gel electrophoresis analysis implied both DNA and G+DNA amplified well in the range of 52-60ºC (Figure S1). Next, we examined the qPCR detection limit of the G+DNA conjugates; both DNA and G+DNA conjugates were serially diluted to provide different concentrations of templates, with final concentrations ranging from 2.4 nM to 16 pM. Standard curve qPCR was carried out to obtain the corresponding critical threshold value (Ct), which is used to compare the concentration of the templates in the qPCR reaction mixture, a low Ct indicating high template concentration. (Ct or threshold cycle is a measurement of signal intensity for qPCR experiments. In a qPCR experiment, PCR is performed in presence of a fluorogenic intercalating dye (SYBR Green in our case). The dye intercalates with double-stranded DNA, as more double-stranded DNA is produced in each PCR cycle the dye increases the fluorescence intensity. Once the fluorescence intensity reaches a threshold level, the cycle number is recorded by the instrument as Ct value. Therefore, sample having large amount of DNA will have a lower Ct value compared to those samples containing relatively less amount of DNA.) Ct values of the conjugates from 2.4 nM to 16 pM concentrations were observed in the range of 5-25, while the negative control (no template control, NTC) gave a Ct value above 30 (Figure S3a). A standard plot of the Ct value *vs* log concentration of the DNA and G+DNA conjugate fit well within a linear relationship indicating the successful application of G+DNA for quantitative detection up to the *pico* molar level (Figure S3b).

**General qPCR protocol for a typical 20 µL reaction**

1 µL DNA-glycan (Antigen) Conjugates were added to 10 µL 2X Maxima SYBR Green/ ROX qPCR Master Mix with 1 µL primers (Forward Primer – 2 µM and Reverse Primer- 2 µM). qPCR was performed with Applied Biosystems Stepone System (50^o^C for 2 mins (Holding), 95^o^C for 10 mins (Holding), 95^o^C for 15 s, 60^o^C for 30 s ,72^o^C for 30 s for 40 cycles).

| Component | Volume (µL) |
| --- | --- |
| 2x Maxima SYBR Green/ROX qPCR master mix | 10 |
| Forward Primer (2 µM) | 1 |
| Reverse Primer (2 µM) | 1 |
| Template (Gradient as stated above) | 1 |
| Nuclease Free Water | 7 |

**Standard curve plot of DNA and G+DNA**

To prove the consistency over concentration for both pure DNA and G+DNA, a qPCR standard curve reaction was conducted over a series of 8 different concentrations (2.4 nM, 1.2 nM, 600 nM, 260 nM, 130 nM, 64 pM, 32 pM, and 16 pM) with standard thermocycling conditions stated above.

**Synthesis of Multivalent glycan DNA conjugates**

Synthesis of headpiece glycan conjugate: The same click protocol described for the monovalent headpiece glycan conjugate was employed with a higher concentration of glycan azide (10 µL of 20 mM glycans). DNA code ligation: The multi headpiece loop has a 5’-phosphorylated end and a sticky 3’ end for the hybridization of the reverse compliment of the DNA code. Hence, 10 µL of MHP-Glycan (20 µM) were mixed with 2 µL of reverse DNA and allowed to hybridize and similar ligation procedure described above was used. Schematic representation of the total protocol is provided in Figure S8.

**Indirect ELISA experiment**

Since sugars have low binding affinity for unmodified plastic surfaces, the coating antigens, BSA-glycans, were conjugated by a linker called Propargyl-N-hydroxysuccinimidyl ester (purchased from Sigma-Aldrich) via NHS reaction and then click reaction (Figure S6). The BSA-glycans were characterized by MALDI-TOF analysis.

The 96-well plate (Costar Polystyrene High Binding Plate 3590) was coated with 100 mL of 8 µg/mL BSA-glycans in 0.01 M PBS (pH 7.4) at 4oC overnight. The coated plate was then washed with 150 µL of 0.05% Tween-20/PBS buffer (pH 7.4) (PBST) for three times and then blocked with 2% (w/v) BSA in PBST for 2 hrs. After washed with another 150 µL PBST for 3 times, 100 µL Biotinylated lectins with a series of dilution were added. After 2 hrs incubation at room temperature, then the plate was washed with 200 µL PBST for 6 times, and HRP-conjugated-streptavidin (Abcam) was added to the plate for 1 hr at room temperature. For detection of antibodies, 100 µL antibody with a series of dilution was added, followed by adding 100 µL of diluted biotinylated-secondary antibody. After washed with another 200 µL PBST for 6 times, 100 µL TMB solution was added to the plate and then stopped by 100 µL 1 M phosphoric acid. The plate was then read at OD450 with a plate reader (PerkinElmer, 2030 Multilabel Reader Victor). The result is calculated by minus OD450 value of control (no antibody).

**Immunoprecipitation of GBP-Glycan-DNA Conjugates**

For Streptavidin Magnetic beads selection: 10 µL of glycan(antigen)-DNA Conjugate (1 pmol) was incubated with 1 µL (50 µg/mL) antibody for 2 hrs at room temperature. After 2 hrs, 0.5 µL of Biotin Conjugated Secondary Antibody and 90 µL of TBST buffer (0.1 % Tween 20, 0.1% BSA) was added and incubated at room temperature for 1 hr. Using the same incubating procedure above except the secondary antibody, biotinylated-lectins (10 µg/mL) were incubated with glycan(antigen)-DNA Conjugate, DEGL-10, DEGL-50 (1 pmol). No target incubation was as control. After 1 hr the solution was incubated with 15 µL prewashed Streptavidin Magnetic beads for 30 mins at room temperature. The beads were then washed 7 times using 100 µL TBST buffer and 2 times using pure water. The washed beads were then diluted in 20 µL of pure water and eluted at 80^o^C for 10 minutes. Collect the supernatant as selected samples for detection. 1 µL of eluted samples were used as a template for PCR, qPCR or NGS.

**NGS Library Preparation**

1 µL of eluate was used for amplification using the universal template primers with Taq polymerase for 18 cycles. The PCR product was purified using Agencourt AMPure XP Magnetic beads (30% PEG) as per user instructions. The purified PCR product was amplified again using NGS adapters (30 Cycles) and purified using Agencourt AMPure XP Magnetic beads. The purified product’s concentration was measured using Agilent 2100 Bioanalyzer and/or Qubit high sensitivity dsDNA assay kit, according to manufacturer’s manual. An equimolar solution of 26 pM for ion PGM and 100 pM for ion S5 were prepared, 25 µL was loaded in Ion Chef for the Chip Preparation and sequencing by ion PGM (DEGL-10) or ion S5 (DEGL-50).

**Coding program**

Program Name: Glycan to DNA mapping webserver.

Owner: Georgia State University, Department of Chemistry.

Instructions of usage: This online program can convert glycan to DNA sequence or can reverse a DNA sequence to glycan. The output can be displayed on a new window or the user can download the out as a text file. We have hosted the program website on Georgia State University (GSU) server with CentOS 6.7 64-bit, 6x IBM System x3850 x5, Intel Xeon Processor E7-4850, 4 CPUs (10 cores per Core Processing Unit), 2.0 gigahertz (GHz) processors, 512 gigabytes (GB) Random Access Memory and 2 terabytes (TB) of scratch storage for jobs. Instructions to use the program are provided in the web portal.

**Data analysis**

Program Name: Sequence count analysis.

Owner: Georgia State University, Department of Chemistry.

Instructions of usage:

This python program takes as many FASTAQ files and generates count of each unique sequences provided in a single excel sheet, then list the count of each sequence in excel file output. It uses Python Packages Panda and OS to perform counting and reading FASTAQ and exporting counts to CSV file.

### 2. Figures

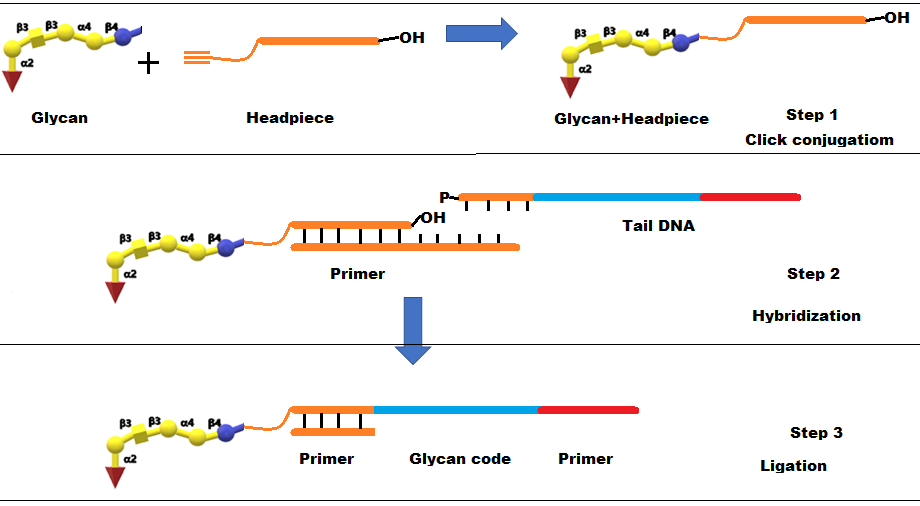

**Figure S1.** Schematic representation of monovalent DEGL synthesis.

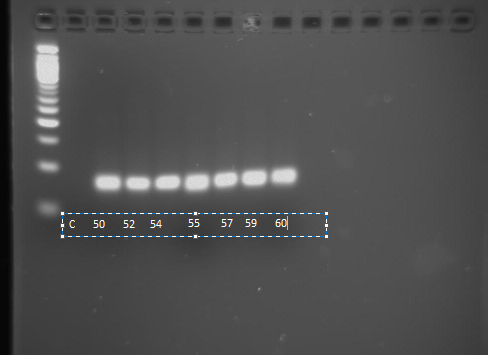

**Figure S2.** Gradient PCR for Tm determination.

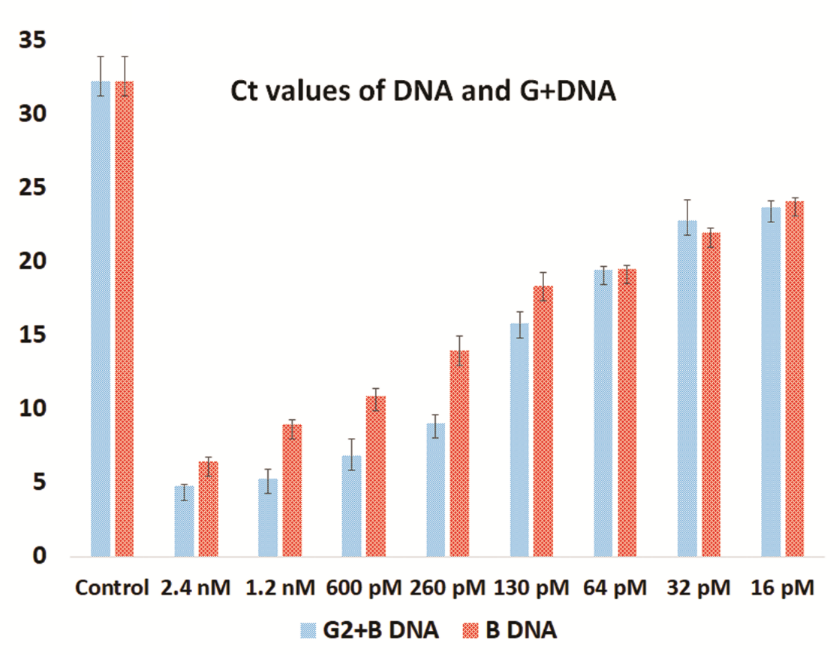

**Figure S3.** Ct value comparison of pure DNA and glycan conjugated DNA

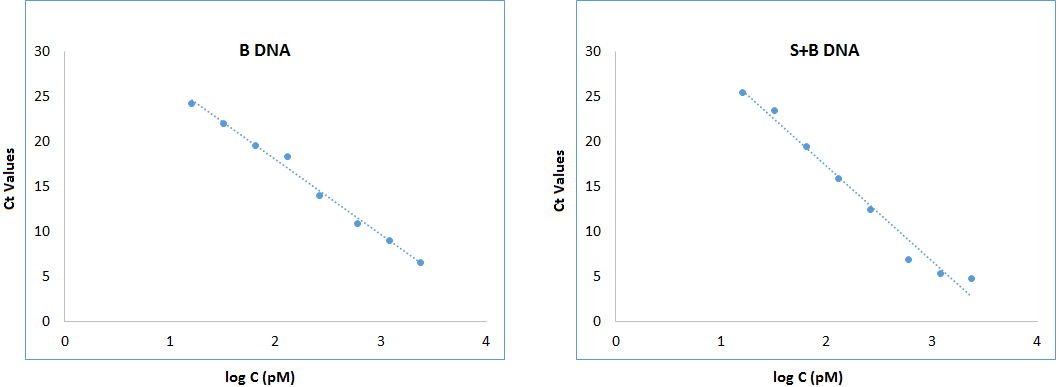

**Figure S4.** Standard curve plot of pure DNA (B DNA) and glycan conjugated B DNA.

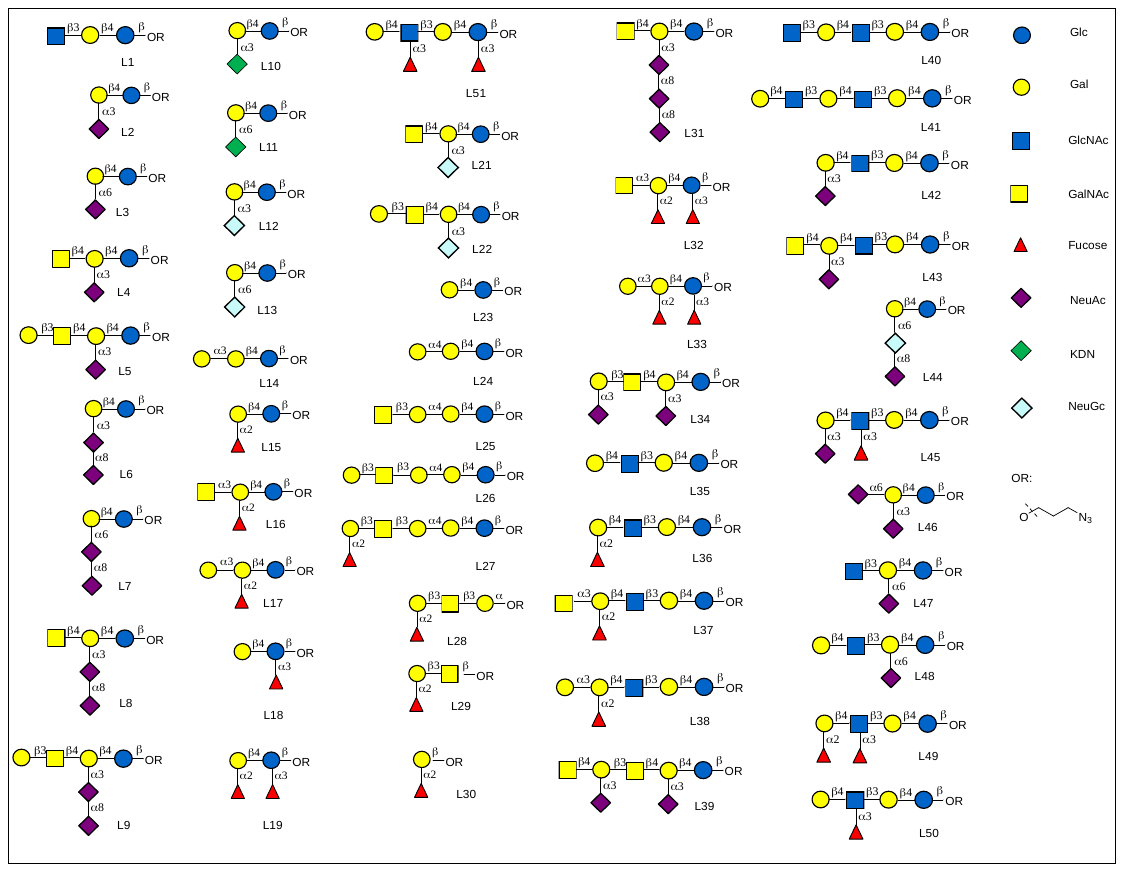

**Figure S5**. Glycan structures used for the library preparation.

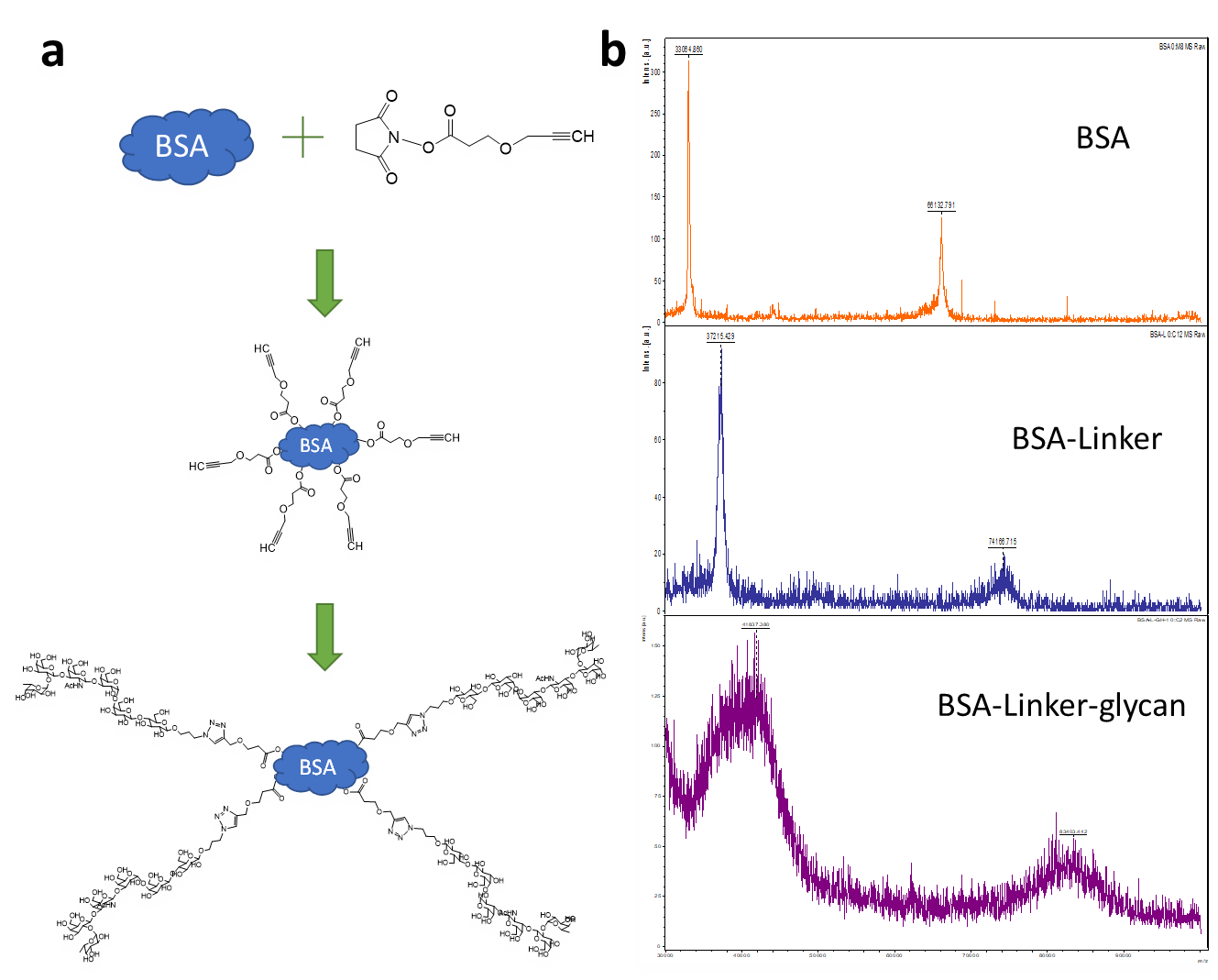

**Figure S6.** Synthesis of indirect glycan antigen for ELISA. To prepare coating protein BSA-glycans, firstly we prepared the BSA-linker which contains alkynyl, allowing the followed click reaction with glycan-N3. The Propargyl-N-hydroxysuccinimidyl ester (10 mg/mL in PBS) was added to the same volume of BSA (20 mg/mL in PBS) and the PH was adjusted to 8.5 using triethylamine. The mixture was stirred at RT for 2 h. Then the solution containing BSA-linker(alkynyl) was ultrafiltrated and washed with PBS (pH 7.4) using 3 KDa centrifugal filter (Millipore). The corresponding linker moieties loading ratio was 73 by MALDI-TOF mass spectrometry analysis. After getting the BSA-linker, the click reaction with glycan-N3 (20-fold of BSA-linker mole) was conducted by adding THPTA, CuSO4 and ascorbic acid at room temperature overnight. Again, the solution containing BSA-glycans was ultrafiltrated and washed with PBS (pH 7.4) using 3 KDa centrifugal filter (Millipore). The corresponding linker moieties loading ratio was ~10 by MALDI-TOF mass spectrometry analysis.

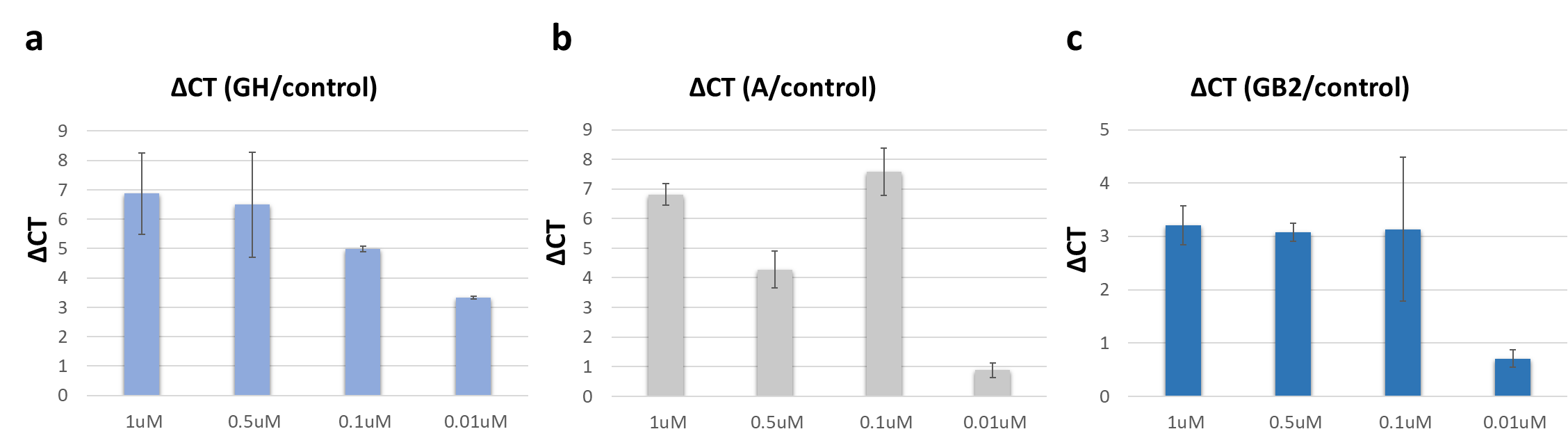

**Figure S7.** qPCR assay for singleplex selection of multivalent glycan conjugates (MHP-DNA-glycan) with different concentration. **a.** qPCR assay for different concentration of MHP-DNA-GH selection with VK9 antibody. The difference of Ct value (ΔCT) between the glycan vs target and no target control in the binding was over 5 even at the lower concentration of 100 nM. **b.** qPCR assay for different concentration of MHP-DNA-A selection with HPA. The difference of Ct value (ΔCT) between the glycan vs target and no target control in the binding was over 7 even at the lower concentration of 100 nM, while under 1 at the concentration of 10 nM. **c.** qPCR assay for different concentration of MHP-DNA-Gb2 selection with HPA. The difference of Ct value (ΔCT) between the glycan vs target and no target control in the binding was over 3 even at the lower concentration of 100 nM, while under 1 at the concentration of 10 nM.

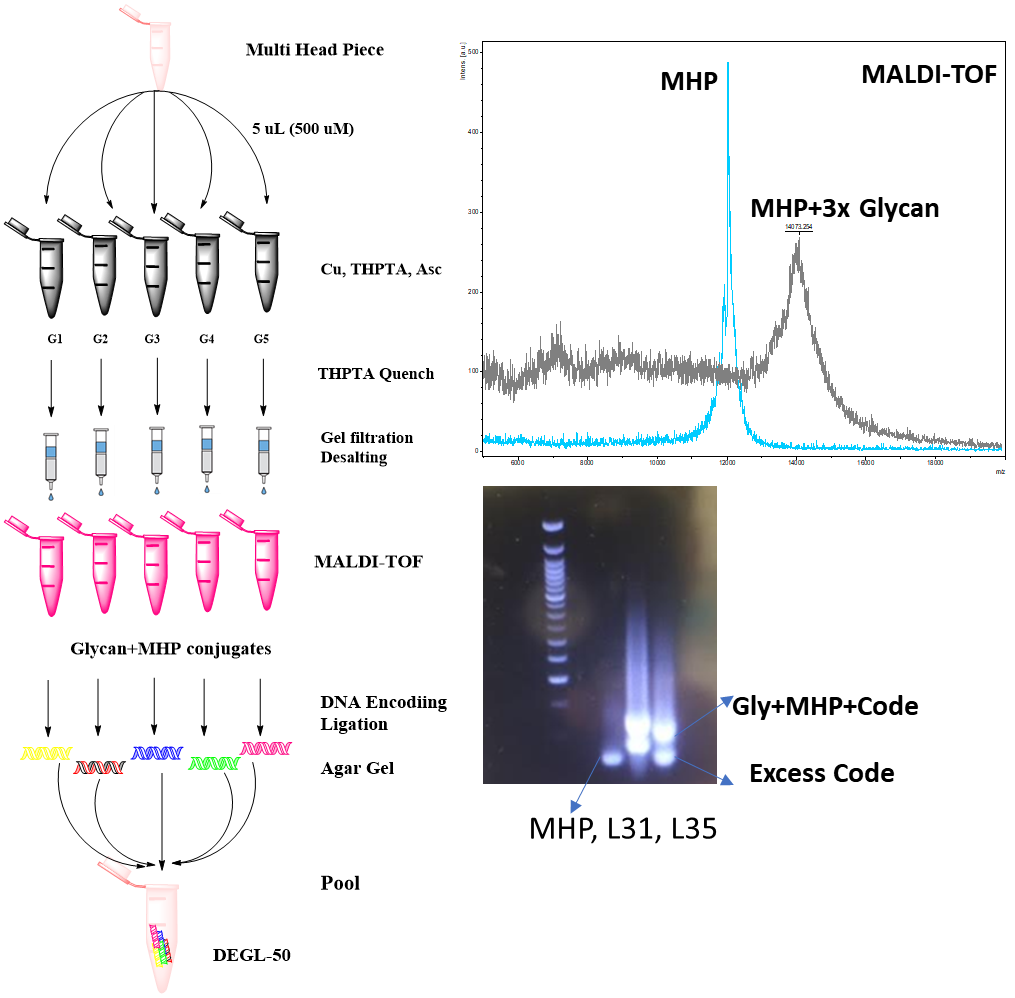

**Figure S8.** General protocol for the high content DEGL. HPLC purified headpiece were split into 50 wells (5 µL/ 500 µM), followed by glycans and click reagents. Following the reaction, excess reagents were filtered out using the gel filtration column. Products eluted to prelabelled tubes and evaluated using MALDI-TOF, concentration was measured using nanodrop. Next, 20 µM of the glycan+MHP conjugates were ligated with the corresponding glycan’s DNA code to final concentration 10 µM. Resulting glycan-MHP-DNA code were tested using agarose gel, then pooled to get the DEGL stock solution. 100 nM solutions were used in all selection procedures.

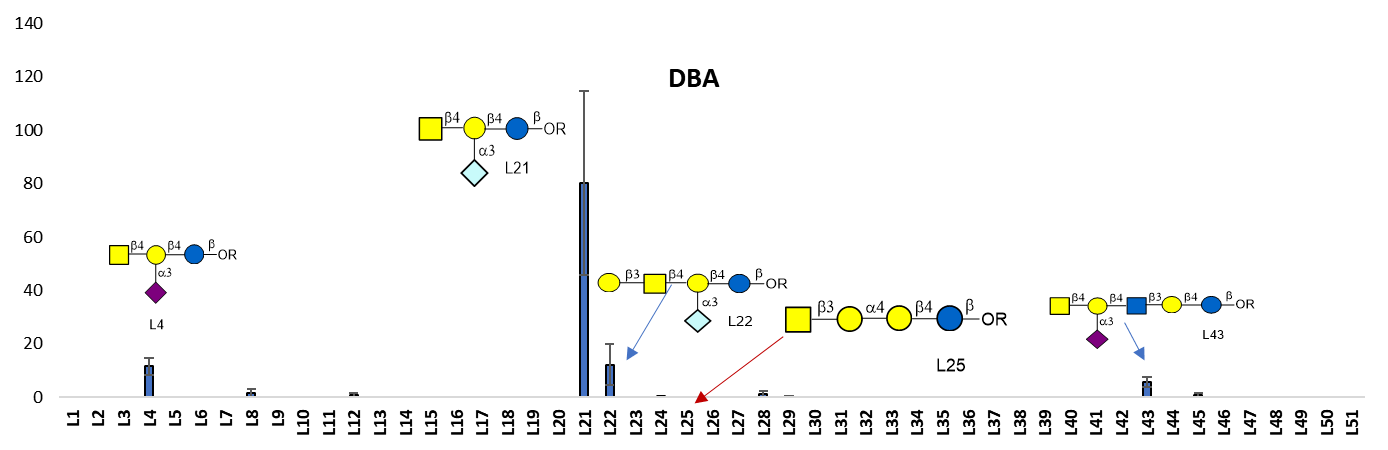

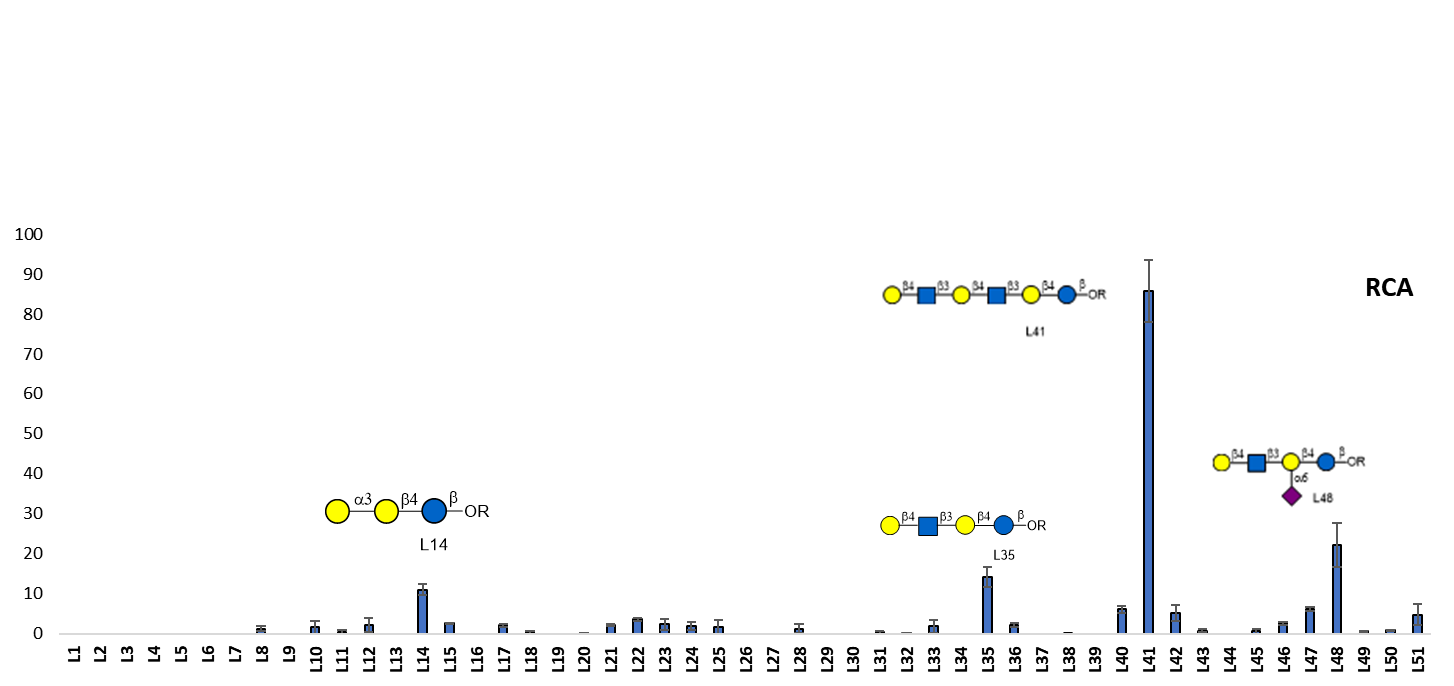

Enrichment fold

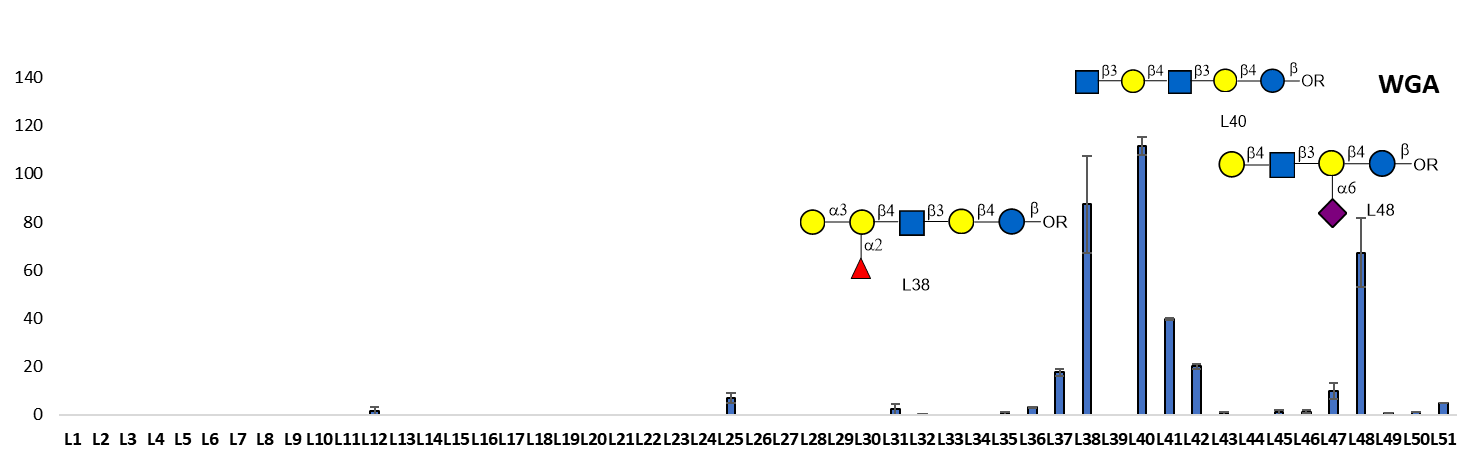

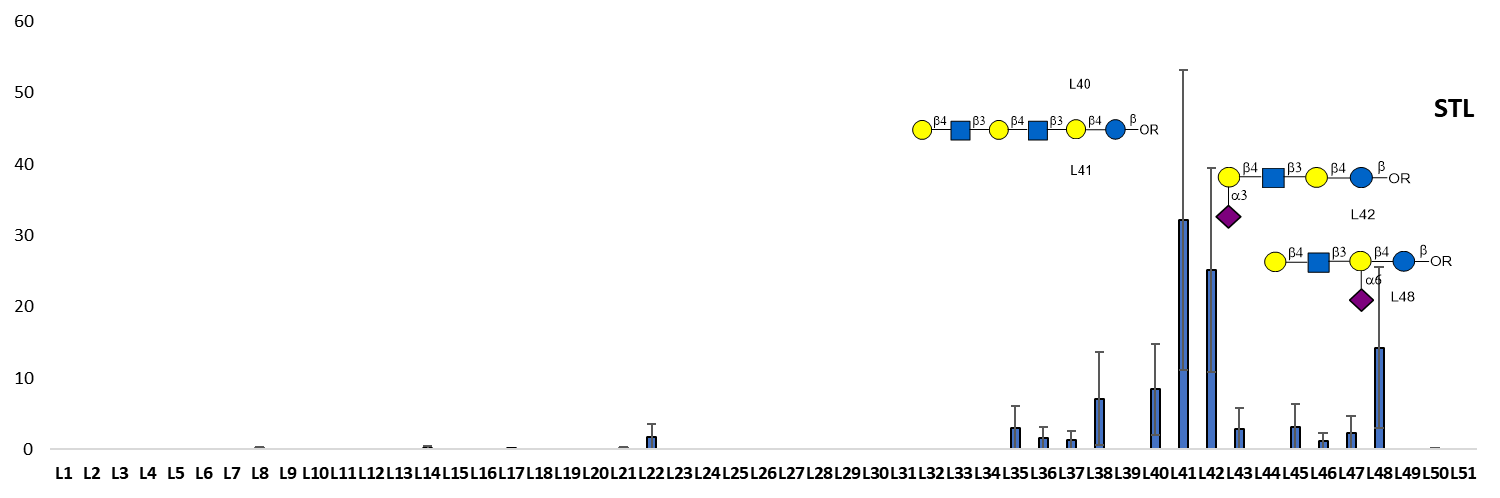

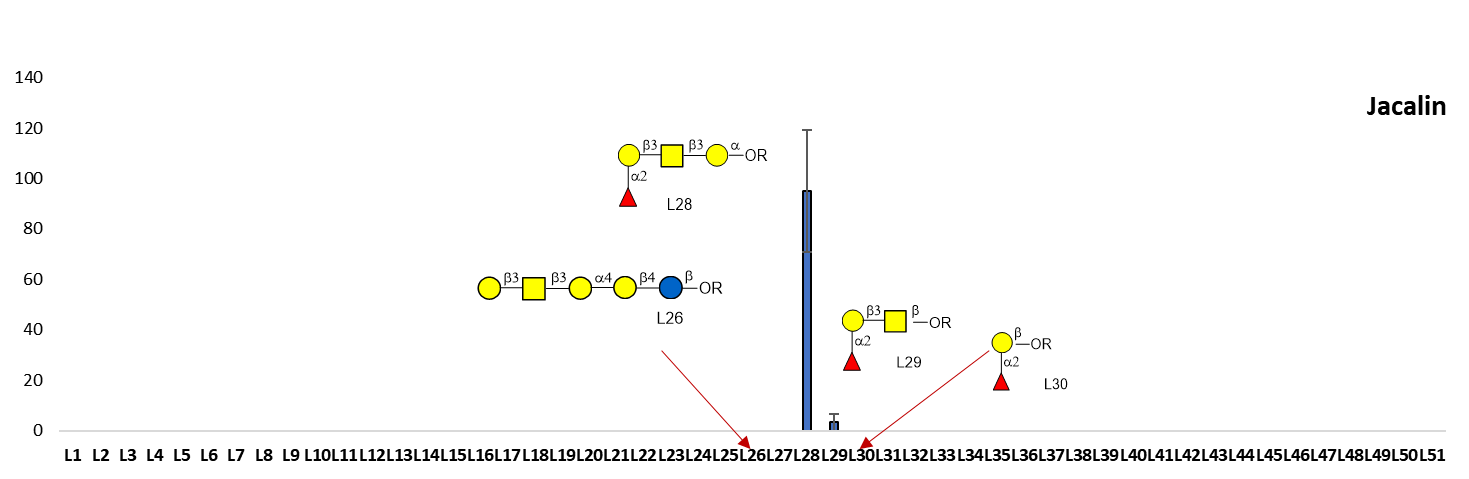

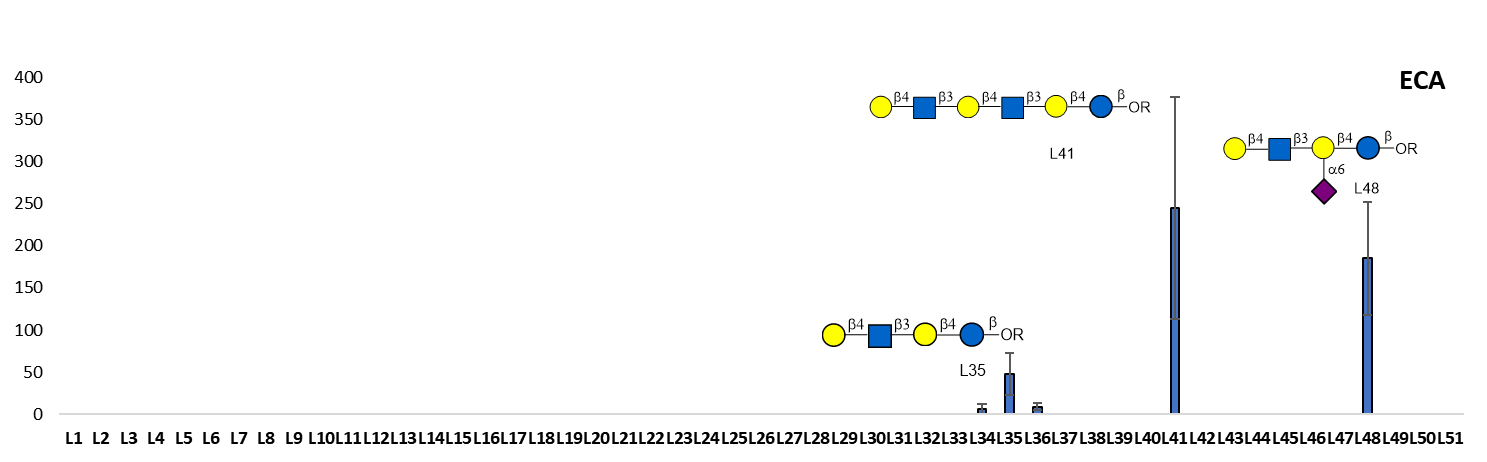

Enrichment fold

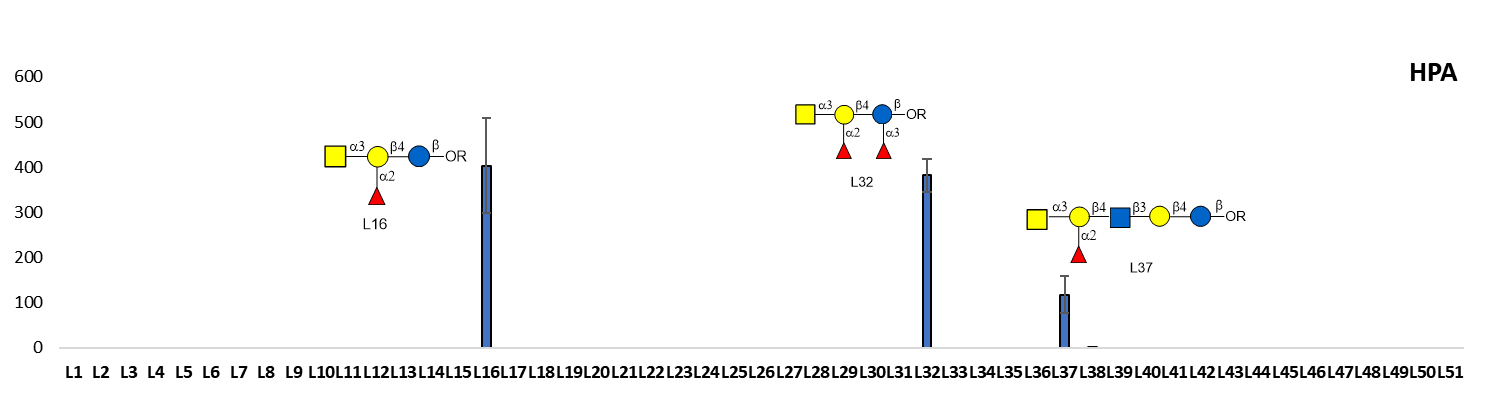

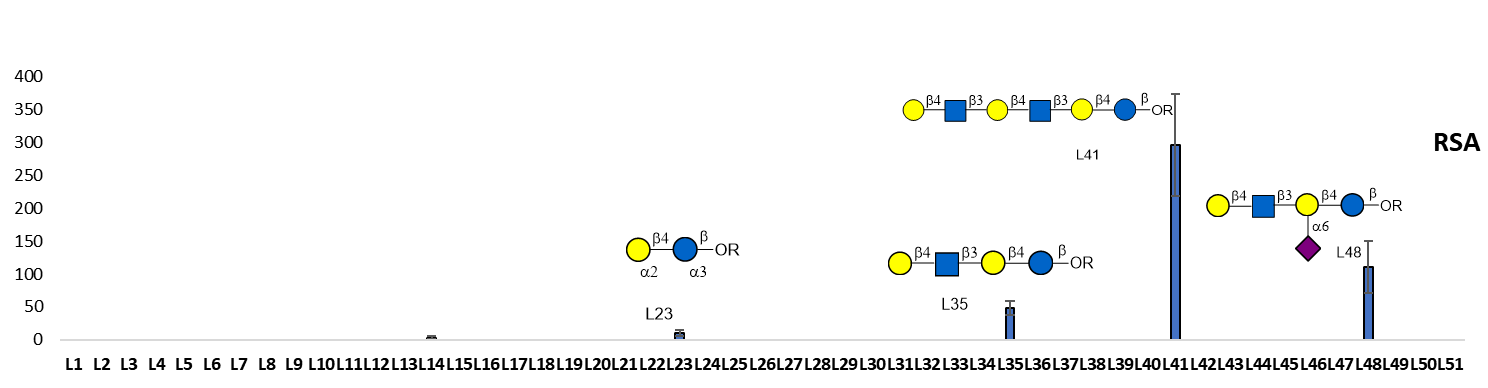

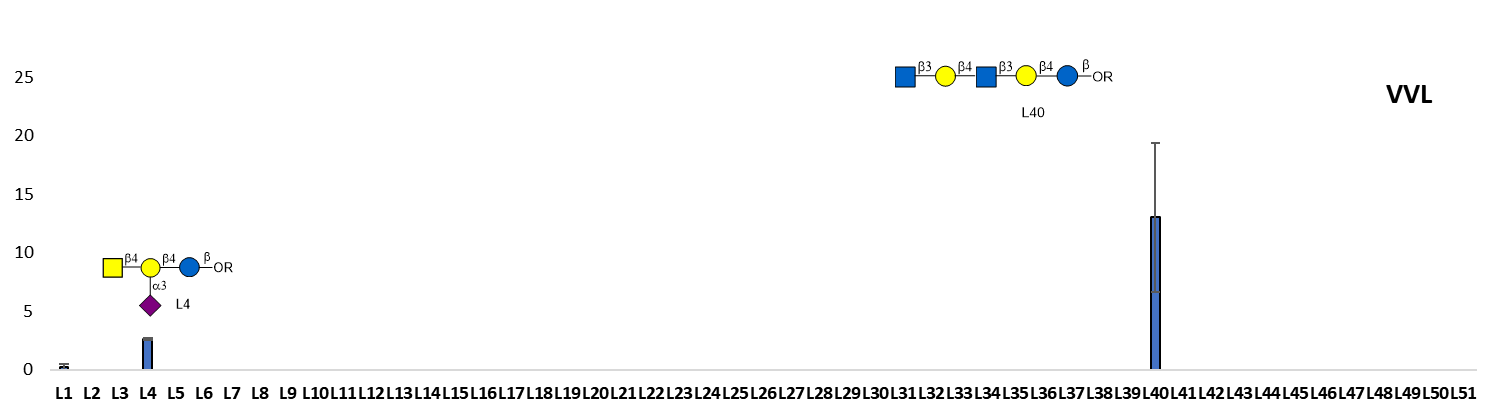

Enrichment fold

**Figure S9.** Binding profiles of lectins. The data is presented as mean ± SEM of two replicates, each experiment was repeated 3 times.

### 3. MALDI TOF MS and HPLC results

#### MALDI TOF MS of monovalent Headpiece-glycan conjugates

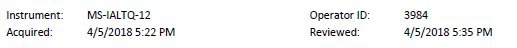

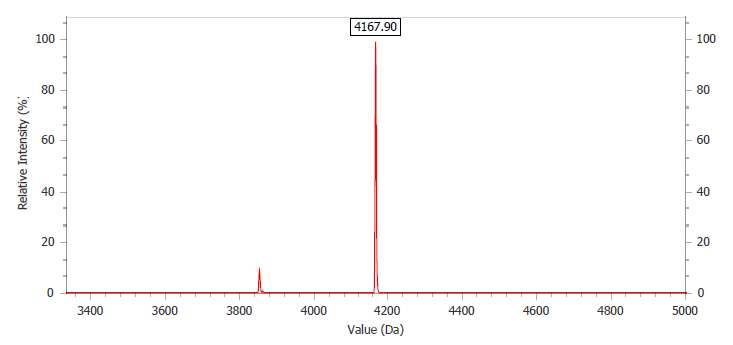

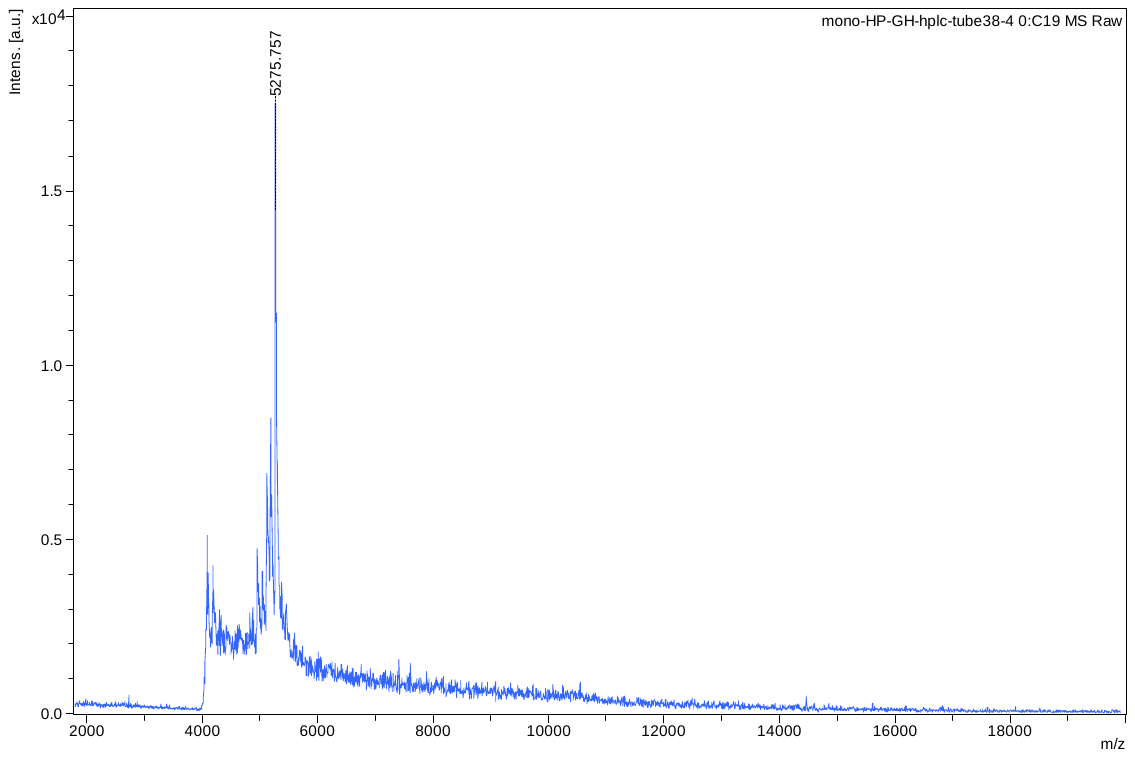

Headpiece DNA+GH

Headpiece DNA

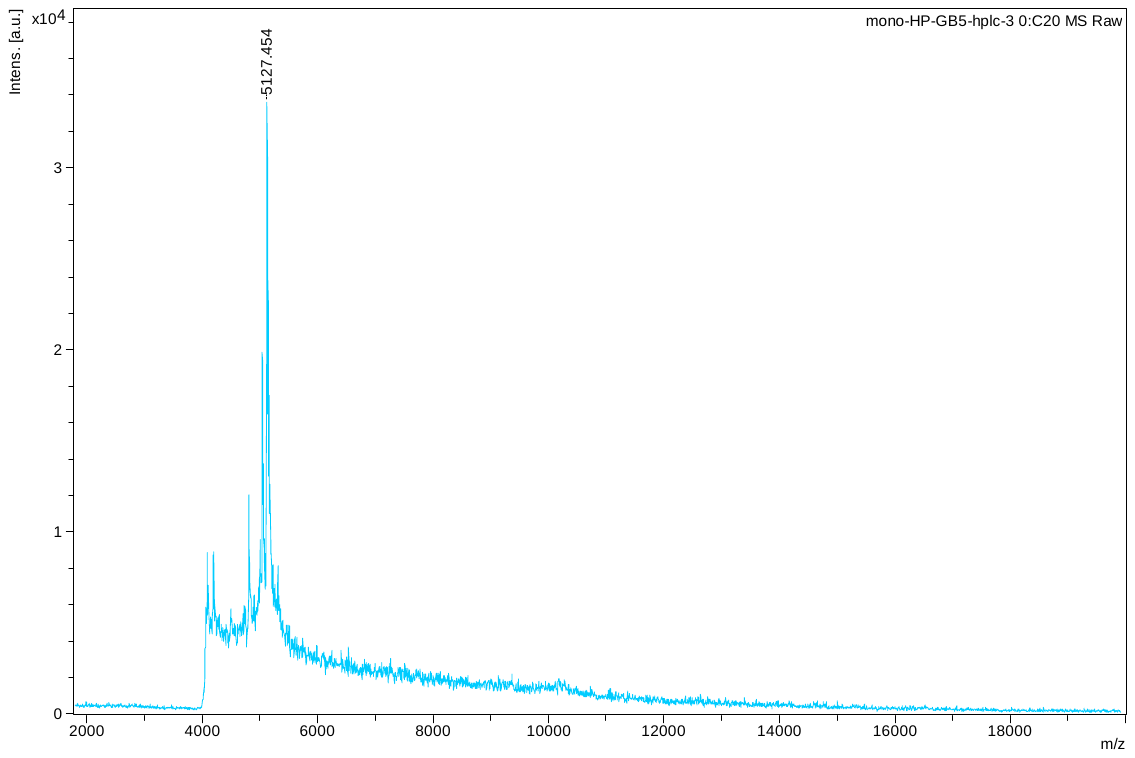

Headpiece DNA + Gb5

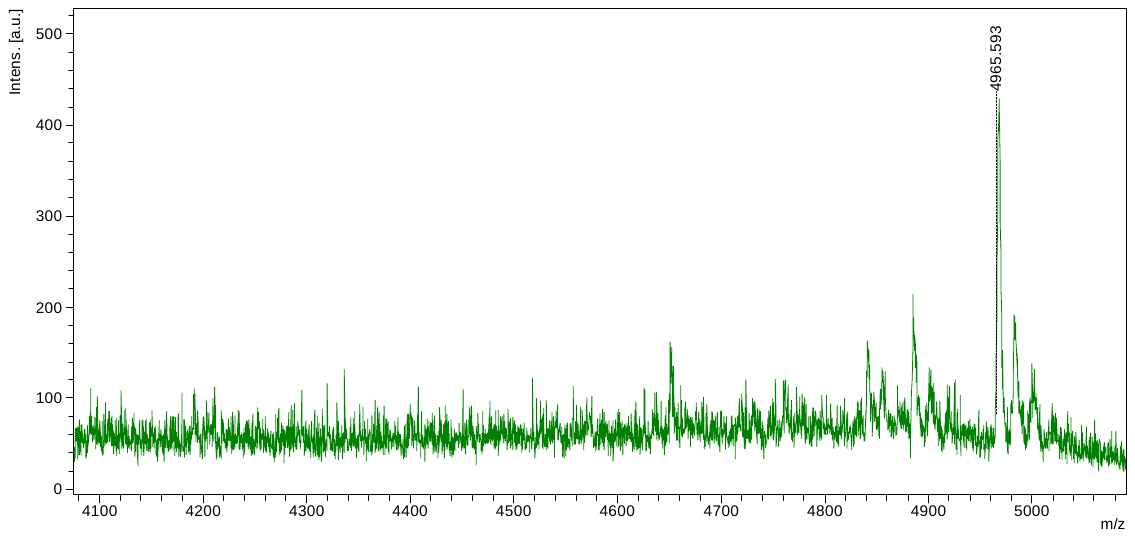

Headpiece DNA+Gb4

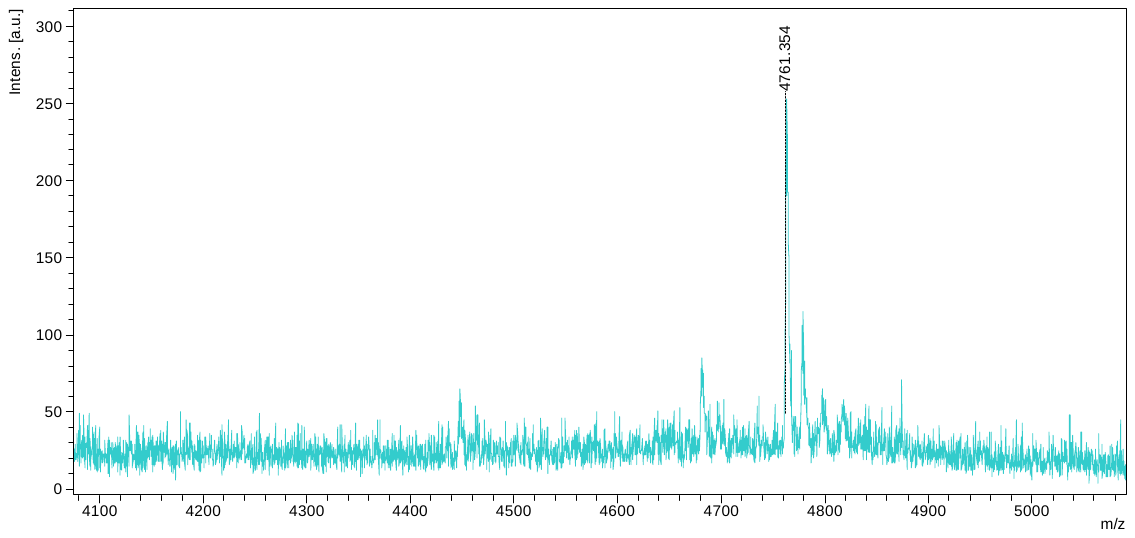

Headpiece DNA+Gb3

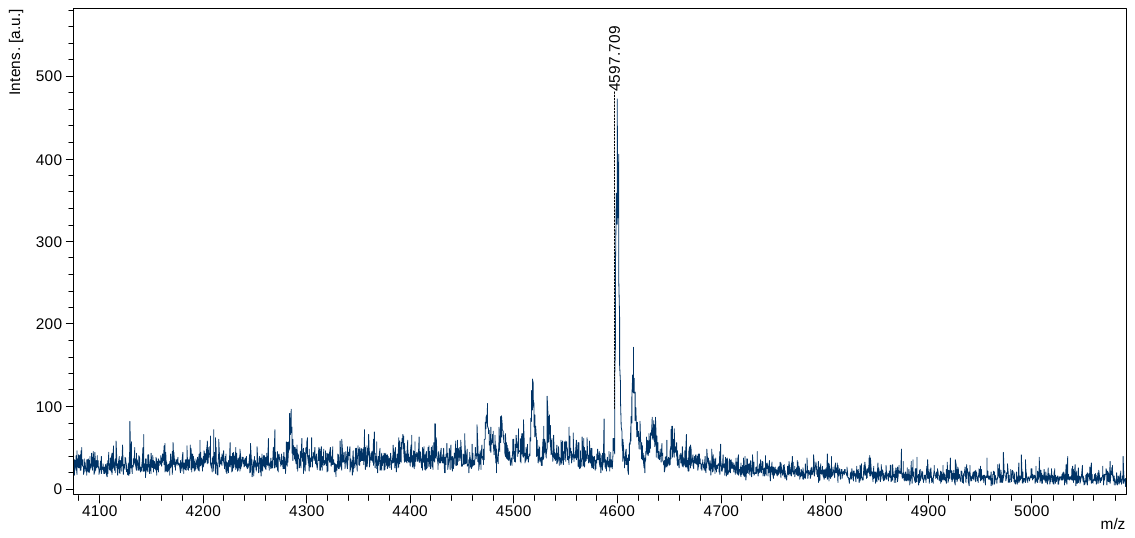

Headpiece DNA+Gb2

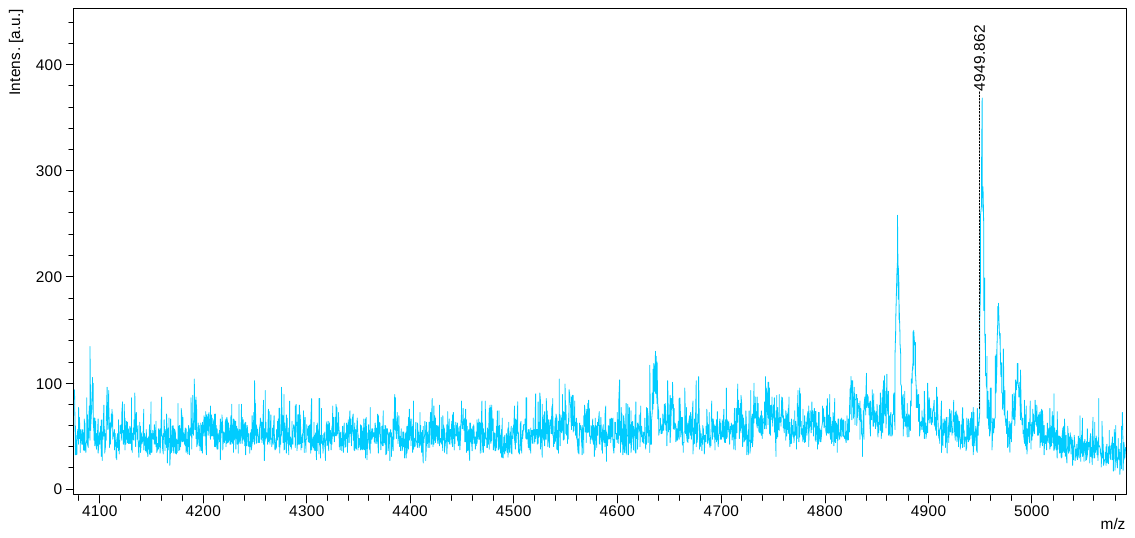

Headpiece DNA+Bb4

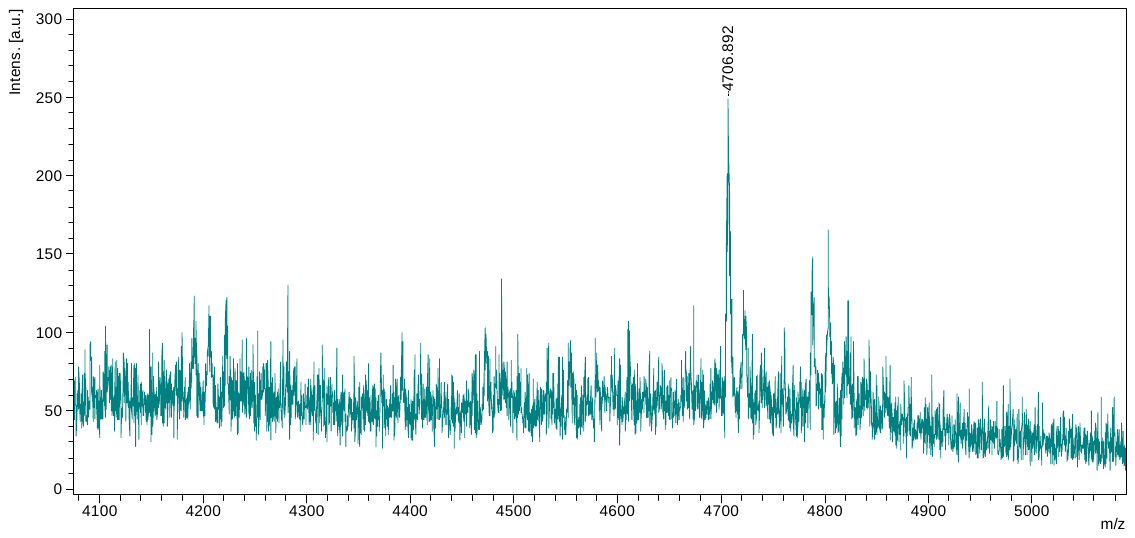

Headpiece DNA+BB3

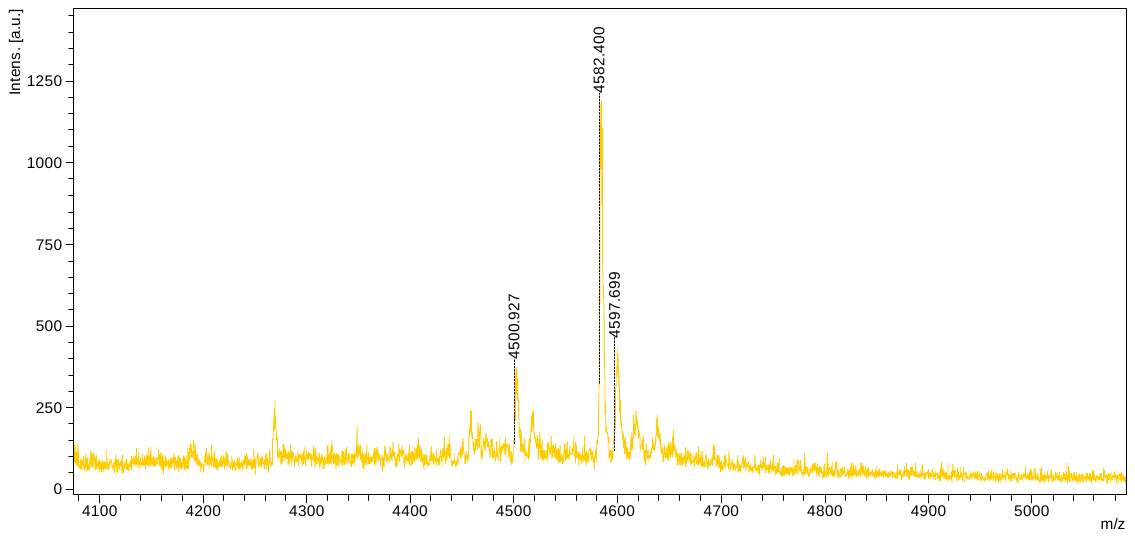

Headpiece DNA+Bb2

#### MALDI TOF MS of Multivalent Headpiece-glycan conjugates

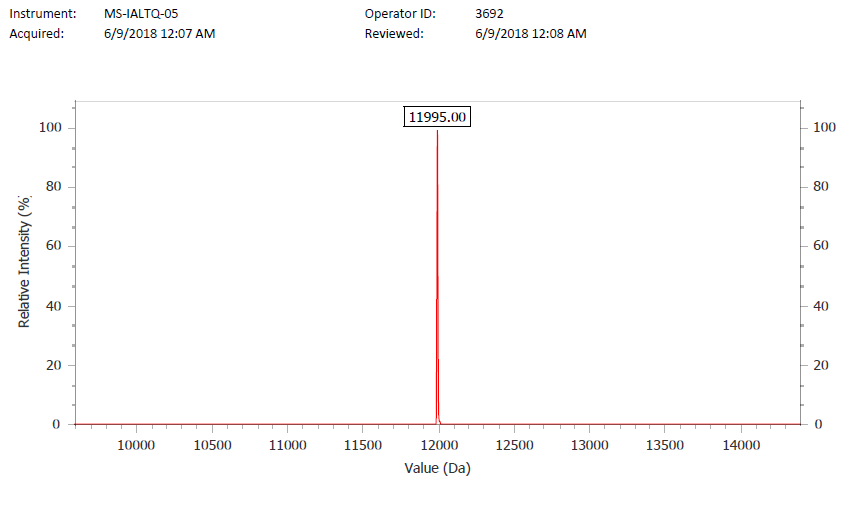

Multi Headpiece (MHP)

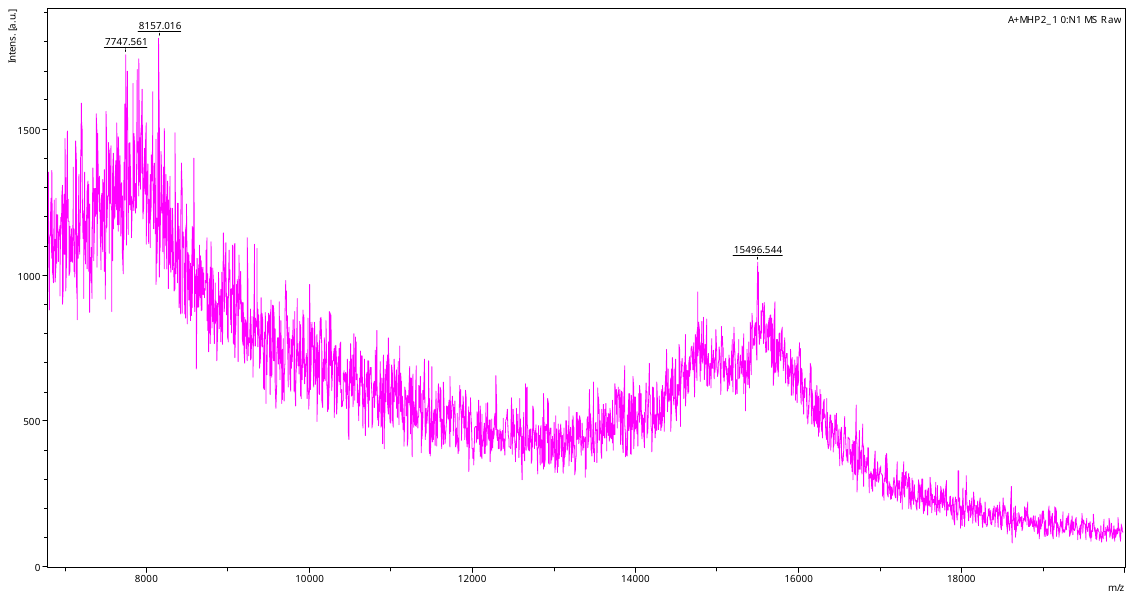

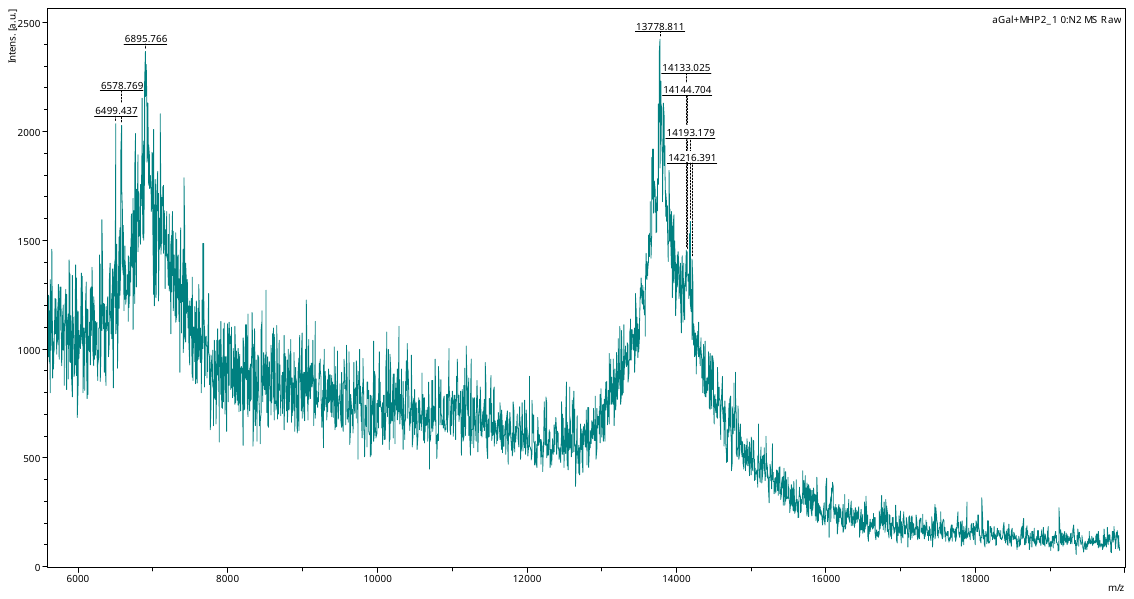

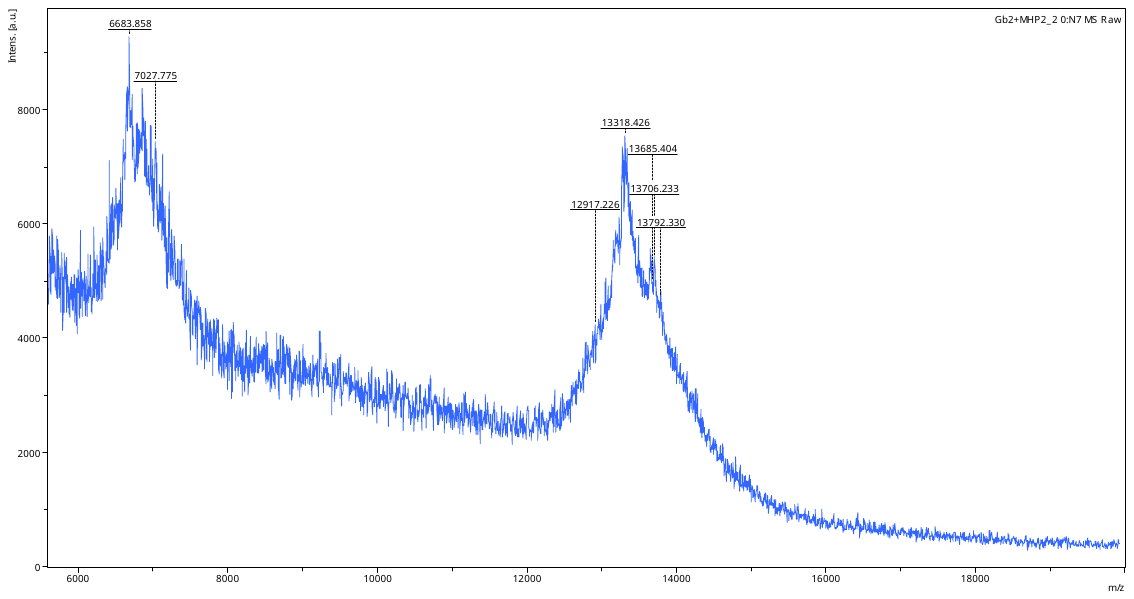

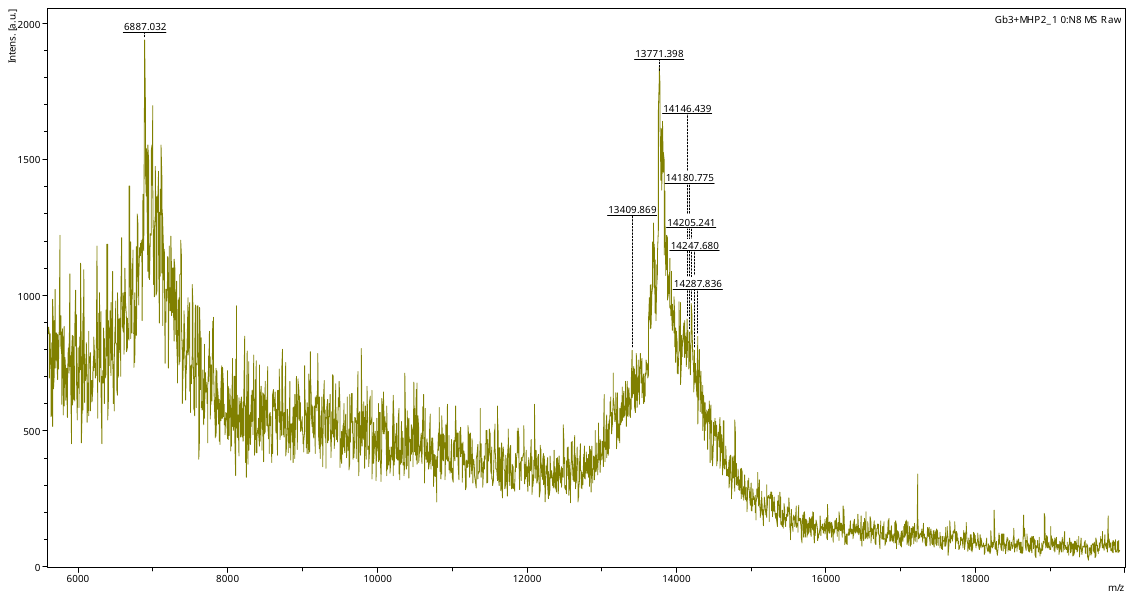

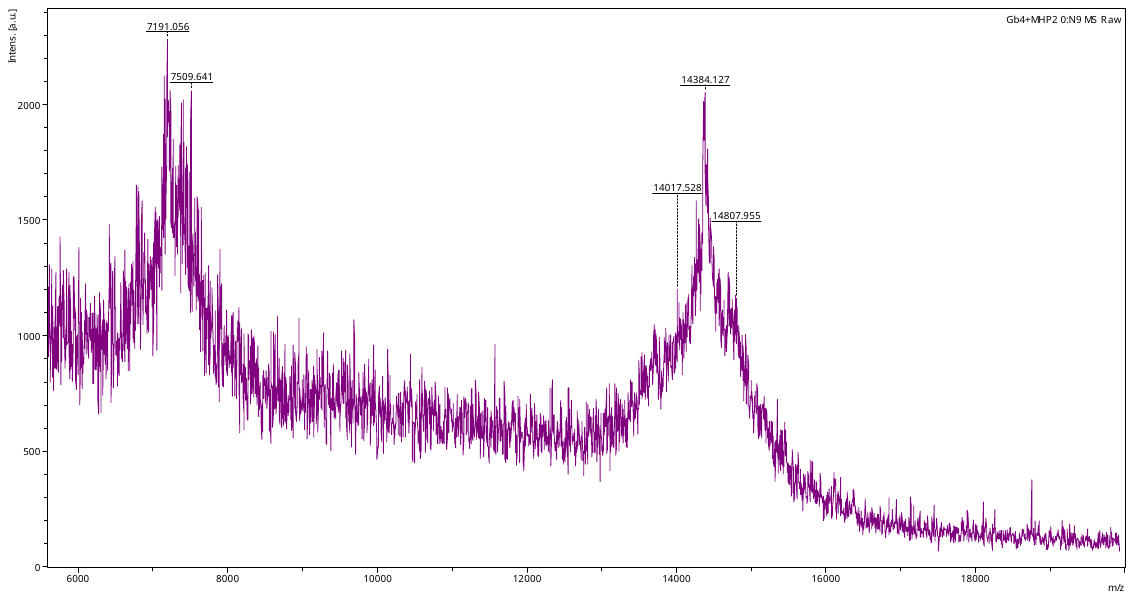

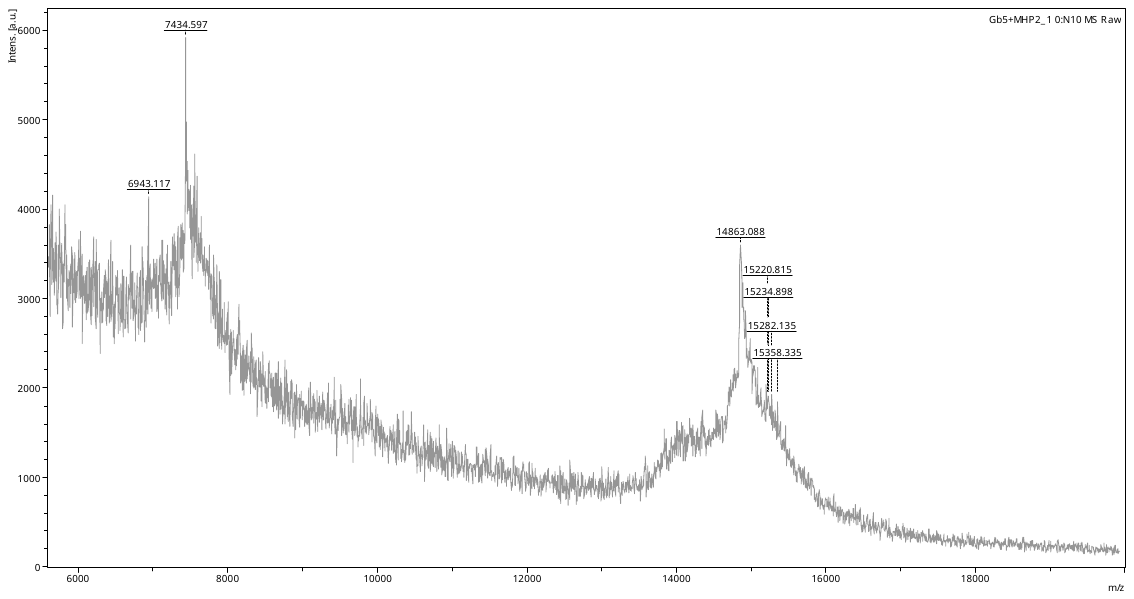

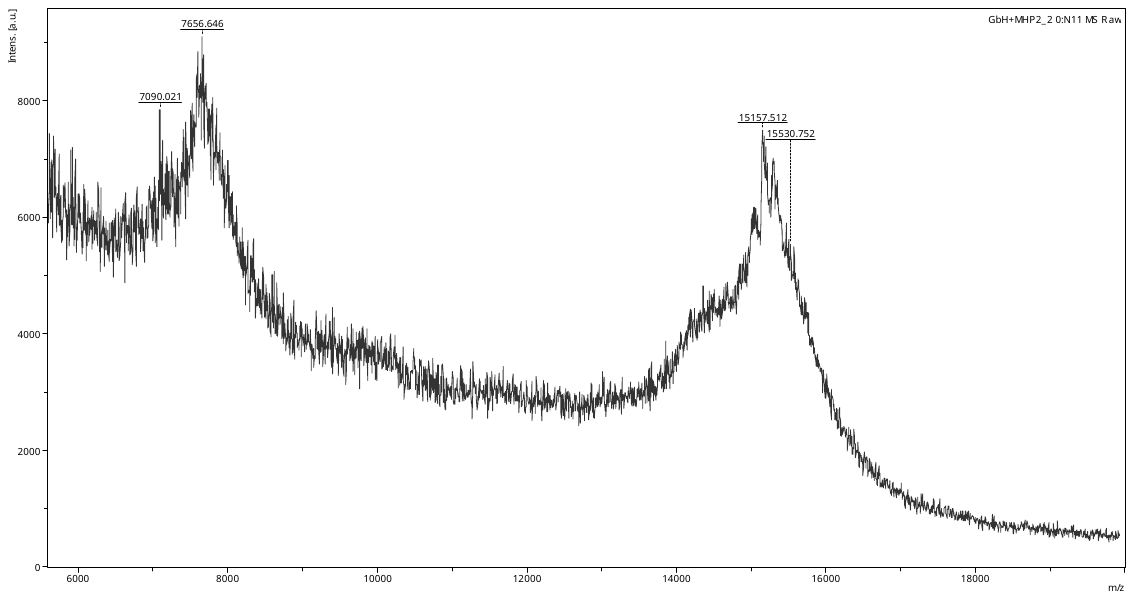
­­
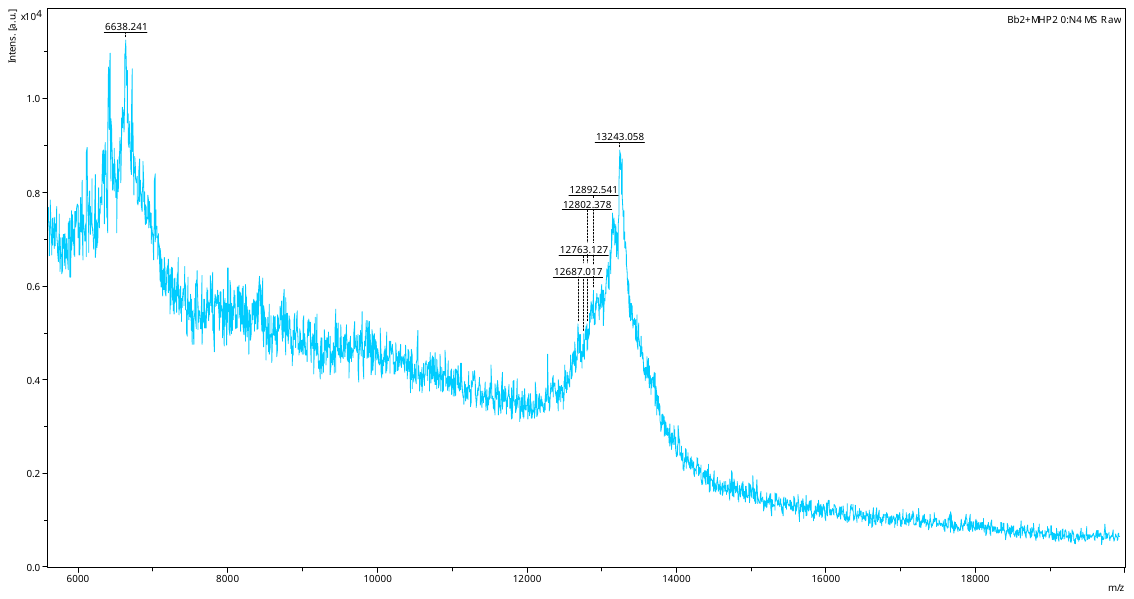

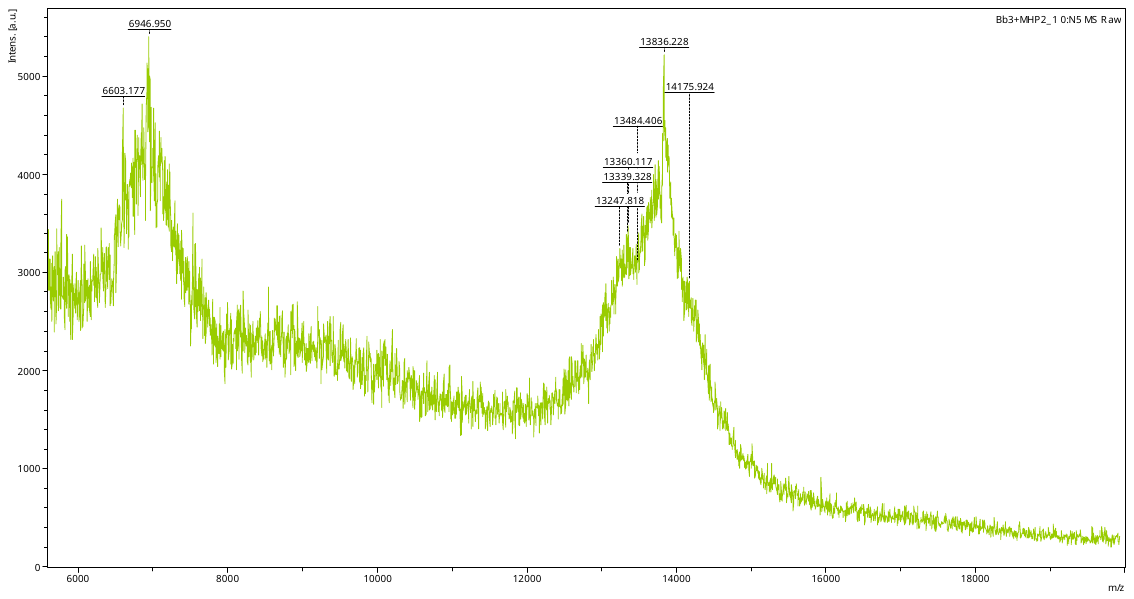

Multi Headpiece (MHP)+A (L37)

MHP+aGal (L14)

MHP+Gb2 (L23)

MHP+Gb3 (L24)

MHP+Gb4 (L25)

MHP+Gb5 (L26)

MHP+GH (L27)

MHP+Bb2 (L30)

MHP+Bb3 (L29)

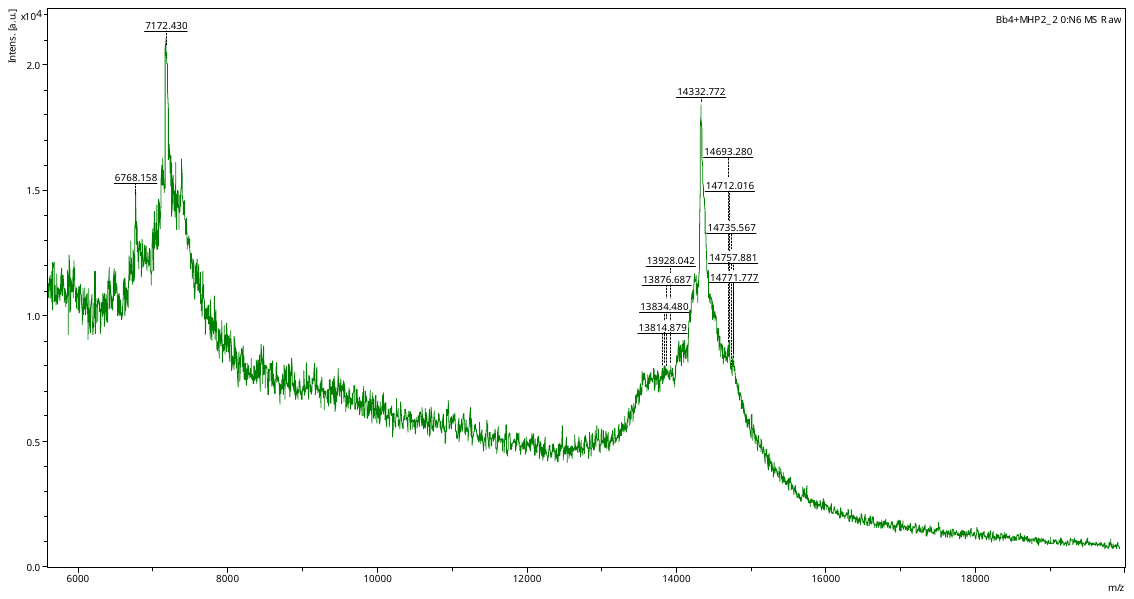

MHP+Bb4 (L28)

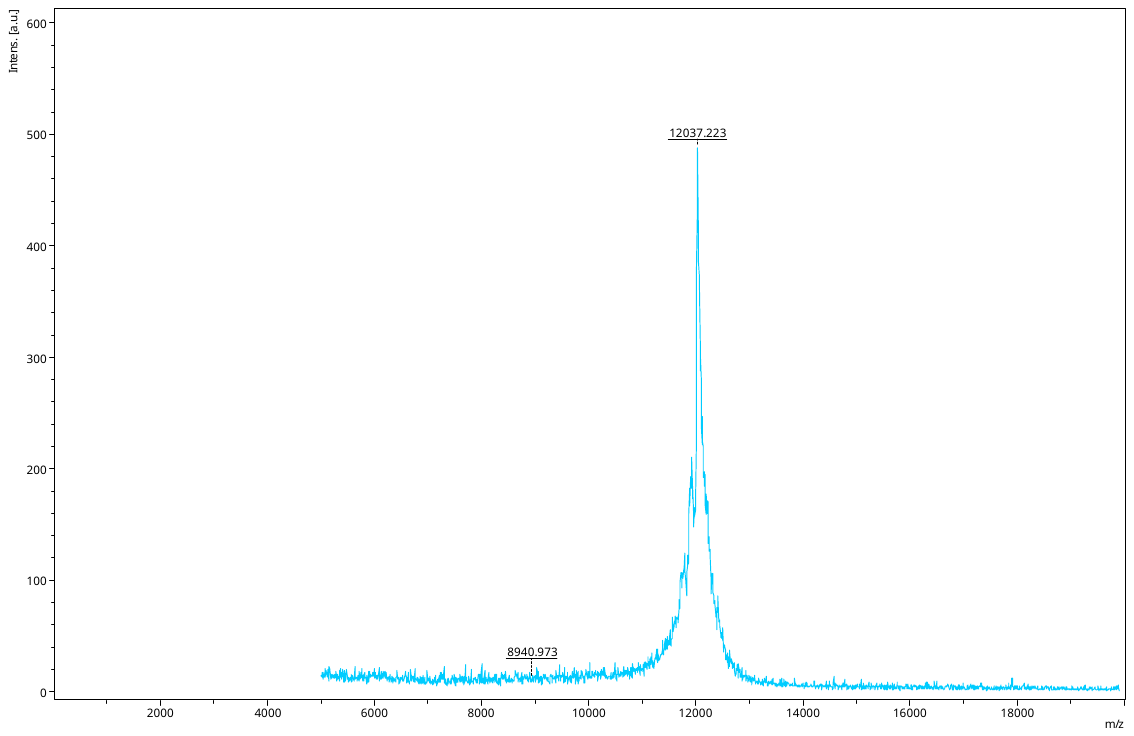

MHP2

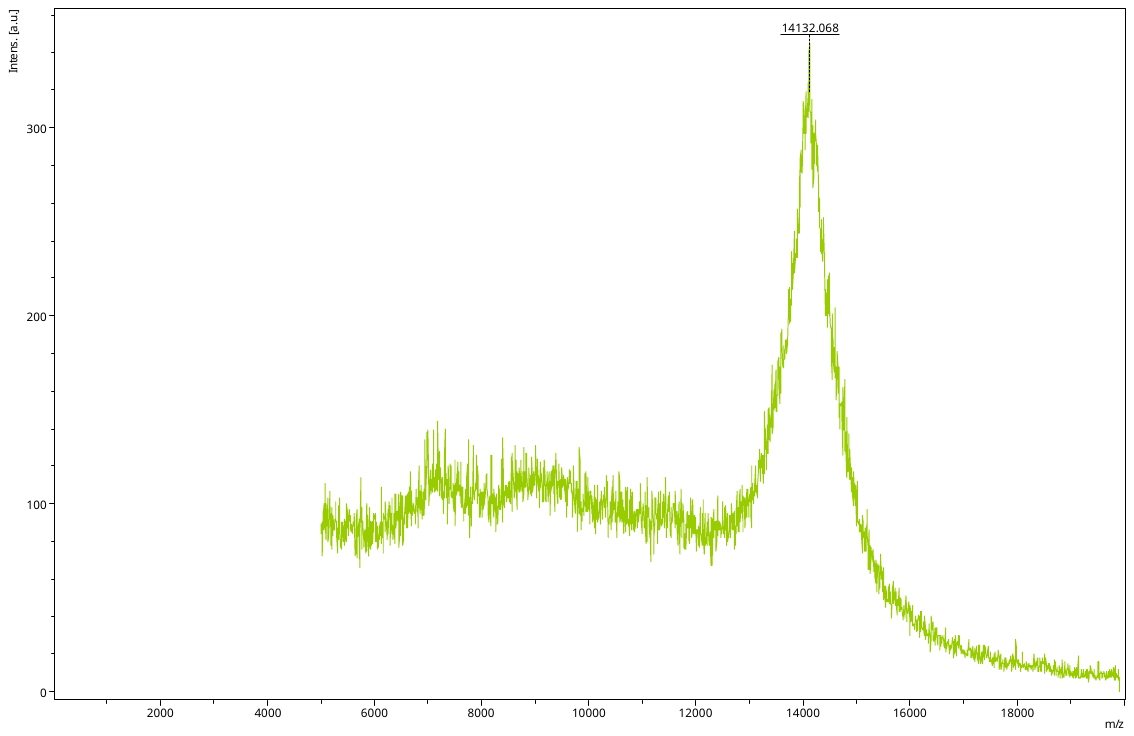

L1+MHP2

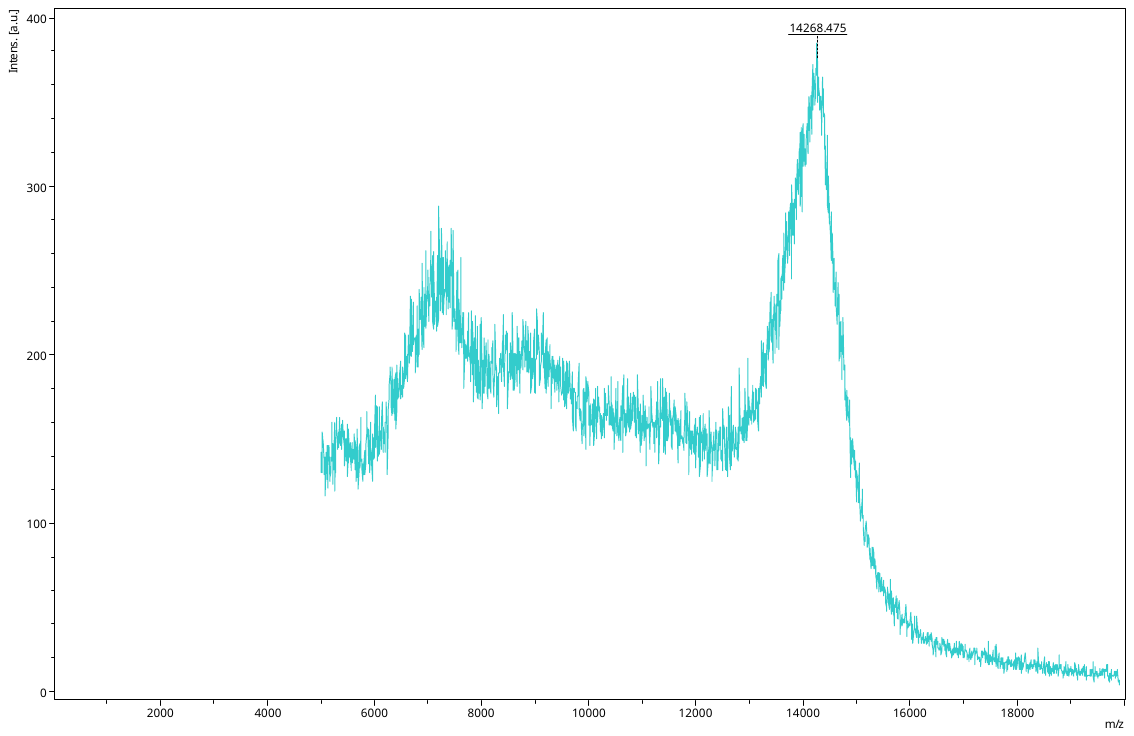

L2+MHP

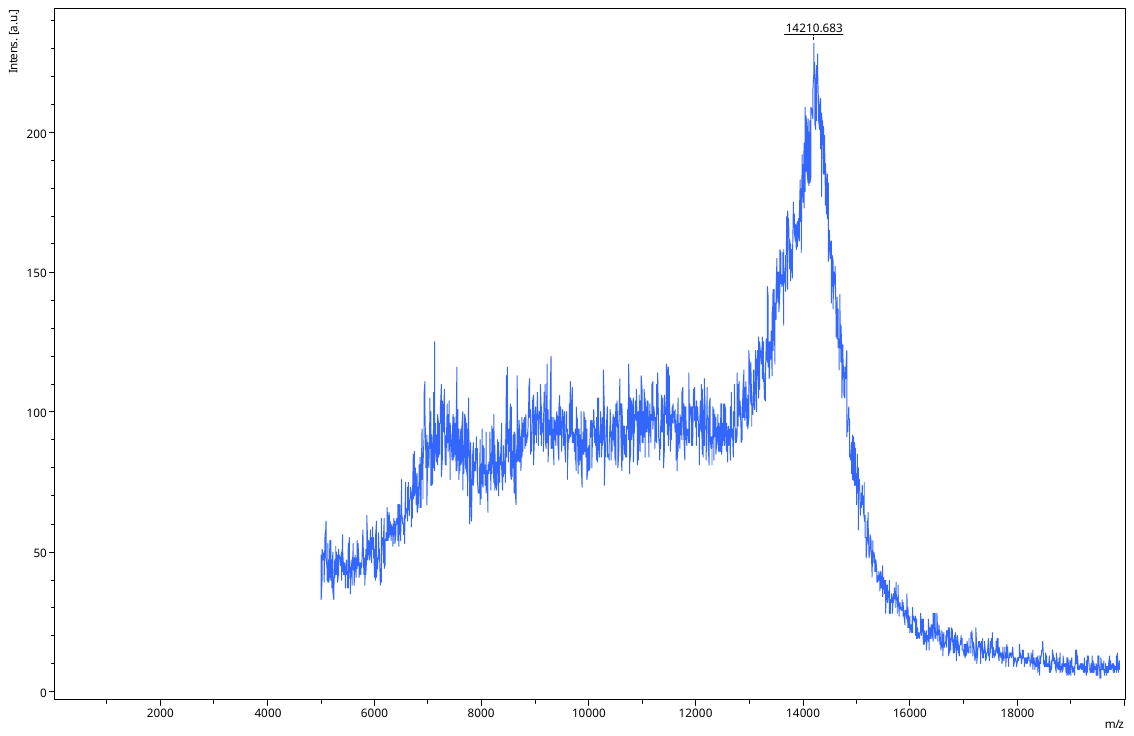

L3+MHP

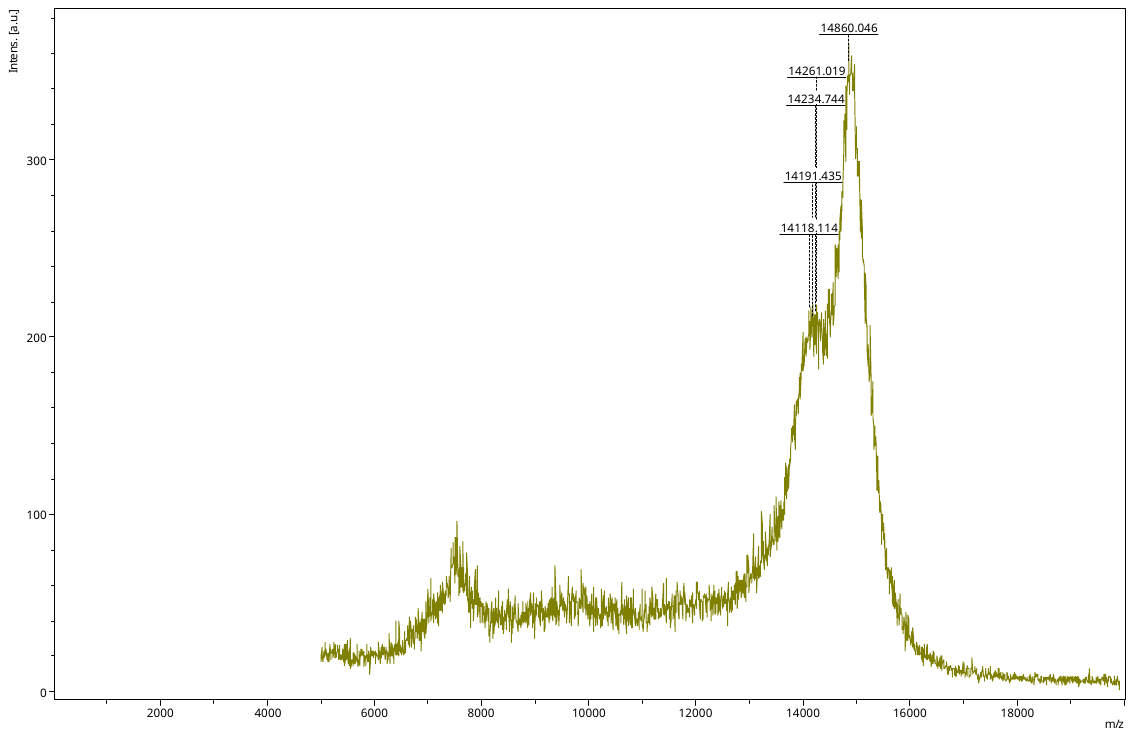

L4+MHP

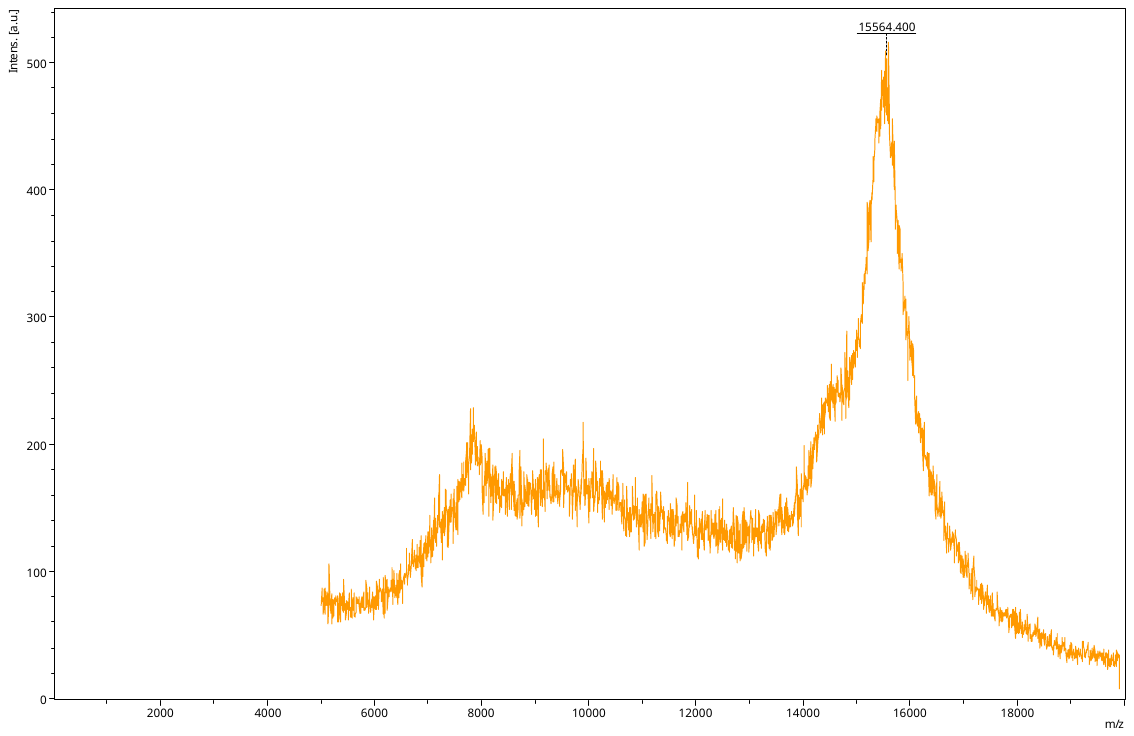

L5+MHP

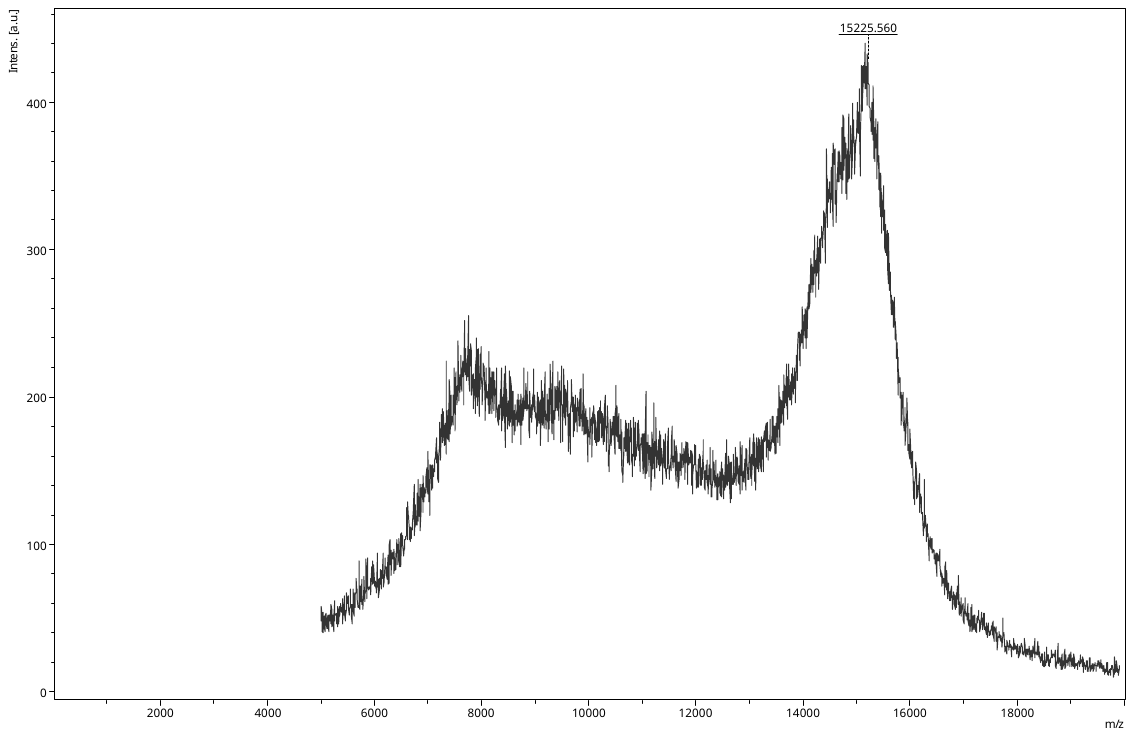

L6+MHP

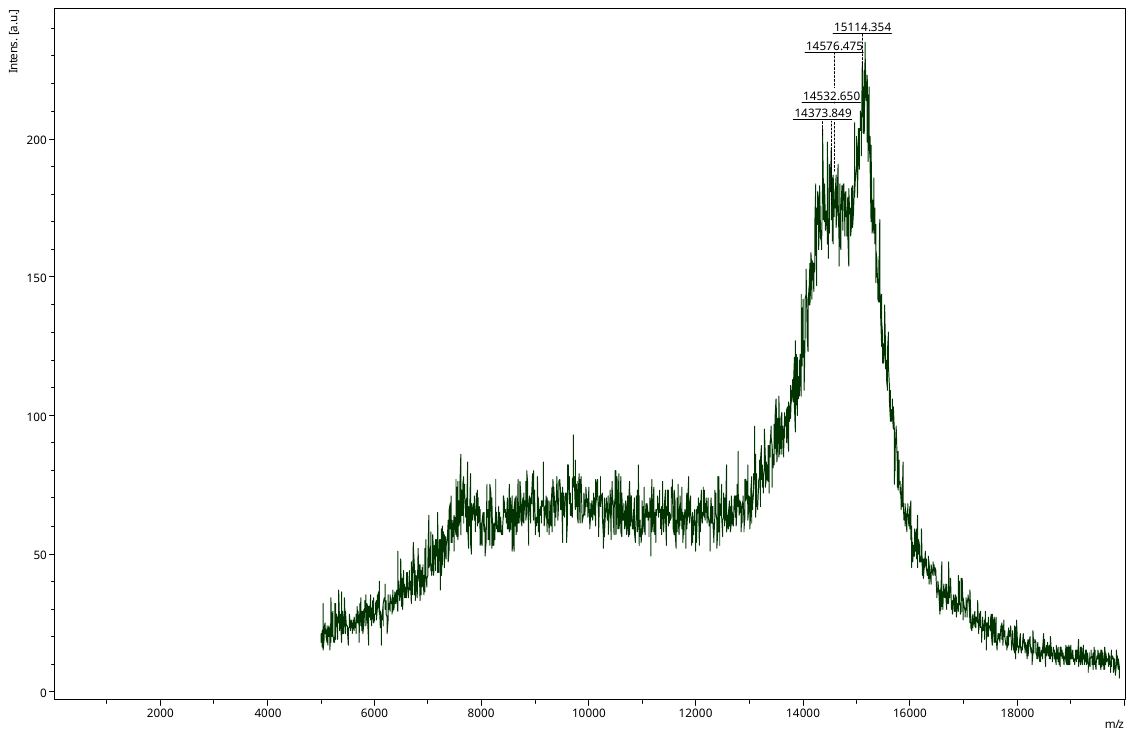

L7+MHP

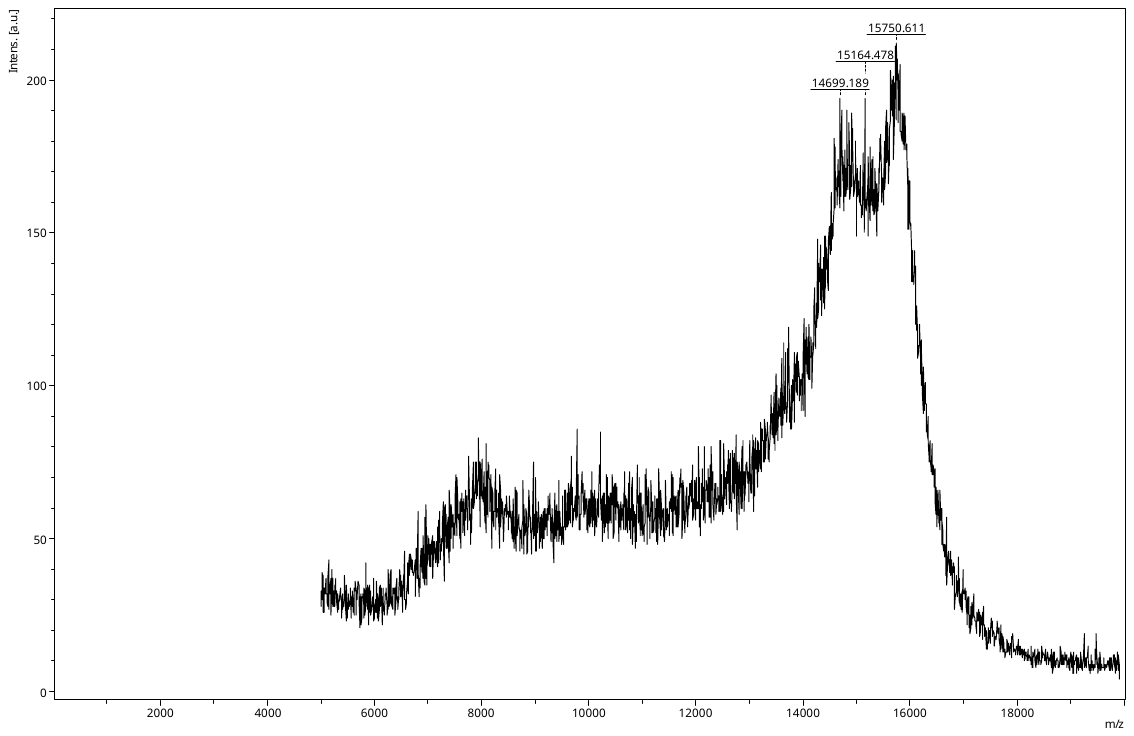

L8_MHP

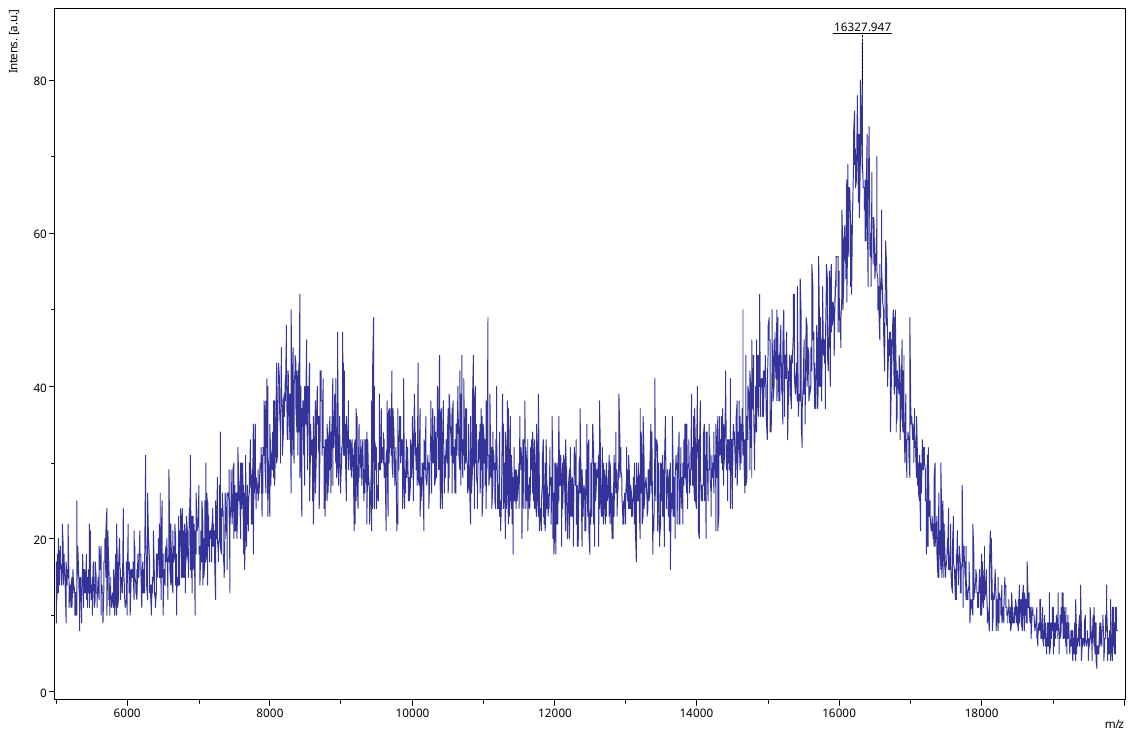

L9+MHP

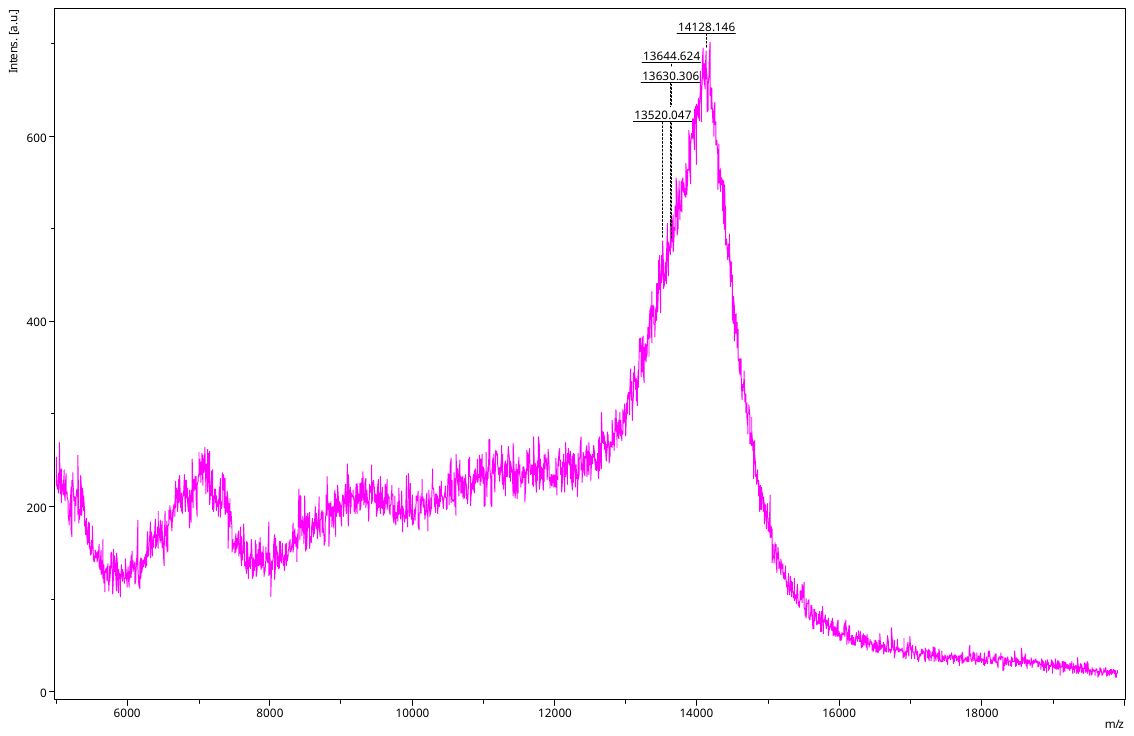

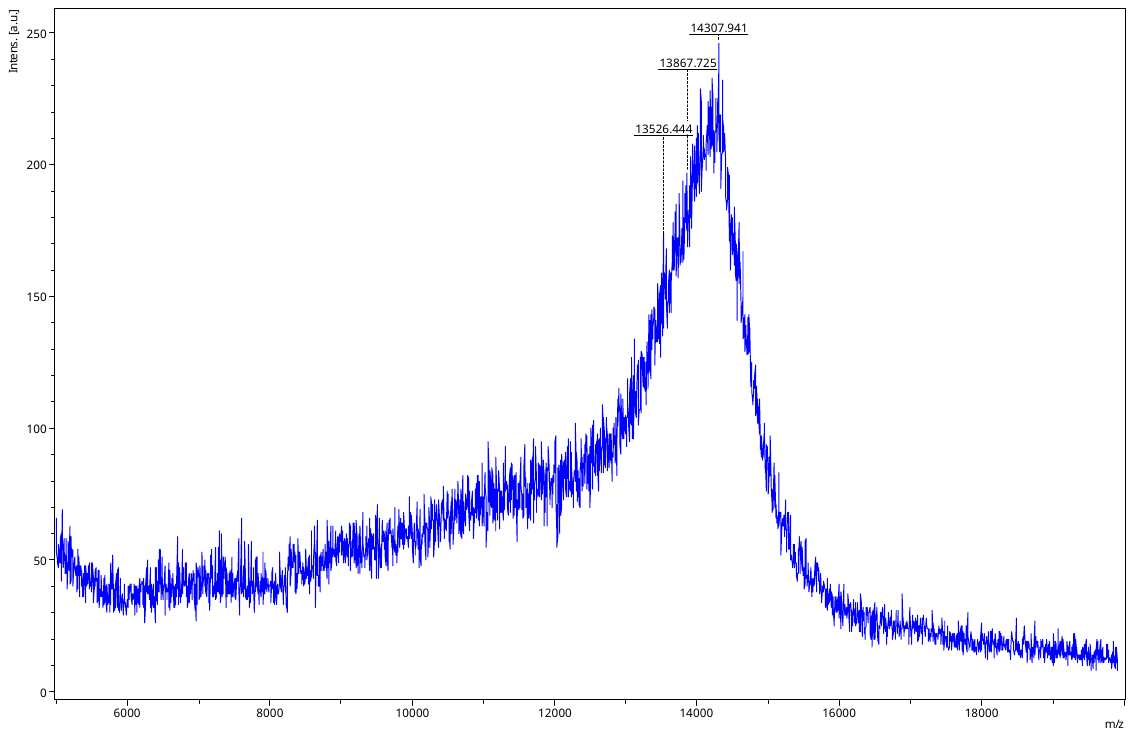
L10+MHP2

L11+MHPL11+MHP

L11+MHP

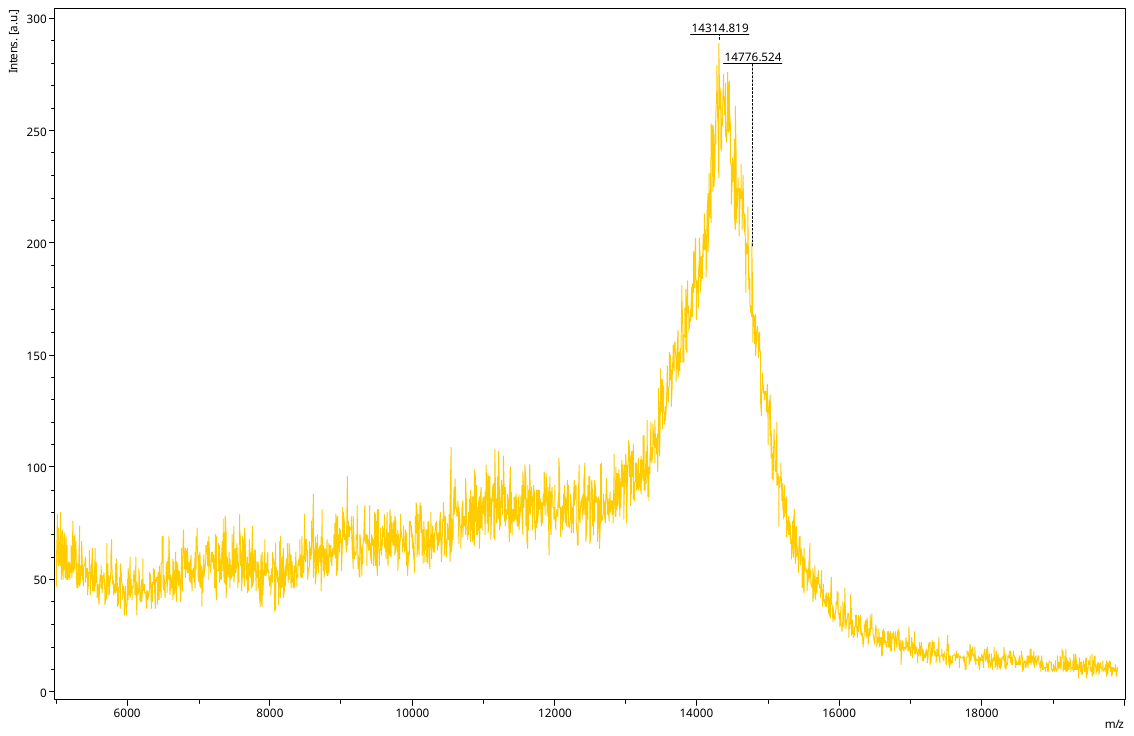

L12+MHP

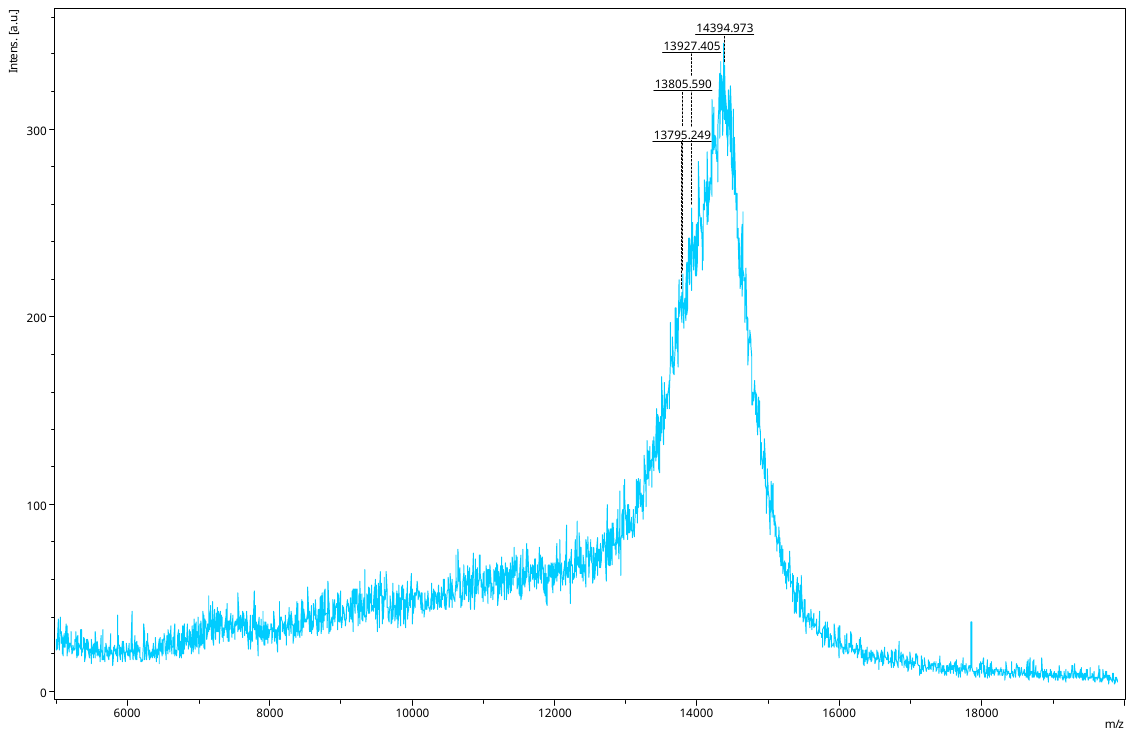

L13+MHP

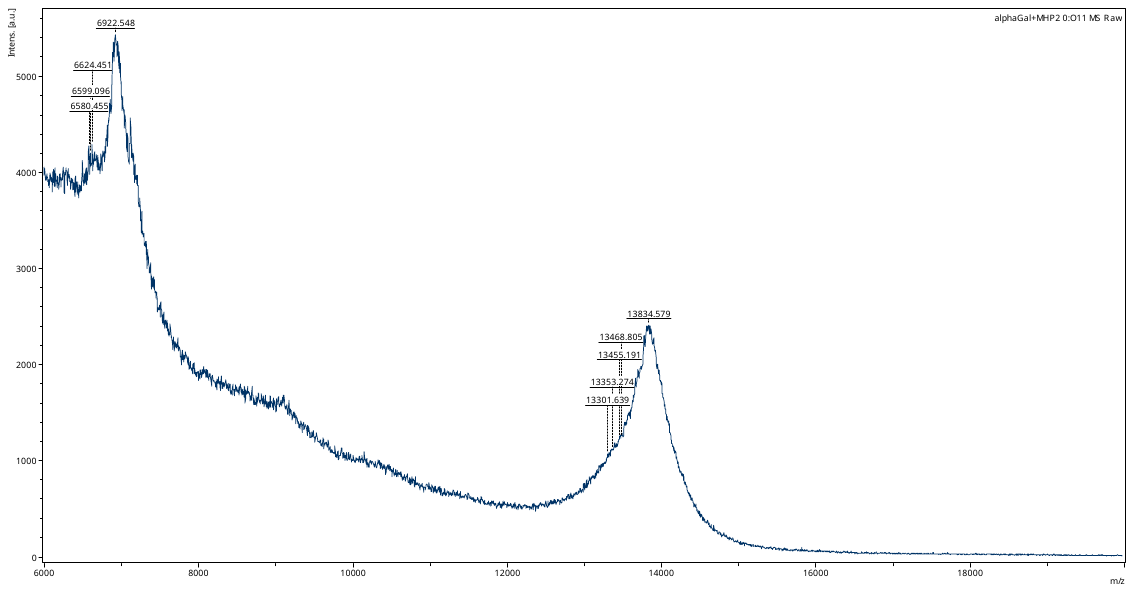

L14+MHP

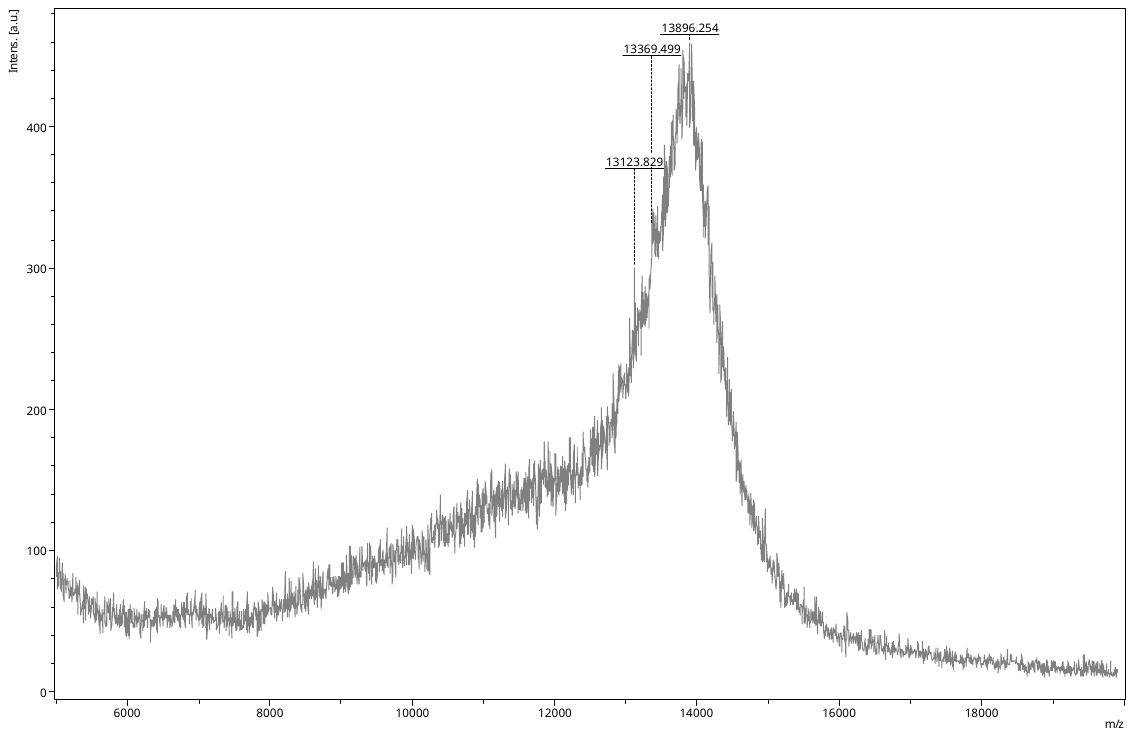

L15+MHP

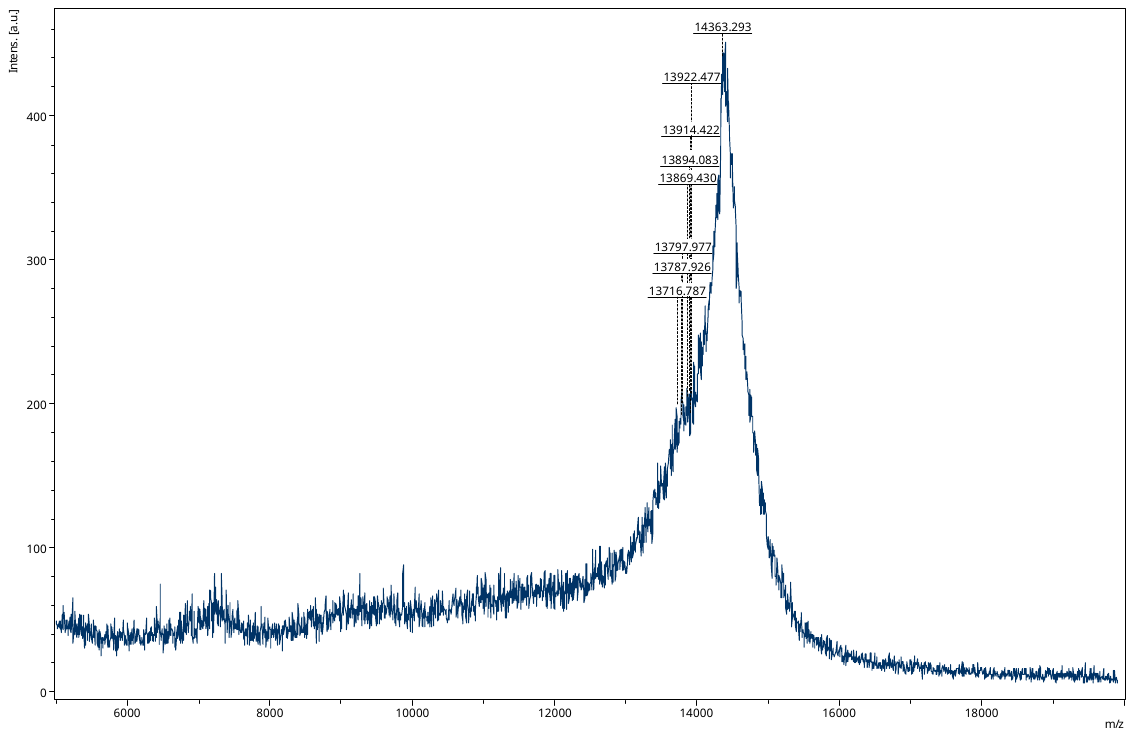

L16+MHP

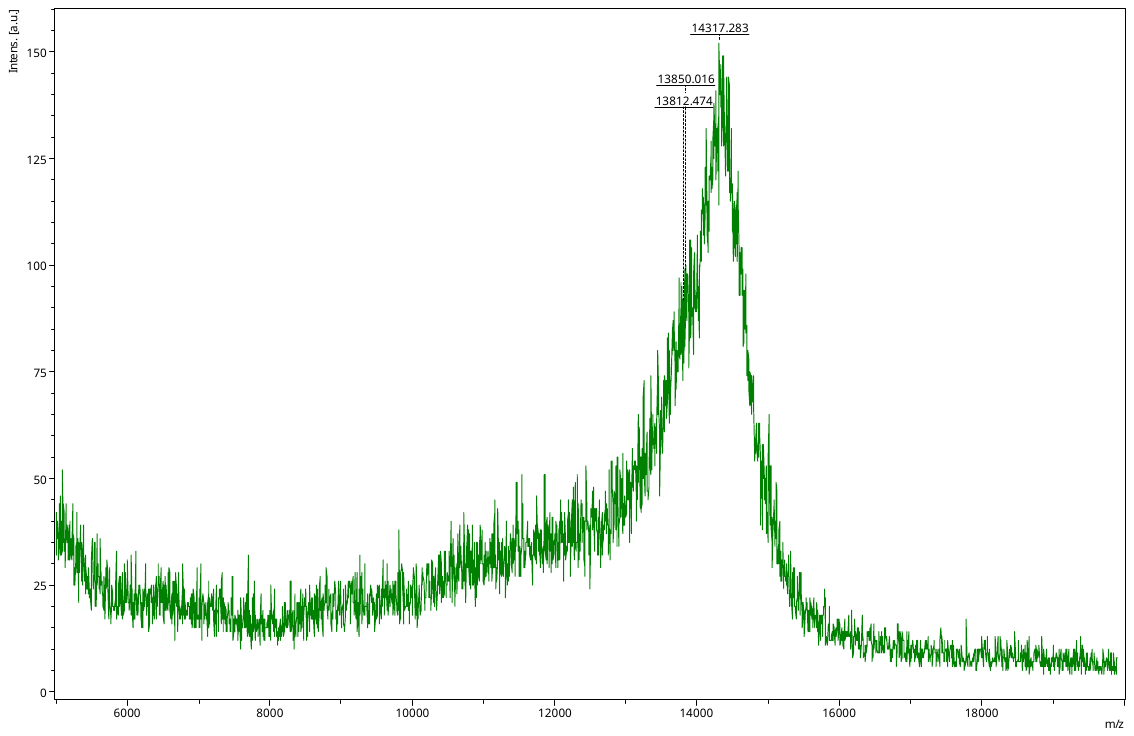

L17+MHP

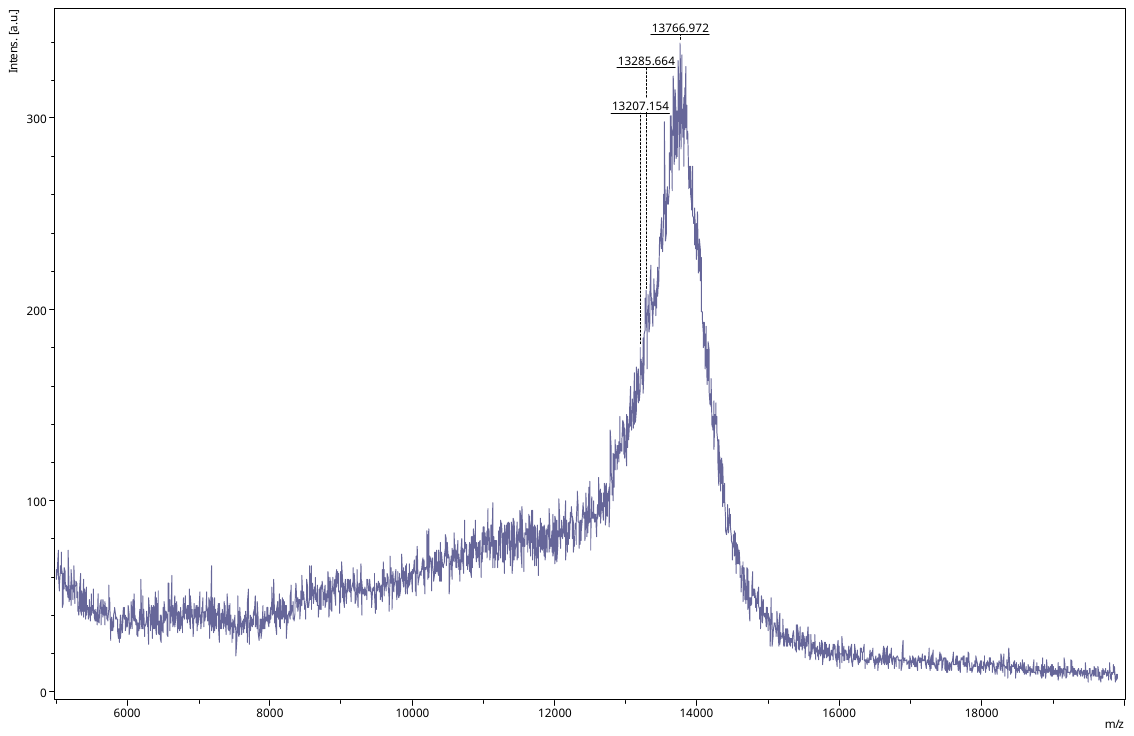

L18+MHP

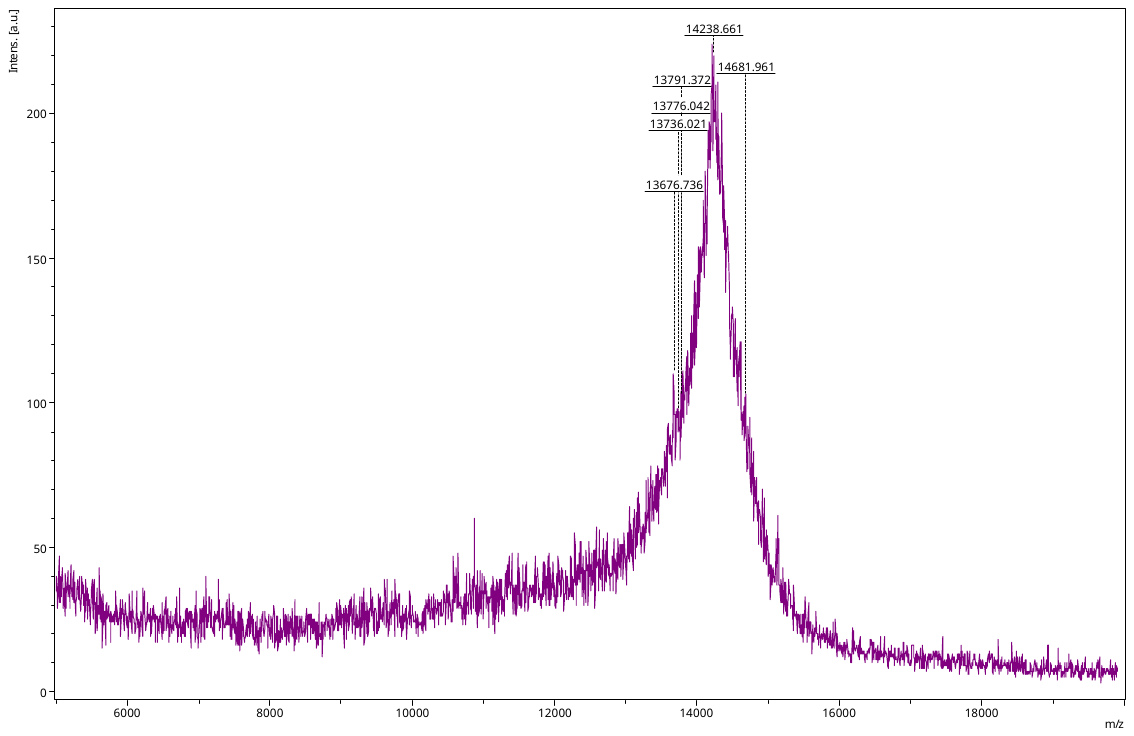

L19+MHP

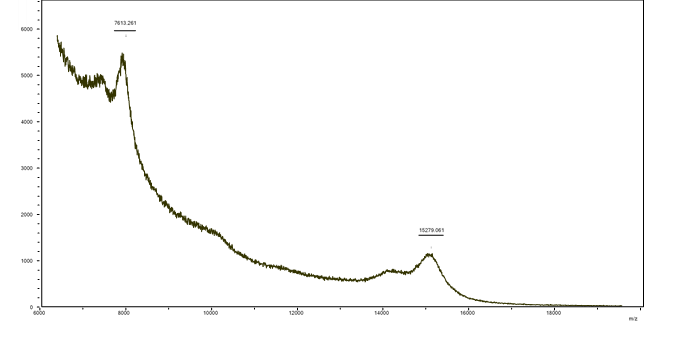

L20+MHP

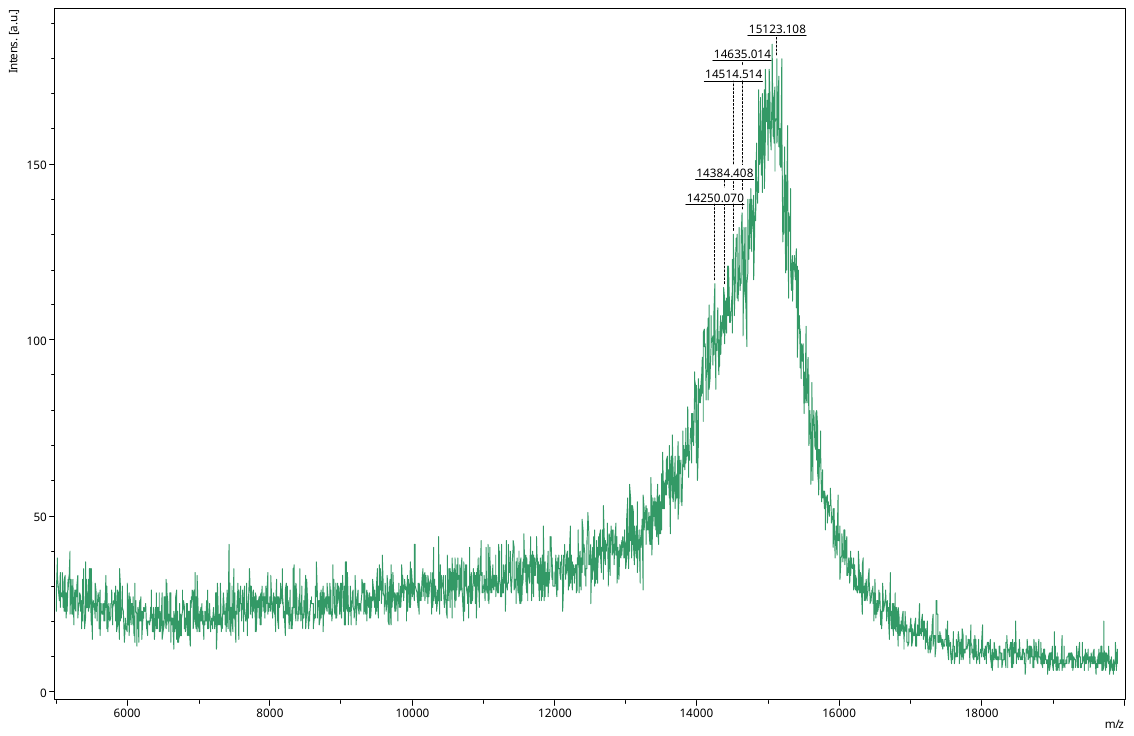

L21+MHP2

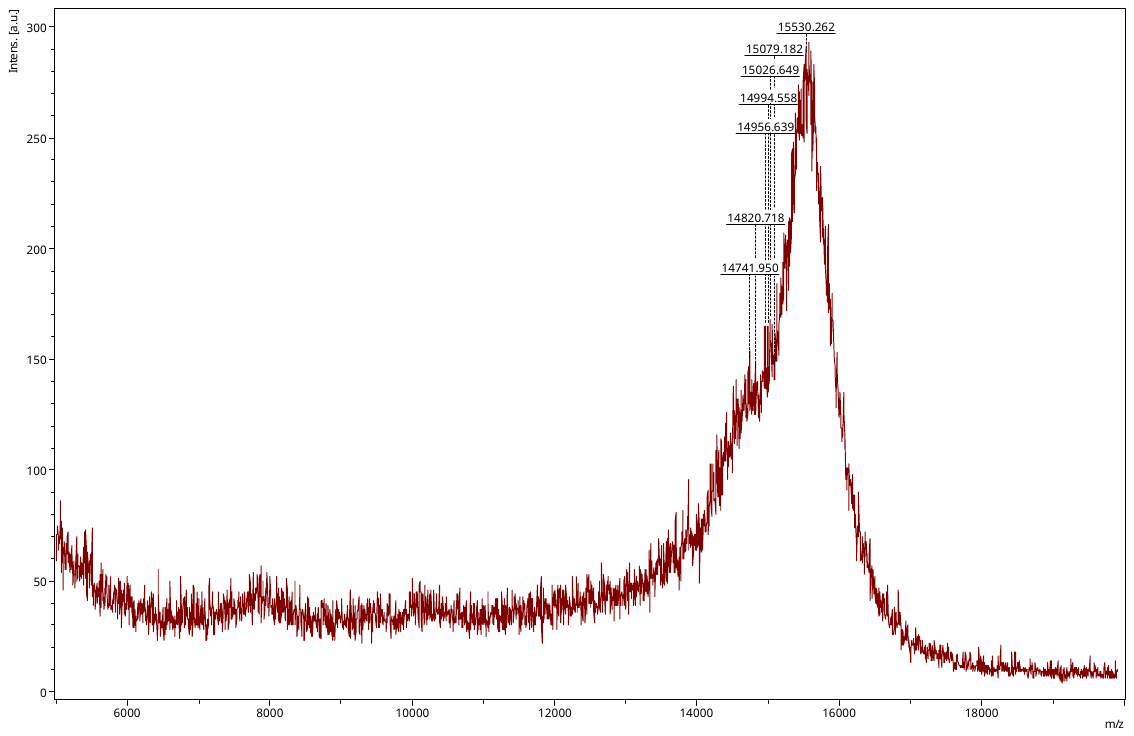

L22+MHP2

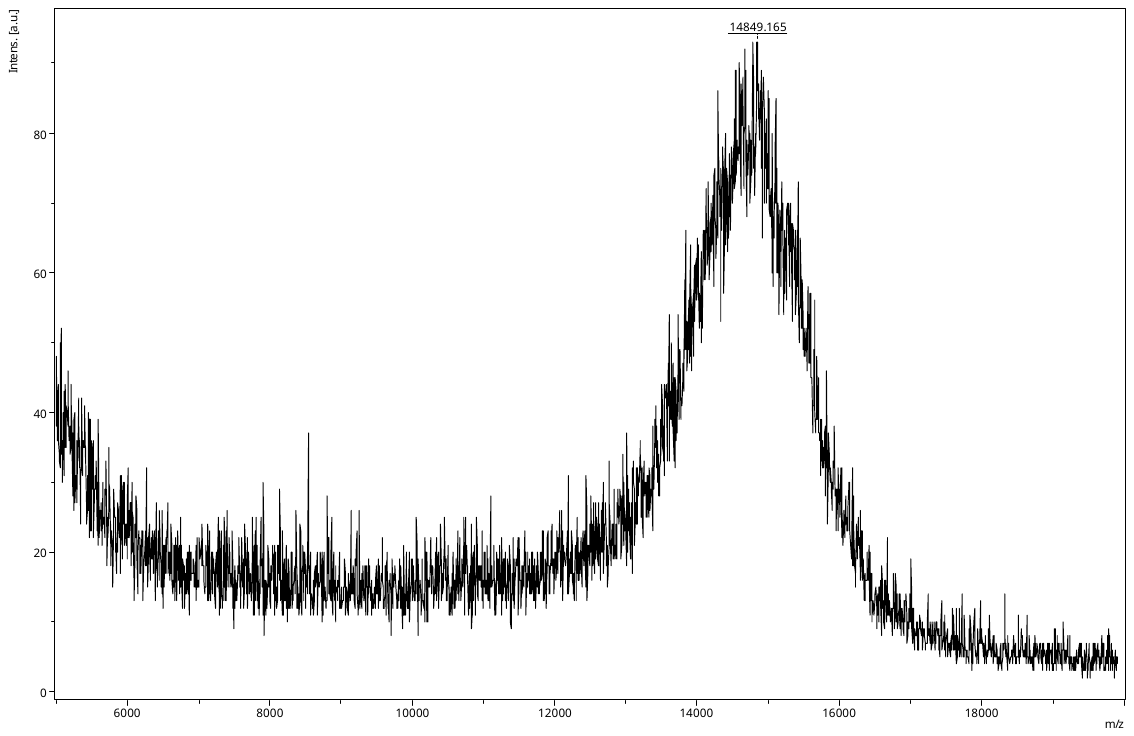

L31+MHP2

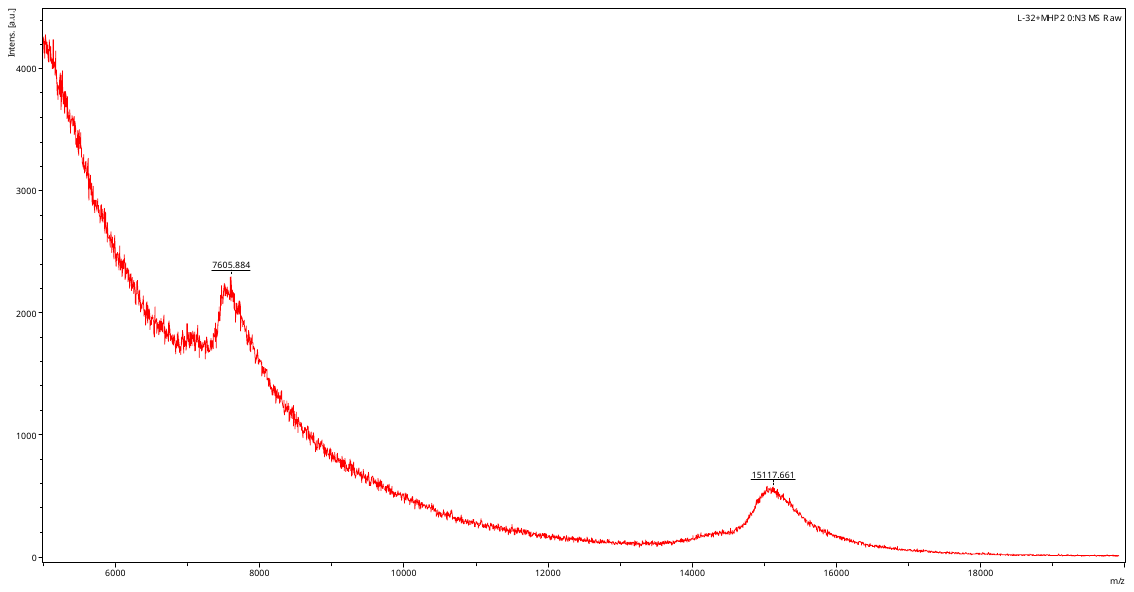

L32+MHP

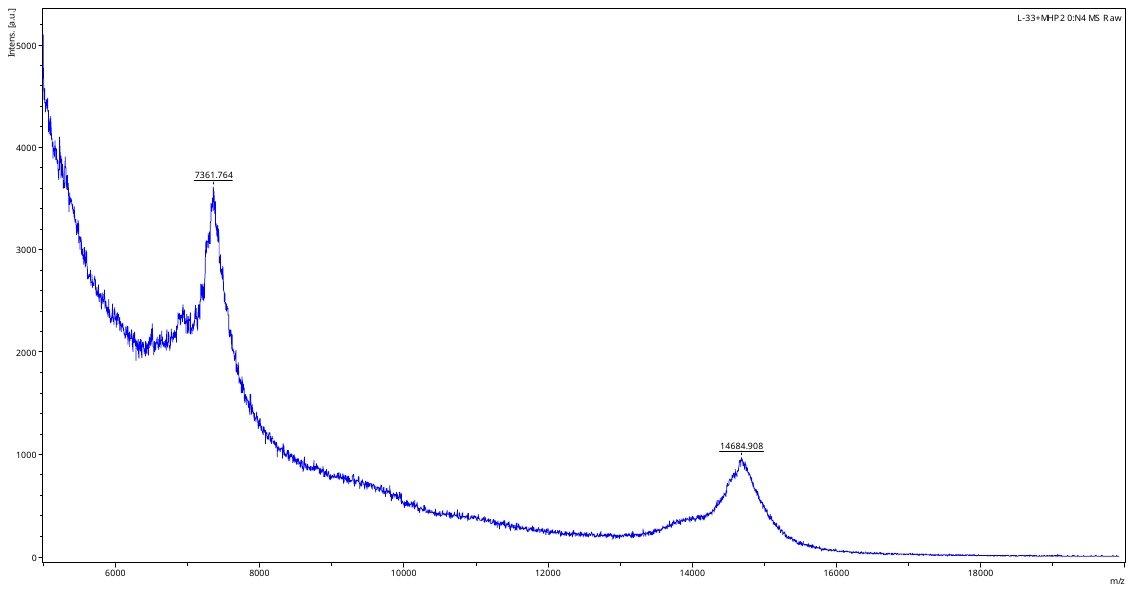

L33+MHP

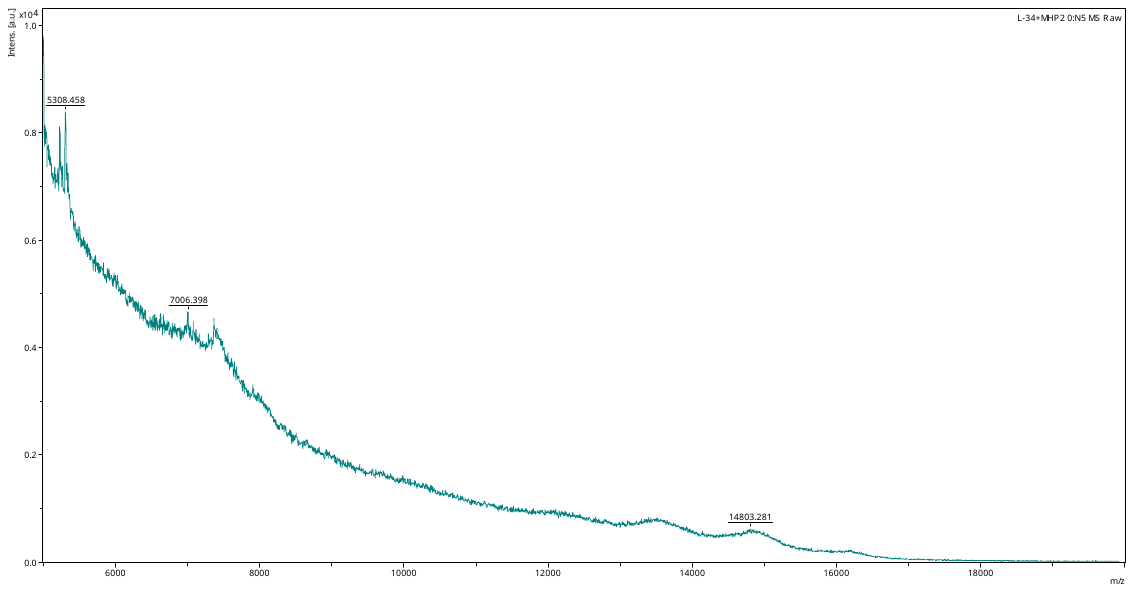

L34+MHP

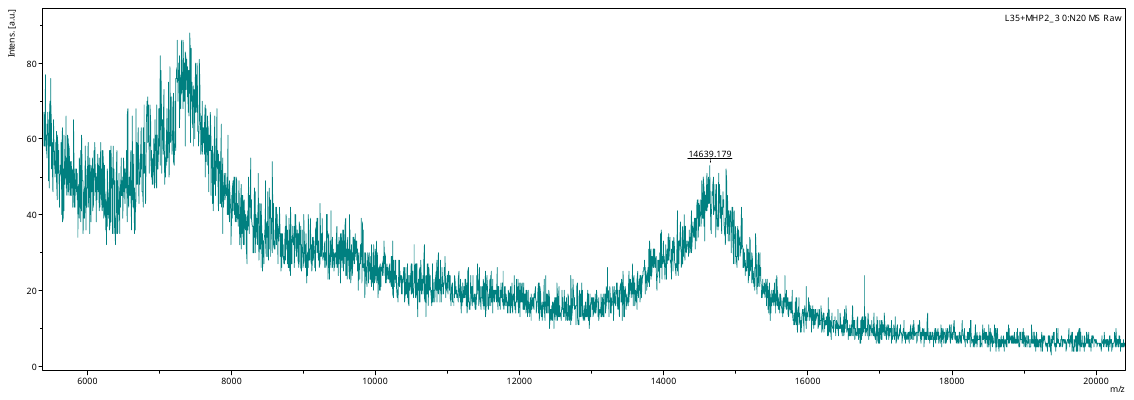

L35+MHP

L36+MHP

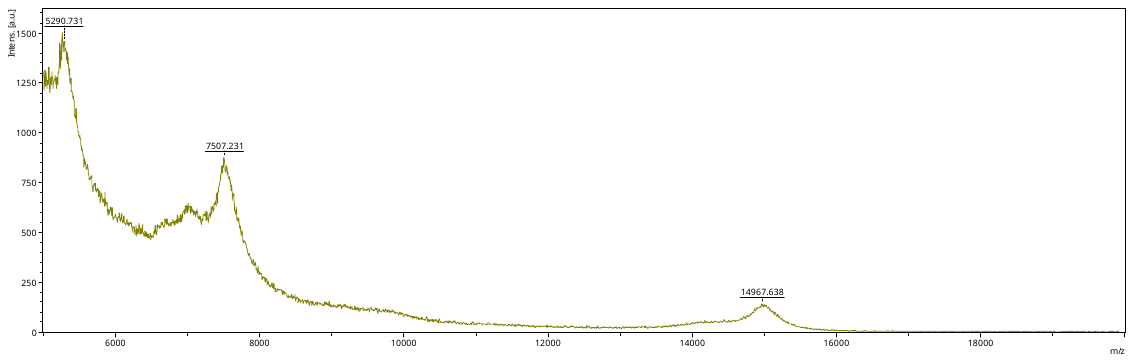

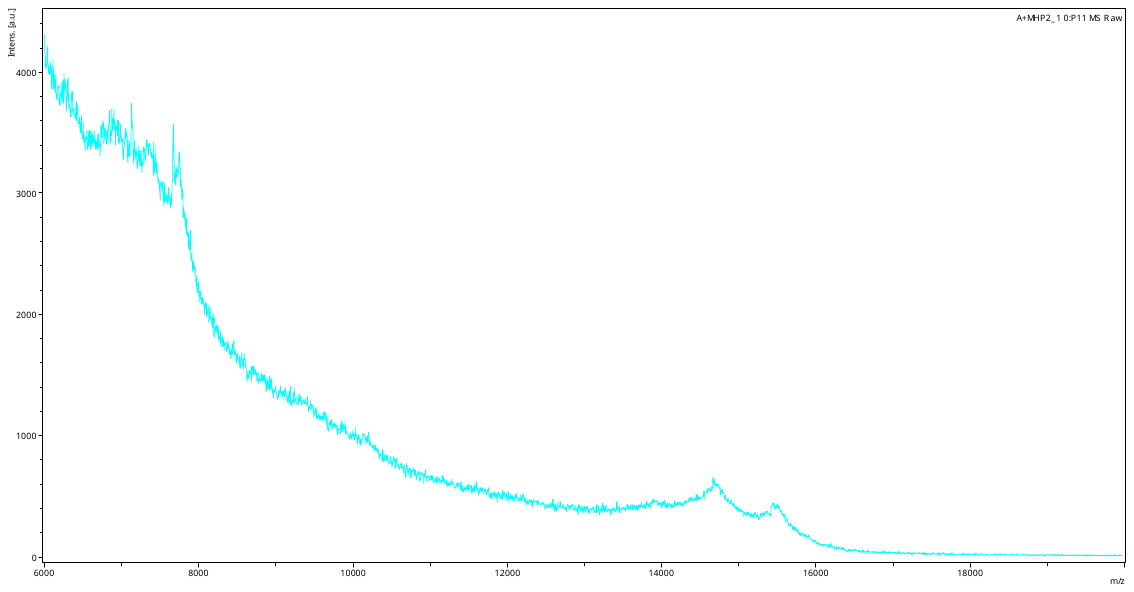

L37+MHP

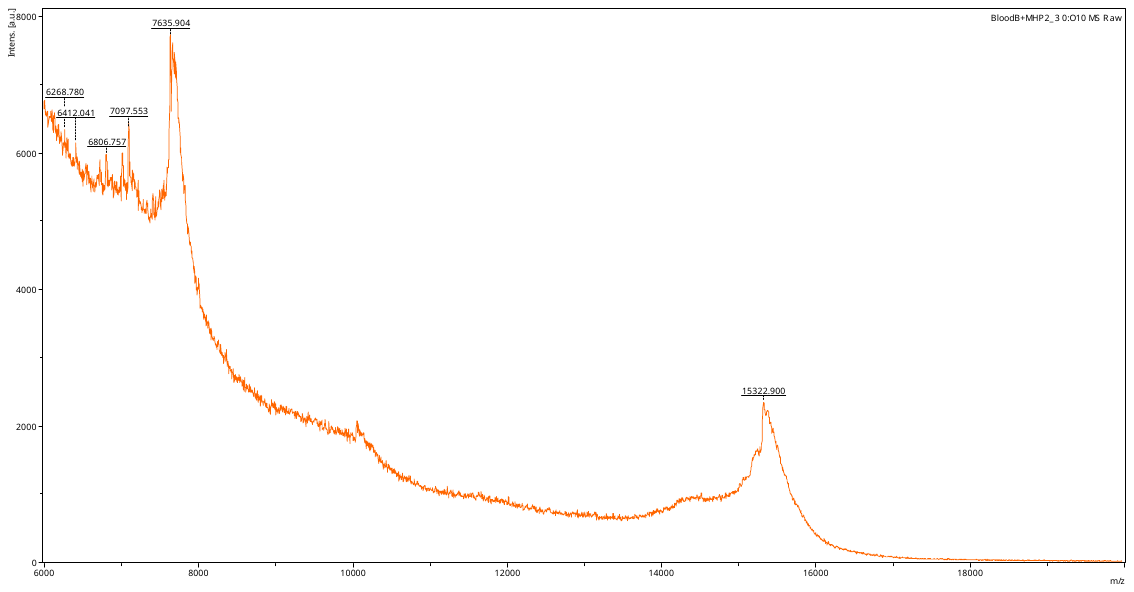

L38+MHP

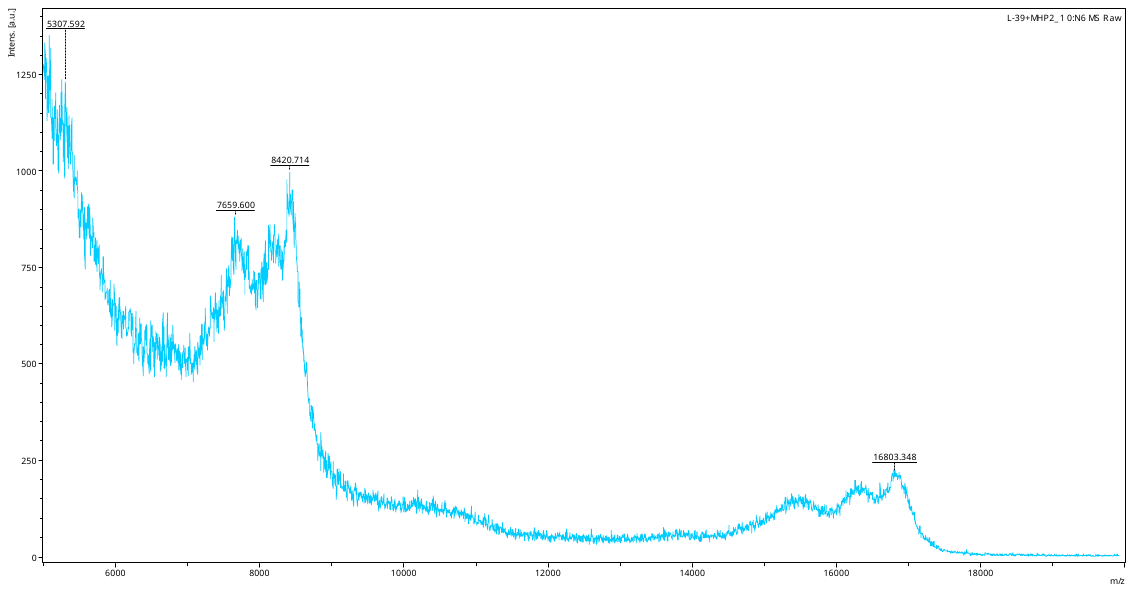

L39+MHP

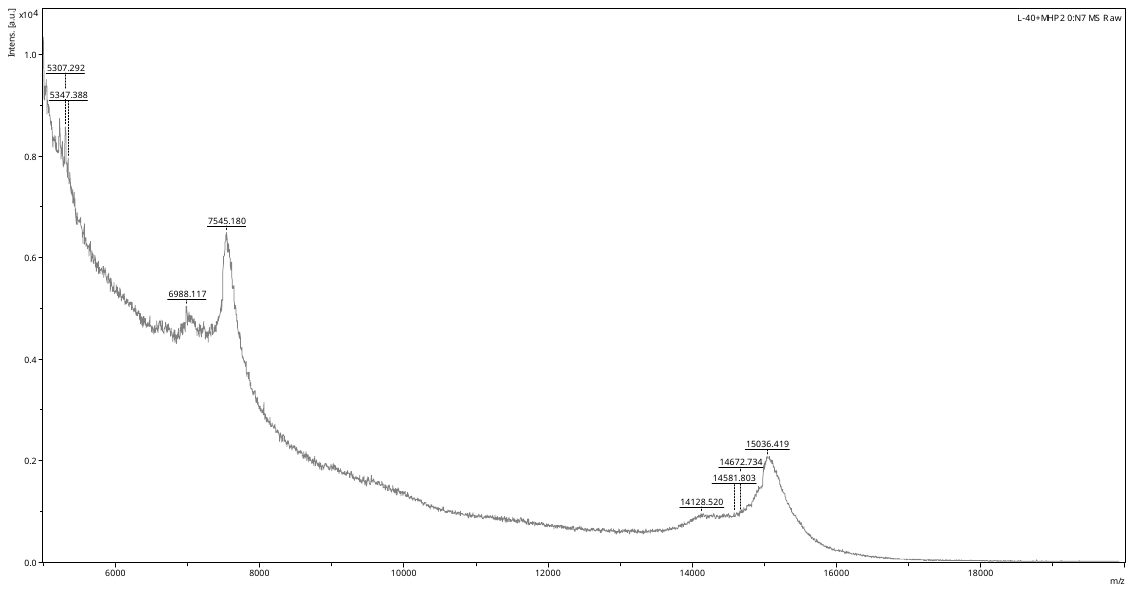

L40+MHP

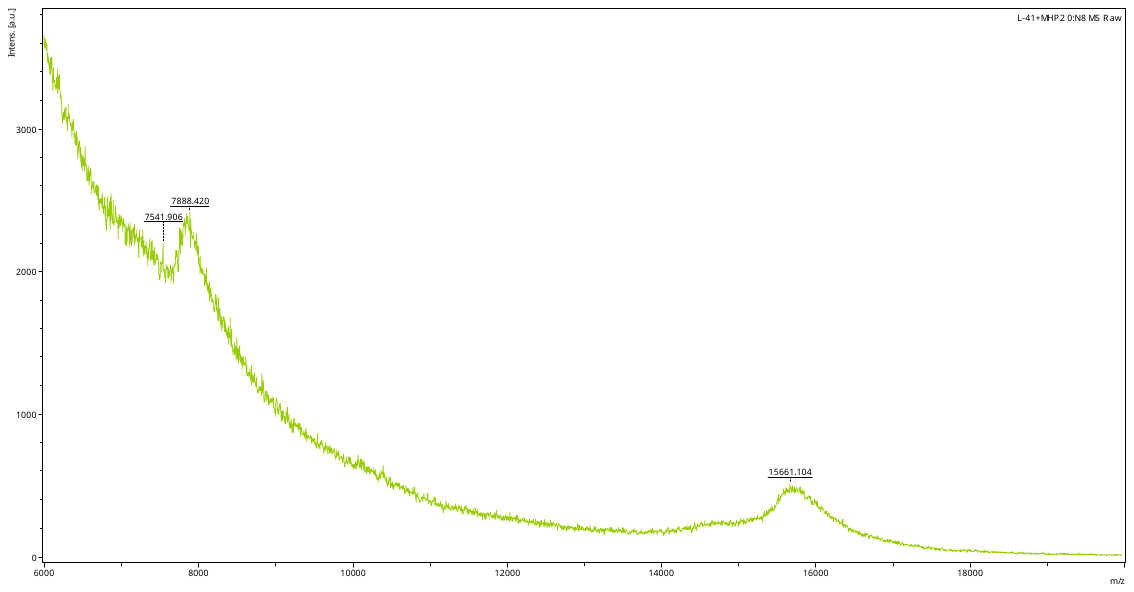

L41+MHP

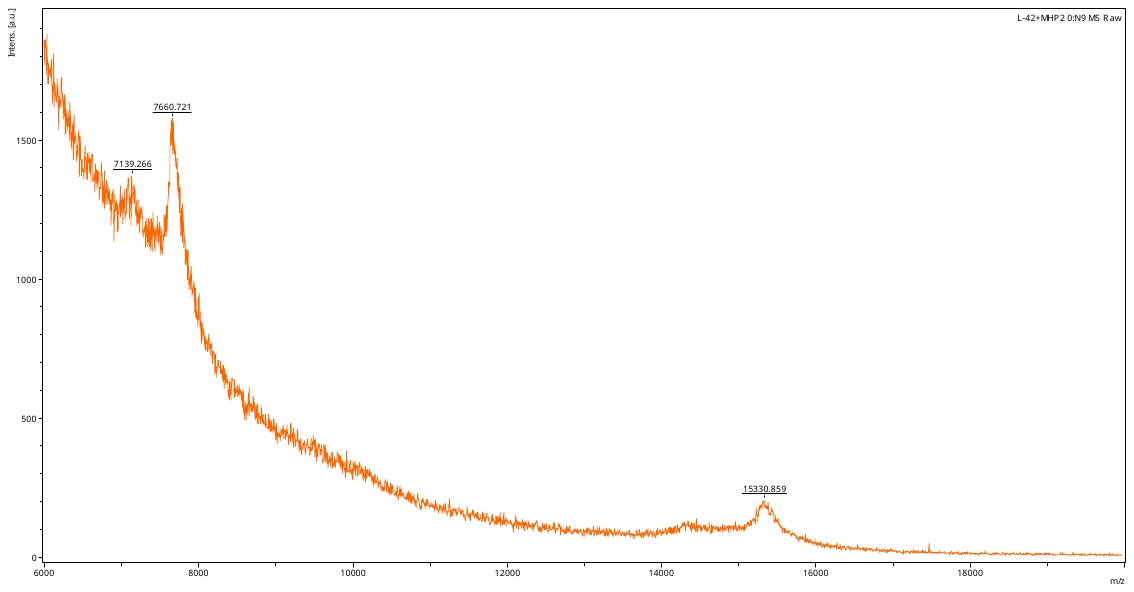

L42+MHP

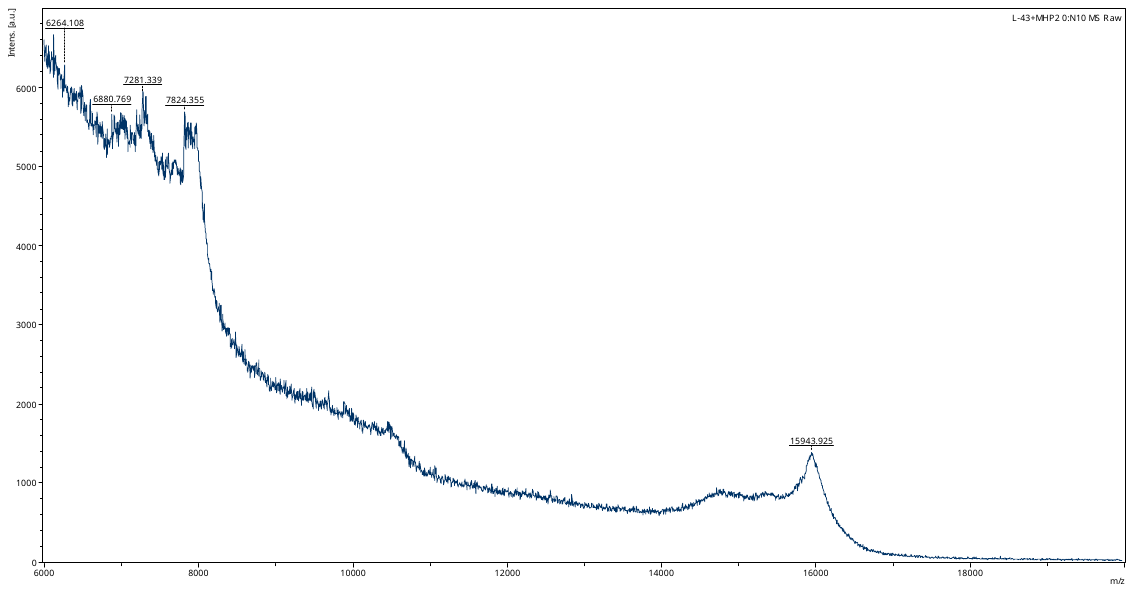

L43+MHP

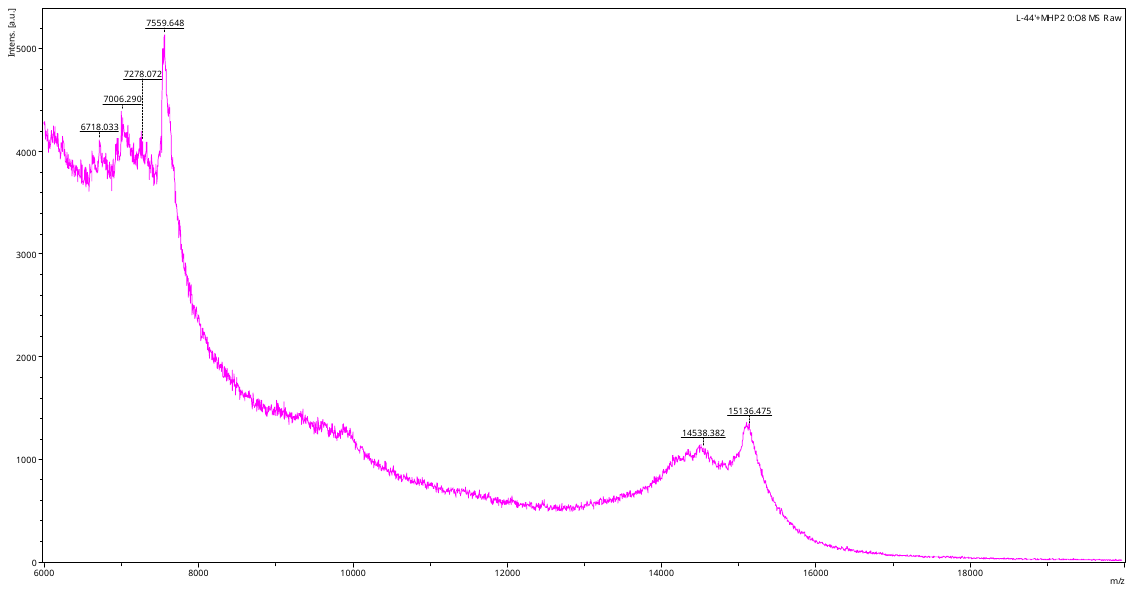

L44+MHP

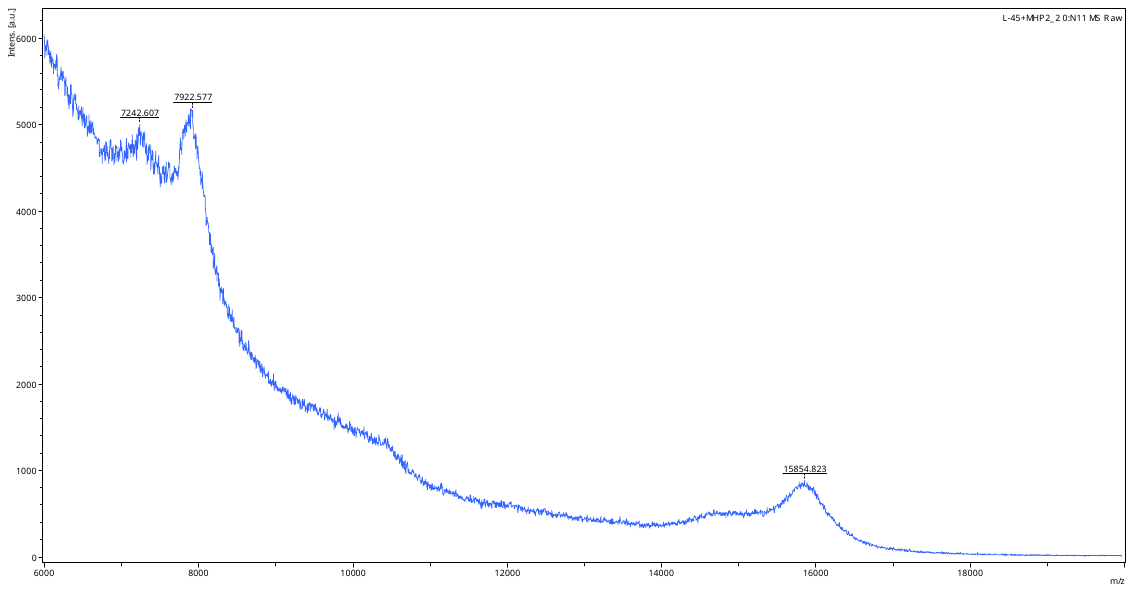

L45+MHP

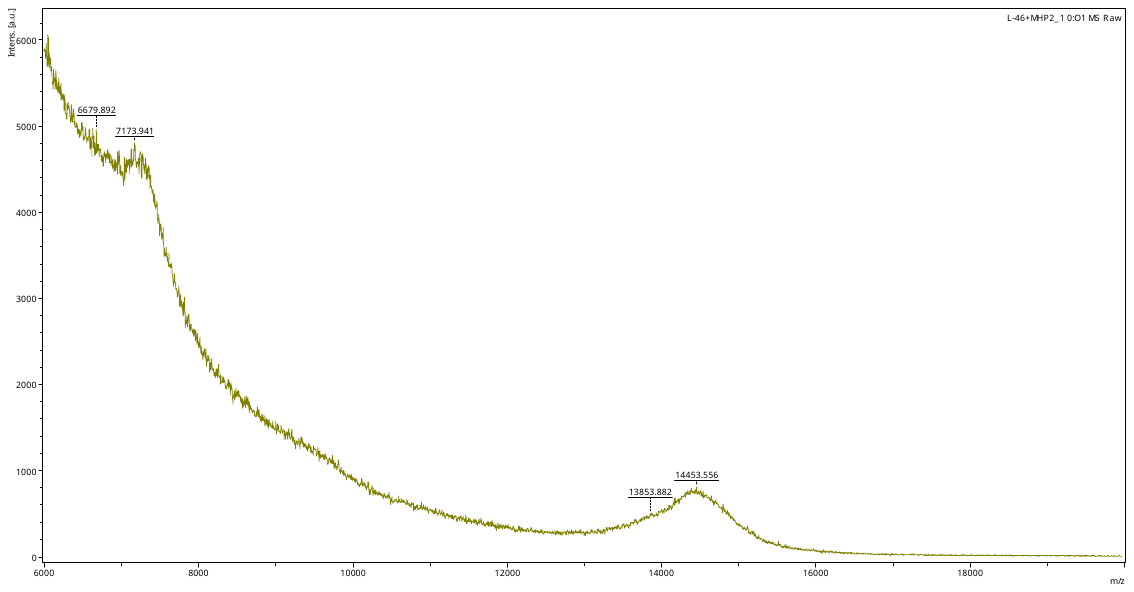

L46+MHP

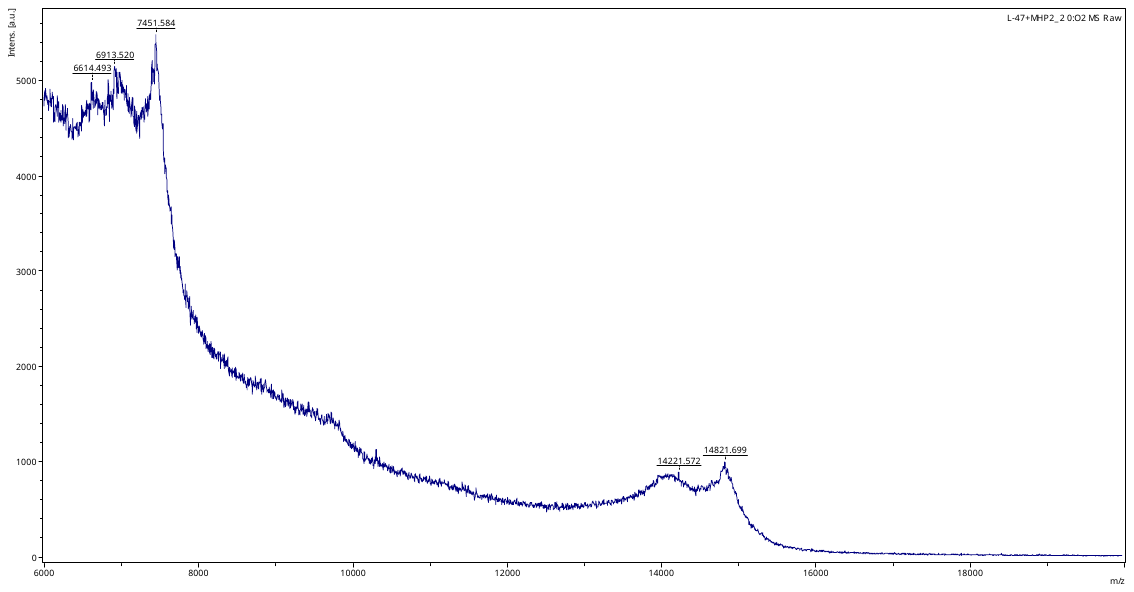

L47+MHP

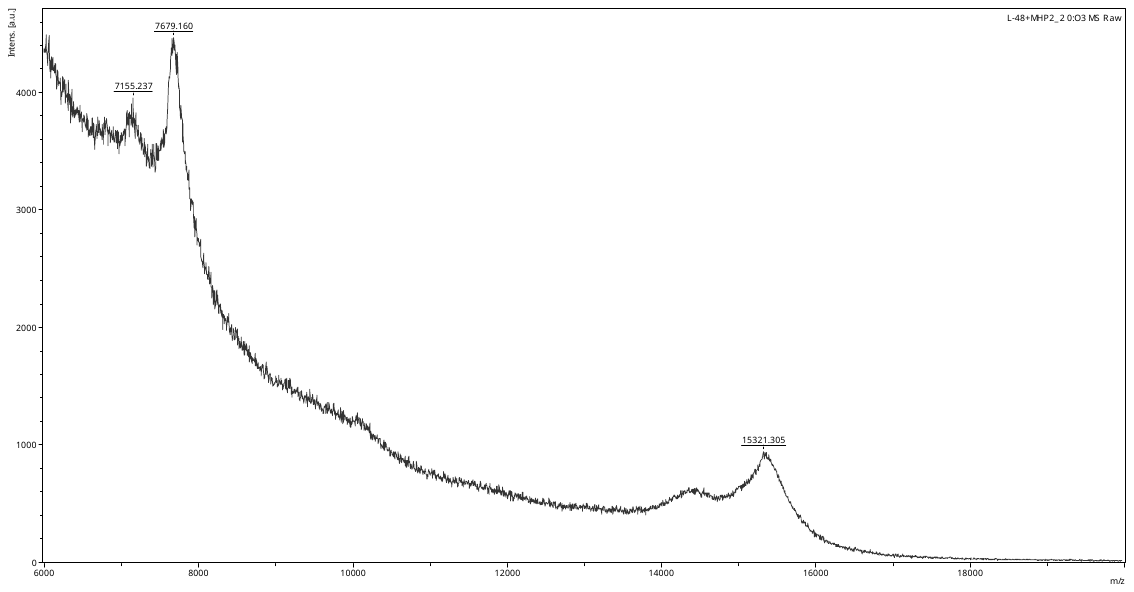

L48+MHP

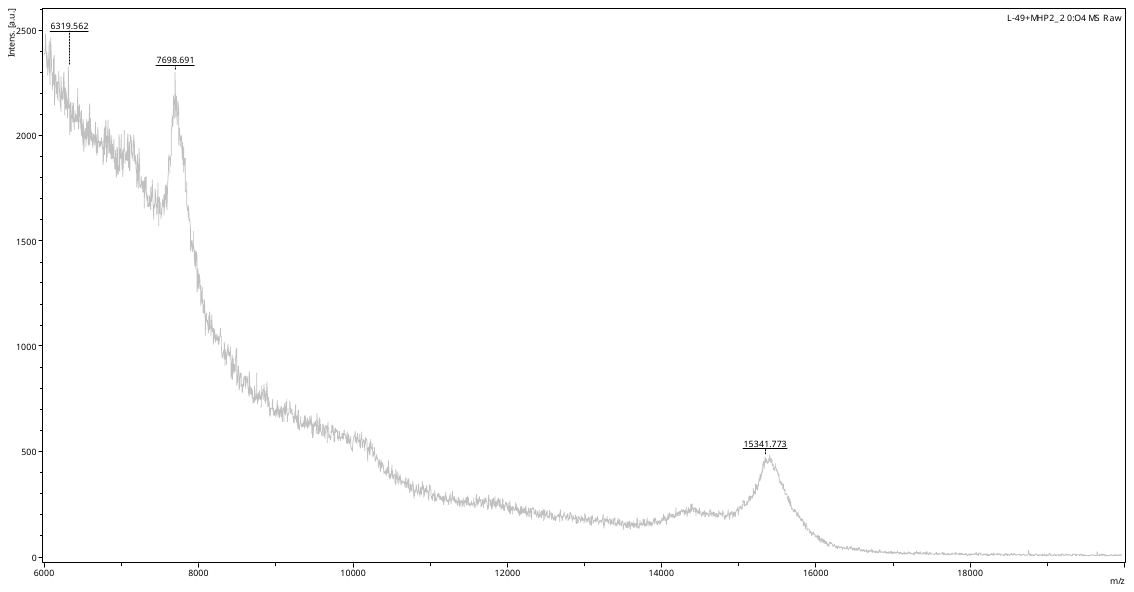

L49+MHP

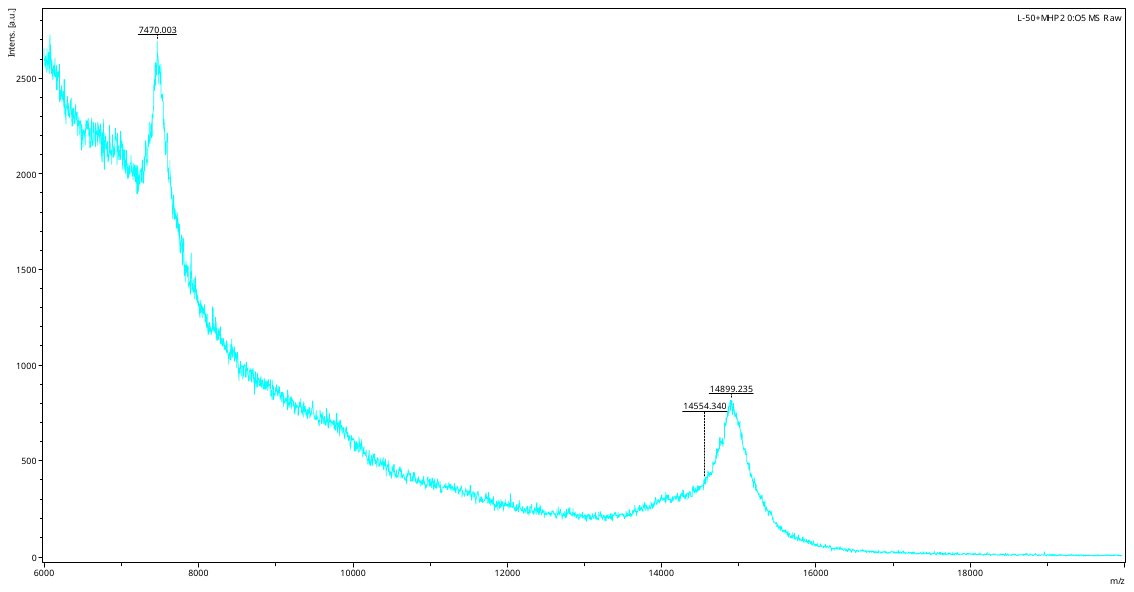

L50+MHP

#### HPLC traces of DNA and DNA glycan conjugates

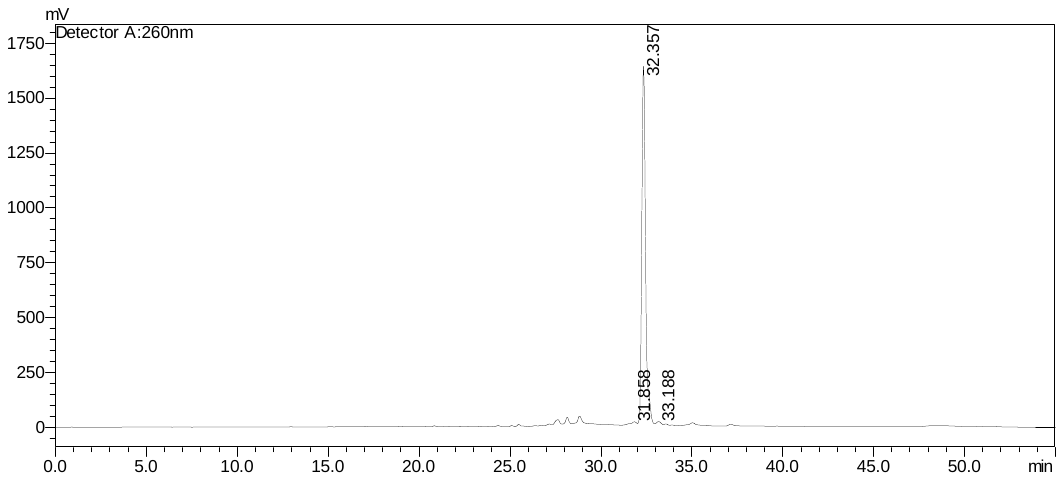

Headpiece DNA

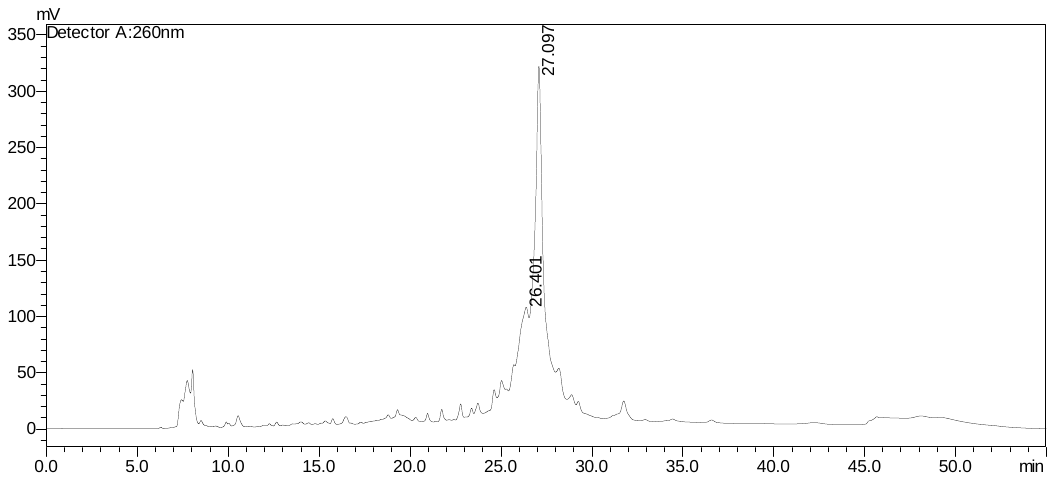

Headpiece DNA + GH

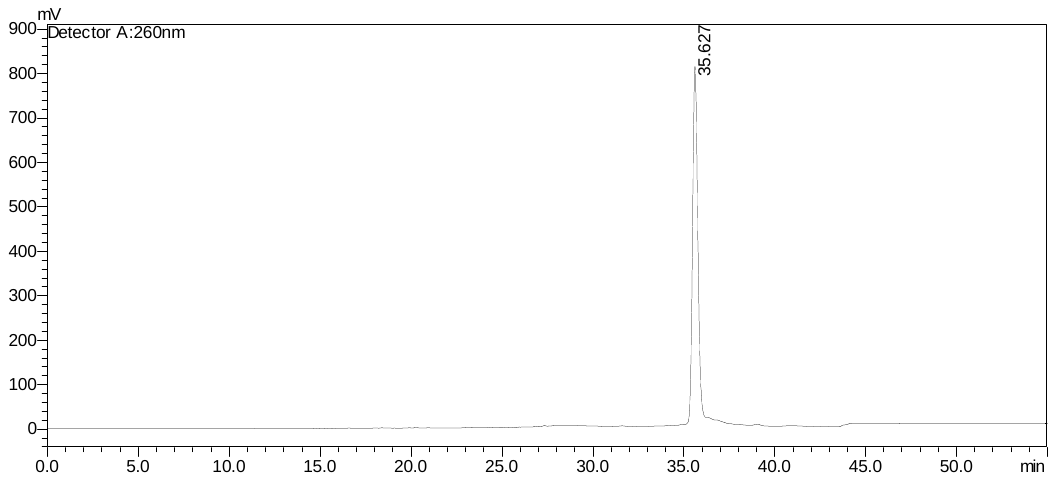

Multi Headpiece DNA

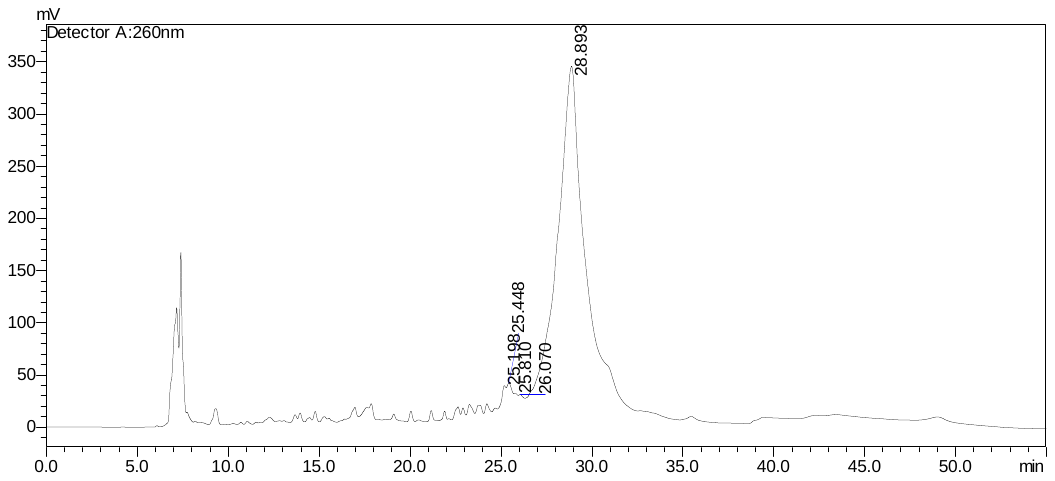

MHP+ GH

### 4. Tables

**Table S1.** **DNA sequences used in headpiece conjugation code ligation (HCCL) library synthesis**

| **Headpiece** | | | |
| --- | --- | --- | --- |
| **1** | Monovalent Headpiece | | /5Hexynyl/AAT GAT ACG GCG A |
| **2** | Multivalent Headpiece | | 5'- /5Phos/GAG TCG TCC TAC /i5OctdU/GC T/i5OctdU/T AC/i5OctdU/ TCG  AGG ACG ACT CAA TG -3' |
| **Tail DNA sequences** | | | |
| **1** | **GH** | /5Phos/CCACCGAAGAAAAGACGATCTACGCTCTACGAATAACGATGTAAGACACGTCTGAACTCCAGTCAC | |
| **2** | **Gb5** | /5Phos/CCACCGAACGATCTACGCTCTACGAATAACGATGTAAGACACGTCTGAACTCCAGTCAC | |
| **3** | **Gb4** | /5Phos/CCACCGAACGCTCTACGAATAACGATGTAAGACACGTCTGAACTCCAGTCAC | |
| **4** | **Gb3** | /5Phos/CCACCGAACGAATAACGATGTAAGACCCCAGTCAGGCCTAACGTACACGTCTGAACTCCAGTCAC | |
| **5** | **Gb2** | /5Phos/CCACCGAACGATGTAAGACCCCAGTCAGGCCTAACGTACACGTCTGAACTCCAGTCAC | |
| **6** | **Bb4** | /5Phos/CCACCGAAGAAAAGACGATCTACGCTCTACGACACGTCTGAACTCCAGTCAC | |
| **7** | **Bb3** | /5Phos/CCACCGAAGAAAAGACGATCTACGCCCCCAGTCAGGCCTAACGTACACGTCTGAACTCCAGTCAC | |
| **8** | **Bb2** | /5Phos/CCACCGAAGAAAAGACGACCCCAGTCAGGCCTAACGTACACGTCTGAACTCCAGTCAC | |
| **Reverse Compliment DNA** | | | |
| **1** | **A’** | GTGACTGGAGTTCAGACGTGTCTTACATCGTAGAGCTTAGATCGTTTTCTTTTCTAAAGTTGCGTTCGGTGGTCGCCGTATCATT | |
| **2** | **α-Gal’** | GTGACTGGAGTTCAGACGTGTCTTACATCGTAGAGCTTAGATCGTTTTCTTTTCTAAAGTTTCGTTCGGTGGTCGCCGTATCATT | |
| **3** | **GH’** | GTGACTGGAGTTCAGACGTGTCTTACATCGTTATTCGTAGAGCGTAGATCGTCTTTTCTTCGGTGGTCGCCGTATCATT | |
| **4** | **Gb5’** | GTGACTGGAGTTCAGACGTGTCTTACATCGTTATTCGTAGAGCGTAGATCGTTCGGTGGTCGCCGTATCATT | |
| **5** | **Gb4’** | GTGACTGGAGTTCAGACGTGTCTTACATCGTTATTCGTAGAGCGTTCGGTGGTCGCCGTATCATT | |
| **6** | **Gb3’** | GTGACTGGAGTTCAGACGTGTACGTTAGGCCTGACTGGGGTCTTACATCGTTATTCGTTCGGTGGTCGCCGTATCATT | |
| **7** | **Gb2’** | GTGACTGGAGTTCAGACGTGTACGTTAGGCCTGACTGGGGTCTTACATCGTTCGGTGGTCGCCGTATCATT | |
| **8** | **Bb4’** | GTGACTGGAGTTCAGACGTGTCGTAGAGCGTAGATCGTCTTTTCTTCGGTGGTCGCCGTATCATT | |
| **9** | **Bb3’** | GTGACTGGAGTTCAGACGTGTACGTTAGGCCTGACTGGGGGCGTAGATCGTCTTTTCTTCGGTGGTCGCCGTATCATT | |
| **10** | **Bb2’** | GTGACTGGAGTTCAGACGTGTACGTTAGGCCTGACTGGGGTCGTCTTTTCTTCGGTGGTCGCCGTATCATT | |

**Table S2.** MALDI Mass of DNA glycan conjugates

| **Glycan Conjugate** | **Glycan M.W.** | **MALDI Mass of conjugates** | **Number of Glycans per conjugate** |
| --- | --- | --- | --- |
| **Headpiece (HP)** |  | 4171.70 |  |
| **GH-HP** | 1099.01 | 5275.75 | 1 |
| **Gb5-HP** | 925.87 | 5127.45 | 1 |
| **Gb4-HP** | 790.73 | 4965.59 | 1 |
| **Gb3-HP** | 587.53 | 4761.35 | 1 |
| **Gb2-HP** | 425.39 | 4597.70 | 1 |
| **BB4-HP** | 774.73 | 4949.86 | 1 |
| **BB3-HP** | 612.59 | 4706.89 | 1 |
| **BB2-HP** | 409.39 | 4582.40 | 1 |
| **Multi headpiece (MHP)** |  | 11994.61 |  |
| **GH-MHP** | 1099.01 | 15157.51 | 3 |
| **Gb5-MHP** | 925.87 | 14863.08 | 3 |
| **Gb4-MHP** | 790.73 | 14384.12 | 3 |
| **Gb3-MHP** | 587.53 | 13771.39 | 3 |
| **Gb2-MHP** | 425.39 | 13318.42 | 3 |
| **BB4-MHP** | 774.73 | 14332.77 | 3 |
| **BB3-MHP** | 612.59 | 13836.22 | 3 |
| **BB2-MHP** | 409.39 | 13243.05 | 3 |
| **A +MHP** | 1140.06 | 15496.54 | 3 |
| **aGal+MHP** | 587.53 | 13778.81 | 3 |

**Table S3:** DNA Sequences used for the synthesis of DEGL-50

| **Glycan ID** | **Sequence** |
| --- | --- |
| L1 | 5’pho/atacggcgaccaccgaacgctctacgatgtaagacaacacgtctgaactccagtcac  ttactatgccgctggtggcttgcgagatgctacattctgttgtgcagacttgaggtcagtg |
| L2 | 5’pho/atacggcgaccaccgaaggcggtacgatgtaagacaacacgtctgaactccagtcac  ttactatgccgctggtggcttccgccatgctacattctgttgtgcagacttgaggtcagtg |
| L3 | 5’pho/atacggcgaccaccgaaggcgttacgatgtaagacaacacgtctgaactccagtcac  ttactatgccgctggtggcttccgcaatgctacattctgttgtgcagacttgaggtcagtg |
| L4 | 5’pho/atacggcgaccaccgaaggcggttttacgctgtaaaacgatgtaagacaacacgtctgaactccagtcac  ttactatgccgctggtggcttccgccaaaatgcgacattttgctacattctgttgtgcagacttgaggtcagtg |
| L5 | 5’pho/atacggcgaccaccgaaggcggttttacgatctacgctgtaaaacgatgtaagacaacacgtctgaactccagtcac  ttactatgccgctggtggcttccgccaaaatgctagatgcgacattttgctacattctgttgtgcagacttgaggtcagtg |
| L6 | 5’pho/atacggcgaccaccgaaggcgataggcggtacgatgtaagacaacacgtctgaactccagtcac  ttactatgccgctggtggcttccgctatccgccatgctacattctgttgtgcagacttgaggtcagtg |
| L7 | 5’pho/atacggcgaccaccgaaggcgataggcgttacgatgtaagacaacacgtctgaactccagtcac  ttactatgccgctggtggcttccgctatccgcaatgctacattctgttgtgcagacttgaggtcagtg |
| L8 | 5’pho/atacggcgaccaccgaaggcgataggcggttttacgctgtaaaacgatgtaagacaacacgtctgaactccagtcac  ttactatgccgctggtggcttccgctatccgccaaaatgcgacattttgctacattctgttgtgcagacttgaggtcagtg |
| L9 | 5’pho/atacggcgaccaccgaaggcgataggcggttttacgatctacgctgtaaaacgatgtaagacaacacgtctgaactccagtcac  ttactatgccgctggtggcttccgctatccgccaaaatgctagatgcgacattttgctacattctgttgtgcagacttgaggtcagtg |
| L10 | 5’pho/atacggcgaccaccgaaggtggtacgatgtaagacaacacgtctgaactccagtcac  ttactatgccgctggtggcttccaccatgctacattctgttgtgcagacttgaggtcagtg |
| L11 | 5’pho/atacggcgaccaccgaaggtgttacgatgtaagacaacacgtctgaactccagtcac  ttactatgccgctggtggcttccacaatgctacattctgttgtgcagacttgaggtcagtg |
| L12 | 5’pho/atacggcgaccaccgaagggggtacgatgtaagacaacacgtctgaactccagtcac  ttactatgccgctggtggcttcccccatgctacattctgttgtgcagacttgaggtcagtg |
| L13 | 5’pho/atacggcgaccaccgaaggggttacgatgtaagacaacacgtctgaactccagtcac  ttactatgccgctggtggcttccccaatgctacattctgttgtgcagacttgaggtcagtg |
| L14 | 5’pho/atacggcgaccaccgaacgaaacacgatgtaagcccccagtcaggcctaacgtacacgtctgaactccagtcac  ttactattatgccgctggtggcttgctttgtgctacattcgggggtcagtccggattgcatgtgcagacttgaggtcagtg |
| L15 | 5’pho/atacgg cgaccaccgaagaaaagacgatgtaagacaacac gtctgaactccagtcac  ttactatgccgctggtggcttcttttctgctacattctgttgtgcagacttgaggtcagtg |
| L16 | 5’pho/atacggcgaccaccgaacgcaactttagaaaagaaaacgatgtaagacaacacgtctgaactccagtcac  ttactatgccgctggtggcttgcgttgaaatcttttcttttgctacattctgttgtgcagacttgaggtcagtg |
| L17 | 5’pho/atacggcgaccaccgaacgaaactttagaaaagaaaacgatgtaagacaacacgtctgaactccagtcac  ttactatgccgctggtggcttgctttgaaatcttttcttttgctacattctgttgtgcagacttgaggtcagtg |
| L18 | 5’pho/atacggcgaccaccgaacgatgttttagaaaacaaaaagacaacacgtctgaactccagtcac  ttactatgccgctggtggcttgctacaaaatcttttgtttttctgttgtgcagacttgaggtcagtg |
| L19 | 5’pho/atacggcgaccaccgaagaaaagacgatgttttagaaaacaaaaagacaacacgtctgaactccagtcac  ttactatgccgctggtggcttcttttctgctacaaaatcttttgtttttctgttgtgcagacttgaggtcagtg |
| L20 | 5’pho/atacggcgaccaccgaaagctctacgatgtaagctcttttaggcgttaaaacgatgtaagacaacacgtctgaactccagtcac  ttactatgccgctggtggctttcgagatgctacattcgagaaaatccgcaattttgctacattctgttgtgcagacttgaggtcagtg |
| L21 | 5’pho/atacggcgaccaccgaagggggttttacgctgtaaaacgatgtaagacaacacgtctgaactccagtcac  ttactatgccgctggtggcttcccccaaaatgcgacattttgctacattctgttgtgcagacttgaggtcagtg |
| L22 | 5’pho/atacggcgaccaccgaagggggttttacgatctacgctgtaaaacgatgtaagacaacacgtctgaactccagtcac  ttactatgccgctggtggcttcccccaaaatgctagatgcgacattttgctacattctgttgtgcagacttgaggtcagtg |
| L23 | 5’pho/atacggcgaccaccgaaacgatgtaagaccccagtcaggcctaacgtacacgtctgaactccagtcac  ttactatgccgctggtggcttgctacattctggggtcagtccggattgcatgtgcagacttgaggtcagtg |
| L24 | 5’pho/atacggcgaccaccgaaacgaataacgatgtaagaccccagtcaggcctaacgtacacgtctgaactccagtcac  ttactatgccgctggtggcttgcttattgctacattctggggtcagtccggattgcatgtgcagacttgaggtcagtg |
| L25 | 5’pho/atacggcgaccaccgaacgctctacgaataacgatgtaagacacgtctgaactccagtcac  ttactatgccgctggtggcttgcgagatgcttattgctacattctgtgcagacttgaggtcagtg |
| L26 | 5’pho/atacgg cgaccaccgaaacgatctacgctctacgaataacgatgtaagacacgtctgaactccagtcac  ttactatgccgctggtggcttgctagatgcgagatgcttattgctacattctgtgcagacttgaggtcagtg |
| L27 | 5’pho/atacggcgaccaccgaaagaaaagacgatctacgctctacgaataacgatgtaagacacgtctgaactccagtcac  ttactatgccgctggtggcttcttttctgctagatgcgagatgcttattgctacattctgtgcagacttgaggtcagtg |
| L28 | 5’pho/atacgg cgaccaccgaaagaaaagacgatctacgctctacgacac gtctgaactccagtcac  ttactatgccgctggtggcttcttttctgctagatgcgagatgctgtgcagacttgaggtcagtg |
| L29 | 5’pho/atacggcgaccaccgaaagaaaagacgatctacgcccccagtcaggcctaacgtacacgtctgaactccagtcac  ttactatgccgctggtggcttcttttctgctagatgcgggggtcagtccggattgcatgtgcagacttgaggtcagtg |
| L30 | 5’pho/atacggcgaccaccgaaagaaaagacgaccccagtcaggcctaacgtacacgtctgaactccagtcac  ttactatgccgctggtggcttcttttctgctggggtcagtccggattgcatgtgcagacttgaggtcagtg |
| L31 | 5’pho/atacggcgaccaccgaaggcgataggcgataggcggttttacgctgtaaaacgatgtaagacaacacgtctgaactccagtcac  ttactatgccgctggtggcttccgctatccgctatccgccaaaatgcgacattttgctacattctgttgtgcagacttgaggtcagtg |
| L32 | 5’pho/atacggcgaccaccgaacgcaactttagaaaagaaaacgatgttttagaaaacaaaaagacaacacgtctgaactccagtcac  ttactatgccgctggtggcttgcgttgaaatcttttcttttgctacaaaatcttttgtttttctgttgtgcagacttgaggtcagtg |
| L33 | 5’pho/atacggcgaccaccgaacgaaactttagaaaagaaaacgatgttttagaaaacaaaaagacaacacgtctgaactccagtcac  ttactatgccgctggtggcttgctttgaaatcttttcttttgctacaaaatcttttgtttttctgttgtgcagacttgaggtcagtg |
| L34 | 5’pho/atacggcgaccaccgaaggcggttttaggcggtacgatctacgctgtaaaacgatgtaagacaacacgtctgaactccagtcac  ttactatgccgctggtggcttccgccaaaatccgccatgctagatgcgacattttgctacattctgttgtgcagacttgaggtcagtg |
| L35 | 5’pho/atacggcgaccaccga acgatgtaagctctacgatgtaagacaacacgtctgaactccagtcac  ttactatgccgctggtggcttgctacattcgagatgctacattctgttgtgcagacttgaggtcagtg |
| L36 | 5’pho/atacgg cgaccaccgaagaaaagacgatctaagctctacgatgtaagacacgtctgaactccagtcac  ttactatgccgctggtggcttcttttctgctagattcgagatgctacattctgtgcagacttgaggtcagtg |
| L37 | 5’pho/atacggcgaccaccgaacgcaactttagaaaagaaaacgatctaagatctacgatgtaacacgtctgaactccagtcac  ttactatgccgctggtggcttgcgttgaaatcttttcttttgctagattctagatgctacattgtgcagacttgaggtcagtg |
| L38 | 5’pho/atacggcgaccaccgaacgaaactttagaaaagaaaacgatctaagctctacgatgtaagacacgtctgaactccagtcac  ttactatgccgctggtggcttgctttgaaatcttttcttttgctagattcgagatgctacattctgtgcagacttgaggtcagtg |
| L39 | 5’pho/atacggcgaccaccgaaggcggttttacgctgtaaaacgatctacgctgttttaggcggtaaaacgatgtaagacacgtctgaactccagtcac  ttactatgccgctggtggcttccgccaaaatgcgacattttgctagatgcgacaaaatccgccattttgctacattctgtgcagacttgaggtcagtg |
| L40 | 5’pho/atacggcgaccaccgaaagctctacgatgtaagctctacgatgtaagacaacacgtctgaactccagtcac  ttactatgccgctggtggctttcgagatgctacattcgagatgctacattctgttgtgcagacttgaggtcagtg |
| L41 | 5’pho/atacggcgaccaccgaacgatgtaagctctacgatgtaagctctacgatgtaagacaacacgtctgaactccagtcac  ttactatgccgctggtggcttgctacattcgagatgctacattcgagatgctacattctgttgtgcagacttgaggtcagtg |
| L42 | 5’pho/atacggcgaccaccgaaggcggtacgatgtaagctctacgatgtaagacaacacgtctgaactccagtcac  ttactatgccgctggtggcttccgccatgctacattcgagatgctacattctgttgtgcagacttgaggtcagtg |
| L43 | 5’pho/atacggcgaccaccgaaggcggttttacgctgtaaaacgatgtaagctctacgatgtaagacaacacgtctgaactccagtcac  ttactatgccgctggtggcttccgccaaaatgcgacattttgctacattcgagatgctacattctgttgtgcagacttgaggtcagtg |
| L44 | 5’pho/atacggcgaccaccgaaggcggttttacgatgtacgctgtaaaacgatgtaagctctacgatgtaagacaacacgtctgaactccagtcac  ttactatgccgctggtggcttccgccaaaatgctacatgcgacattttgctacattcgagatgctacattctgttgtgcagacttgaggtcagtg |
| L45 | 5’pho/atacggcgaccaccgaaggcggtacgatgttttagaaaacaaaaagctctacgatgtaagacaacacgtctgaactccagtcac  ttactatgccgctggtggcttccgccatgctacaaaatcttttgtttttcgagatgctacattctgttgtgcagacttgaggtcagtg |
| L46 | 5’pho/atacggcgaccaccgaaggcggttttaggggttaaaacgatgtaagacaacacgtctgaactccagtcac  ttactatgccgctggtggcttccgccaaaatccccaattttgctacattctgttgtgcagacttgaggtcagtg |
| L47 | 5’pho/atacggcgaccaccgaacgatgttttagaaaacaaaaagctctacgatgtaagacaacacgtctgaactccagtcac  ttactatgccgctggtggcttgctacaaaatcttttgtttttcgagatgctacattctgttgtgcagacttgaggtcagtg |
| L48 | 5’pho/atacggcgaccaccgaagaaaagacgatgttttagaaaacaaaaagctctacgatgtaagacaacacgtctgaactccagtcac  ttactatgccgctggtggcttcttttctgctacaaaatcttttgtttttcgagatgctacattctgttgtgcagacttgaggtcagtg |
| L49 | 5’pho/atacggcgaccaccgaaagctcttttaggcgttaaaacgatgtaagacaacacgtctgaactccagtcac  ttactatgccgctggtggctttcgagaaaatccgcaattttgctacattctgttgtgcagacttgaggtcagtg |
| L50 | 5’pho/atacggcgaccaccgaacgatgtaagctcttttaggcgttaaaacgatgtaagacaacacgtctgaactccagtcac  ttactatgccgctggtggcttgctacattcgagaaaatccgcaattttgctacattctgttgtgcagacttgaggtcagtg |

**Sugar Dictionary (Carb Dictionary):**

**Table S4.** Monosaccharides and their codes (library A)

| **S.No** | **Sugar Monomer** | **Sugar Name** | **DNA Code (as in dictionary)** |
| --- | --- | --- | --- |
| **1.** | All | Allose | AAAA |
| **2.** | AllNAc | N-Acetyl-Allosamine | AAAC |
| **3.** | AllN | Allosamine | AAAG |
| **4.** | AllA | Alluronic Acid | AAAT |
| **5.** | Alt | Altrose | AACA |
| **6.** | AltNAc | N-Acetyl-Alltrosamine | AACC |
| **7.** | AltN | Alltrosamine | AACG |
| **8.** | AltA | Alturonic Acid | AACT |
| **9.** | Glc | Glucose | AAGA |
| **10.** | GlcNAc | N-Acetyl-Glucosamine | AAGC |
| **11.** | GlcN | Glucosamine | AAGG |
| **12.** | GlcA | Glucoronic Acid | AAGT |
| **13.** | Man | Mannose | AATA |
| **14.** | ManNAc | N-Acetyl-Mannosamine | AATC |
| **15.** | ManN | Mannosamine | AATG |
| **16.** | ManA | Mannuronic Acid | AATT |
| **17.** | G µL | Gulose | ACAA |
| **18.** | GulNAc | N-Acetyl-Gulosamine | ACAC |
| **19.** | GulN | Gulosamine | ACAG |
| **20.** | GulA | Guluronic Acid | ACAT |
| **21.** | Ido | Idose | ACCA |
| **22.** | IdoNAc | N-Acetyl-Idosamine | ACCC |
| **23.** | IdoN | Idosamine | ACCG |
| **24.** | IdoA | Idouronic Acid | ACCT |
| **25.** | Gal | Galactose | ACGA |
| **26.** | GalNAc | N-Acetyl-Galactosamine | ACGC |
| **27.** | GalN | Galactosamine | ACGG |
| **28.** | GalA | Galacturonic Acid | ACGT |
| **29.** | Tal | Talose | ACTA |
| **30.** | TalNAc | N-Acetyl-Talosamine | ACTC |
| **31.** | TalN | Talosamine | ACTG |
| **32.** | TalA | Taluronic Acid | ACTT |
| **33.** | Tag | Tagatose | ATCG |
| **34.** | TagA | Tagaturonic Acid | ATCT |
| **35.** | Hep | Heptose | ATCG |
| **36.** | DDManHep | D-Glycero-DMannoHeptose | ATGG |
| **37.** | Dha | 3-deoxy-D-lyxo-heptulosaric acid | ATGT |
| **38.** | LDManHep | L-glycero-D-mannoheptose | ATTA |
| **39.** | Fuc | Fucose | AGAA |
| **40.** | FucNAc | *N*-acetyl-L-fucosamine | AGAC |
| **41.** | FucN | Fucosamine | AGAG |
| **42.** | Qui | Quinovose | AGAT |
| **43.** | QuiNAc | *N*-acetyl-D-quinovosamine | ATAA |
| **44.** | QuiN | Quinovosamine | ATAC |
| **45.** | Rha | Rhamnose | ATAG |
| **46.** | RhaNAc | *N*-acetyl-L-rhamnosamine | ATAT |
| **47.** | RhaN | Rhamnosamine | ATCA |
| **48.** | Tyv | Tyvelose | ATTC |
| **49.** | Oli | Olivose | ATTG |
| **50.** | Par | Paratose | ATTT |
| **51.** | Dig | Digitoxose | CAAA |
| **52.** | 6dAlt | 6-deoxy-L-altrose | CAAC |
| **53.** | 6dTal | 6-deoxy-D-talose | CAAG |
| **54.** | Abe | Abequose | CAAT |
| **55.** | Api | Apiose | CACA |
| **56.** | Col | Colitose | CACC |
| **57.** | Neu5Ac | *N*-acetylneuraminic acid | AGGC |
| **58.** | Neu5GC | *N*-glycolylneuraminic acid | AGGG |
| **59.** | Kdn | 3-Deoxy-D-glycero-D-galacto-nonulosonic acid | AGGT |
| **60.** | Kdo | 3-Deoxy-D-manno-octulosonic acid | ATCC |
| **61.** | Mur | Muramic acid | CACG |
| **62.** | MurNAc | *N*-acetylmuramic acid | CACT |
| **63.** | MurNGc | *N*-glycolylmuramic acid | CCAA |
| **64.** | Sia | Sialic Acid | CCAG |
| **65.** | Neu | Neuraminic acid | CCAC |
| **66.** | Fru | Fructose | ATGA |
| **67.** | Sor | Sorbose | ATGC |
| **68.** | Psi | Psicose | ATAA |
| **69.** | Rib | Ribose | GGGG |
| **70.** | Ara | Arabinose | ATAT |
| **71.** | Xyl | Xylose | ATCA |
| **72.** | Lyx | Lyxose | ATCT |
| **73.** | Bac | Bacillosamine | CCAT |
| **74.** | R | For extending the short nucleotide sequences. | CCCCAGTCAGGCCTAACGTA |

**Linkages Dictionary (Link Dict)**

**Table S5.** Linkages/branching and their codes (library B)

| **S.No** | **Linkage** | **DNA Code(as in Dictionary)** |
| --- | --- | --- |
| **1.** | a1-1 | AAT |
| **2.** | a1-2 | AAG |
| **3.** | a1-3 | AAC |
| **4.** | a1-4 | ATA |
| **5.** | a1-5 | AGA |
| **6.** | a1-6 | ACA |
| **7.** | a1-7 | ATT |
| **8.** | a1-8 | ATC |
| **9.** | a2-1 | GTA |
| **10.** | a2-2 | GTC |
| **11.** | a2-3 | GGT |
| **12.** | a2-4 | GGC |
| **13.** | a2-5 | GGA |
| **14.** | a2-6 | GTT |
| **15.** | a2-7 | GAC |
| **16.** | a2-8 | GAT |
| **17.** | b1-1 | ATG |
| **18.** | b1-2 | TCA |
| **19.** | b1-3 | TCT |
| **20.** | b1-4 | TGT |
| **21.** | b1-5 | TCC |
| **22.** | b1-6 | TTA |
| **23.** | b1-7 | TTC |
| **24.** | b1-8 | GAA |
| **25.** | a? | GCC |
| **26.** | b? | CAA |
| **27.** | ?1 | CAC |
| **28.** | ?2 | CTC |
| **29.** | ?3 | CCA |
| **30.** | ?4 | GGG |
| **31.** | ?5 | CCT |
| **32.** | ?6 | AGG |
| **33.** | ?7 | CCG |
| **34.** | ?8 | CGC |
| **35.** | ?? | AGT |
| **36.** | ) | AAA |
| **37.** | ( | TTT |

**Modification Dictionary (Mod Dict):**

**Table S6.** Modifications and their codes (library C)

| # | Compound | Modification | **DNA Code (as in Dictionary)** |
| --- | --- | --- | --- |
| **1.** | Anhydrous | [2Y] | AGCA \| |
| **2.** |  | [4Y] | AGCC |
| **3.** |  | [7Y] | AGCG |
| **4.** |  | [8Y] | AGCT |
| **5.** |  | [?Y] | CCCA |
| **6.** | Hydroxyl | [2OH] | AGTA |
| **7.** |  | [4OH] | AGTC |
| **8.** |  | [7OH] | AGTG |
| **9.** |  | [8OH] | AGTT |
| **10.** |  | [?OH] | CAGA |
| **11.** | Pyruvate | [2V] | CAGC |
| **12.** |  | [4V] | CAGG |
| **13.** |  | [7V] | CAGT |
| **14.** |  | [8V] | CATA |
| **15.** |  | [?V] | CATC |
| **16.** | Sulphate  0 | [2S] | CATG |
| **17.** |  | [4S] | CATT |
| **18.** |  | [7S] | CGAA |
| **19.** |  | [8S] | CGAC |
| **20.** |  | [?S] | CGAG |
| **21.** | Phosphate | [2P] | CGAT |
| **22.** |  | [4P] | CGGA |
| **23.** |  | [7P] | CGGC |
| **24.** |  | [8P] | CGGG |
| **25.** |  | [?P] | CGGT |
| **26.** | n-glycolyl | [2J] | CGTA |
| **27.** |  | [4J] | CGTC |
| **28.** |  | [7J] | CGTG |
| **29.** |  | [8J] | CGTT |
| **30.** |  | [?J] | CTAA |
| **31.** | n-acetyl | [2NAc] | CTAC |
| **32.** |  | [4NAc] | CTAG |
| **33.** |  | [7NAc] | CTAT |
| **34.** |  | [8NAc] | CTGA |
| **35.** |  | [?NAc] | CTGC |
| **36.** | o-acetyl | [2Ac] | CTGG |
| **37.** |  | [4Ac] | CTGT |
| **38.** |  | [7Ac] | CTTA |
| **39.** |  | [8Ac] | CTTC |
| **40.** |  | [?Ac] | CTTG |
| **41.** | Carboxylate | [2COOH] | CTTT |
| **42.** |  | [4COOH] | GAGA |
| **43.** |  | [7COOH] | GAGC |
| **44.** |  | [8COOH] | GAGG |
| **45.** |  | [?COOH] | GAGT |
| **46.** | Inositol | [2IN] | GCAA |
| **47.** |  | [4IN] | GCAC |
| **48.** |  | [7IN] | GCAG |
| **49.** |  | [8IN] | GCAT |
| **50.** |  | [?IN] | GCGA |
| **51.** | pentyl | [2EE] | GCGC |
| **52.** |  | [4EE] | GCGG |
| **53.** |  | [7EE] | GCGT |
| **54.** |  | [8EE] | GCTA |
| **55.** |  | [?EE] | GCTC |
| **56.** | octyl | [2EH] | GCTG |
| **57.** |  | [4EH] | GCTT |
| **58.** |  | [7EH] | GTGA |
| **59.** |  | [8EH] | GTGC |
| **60.** |  | [?EH] | GTGG |
| **61.** | deactylated-n-actyl | [2Q] | GTGT |
| **62.** |  | [4Q] | TAAA |
| **63.** |  | [7Q] | TAAC |
| **64.** |  | [8Q] | TAAG |
| **65.** |  | [?Q] | TAAT |
| **66.** | N-Sulfate | [2QS] | TACA |
| **67.** |  | [4QS] | TACC |
| **68.** |  | [7QS] | TACG |
| **69.** |  | [8QS] | TACT |
| **70.** |  | [?QS] | TAGA |
| **71.** | Pyruvate Acetal | [2PYR] | TAGC |
| **72.** |  | [4PYR] | TAGG |
| **73.** |  | [7PYR] | TAGT |
| **74.** |  | [8PYR] | TATA |
| **75.** |  | [?PYR] | TATC |
| **76.** | N-methylcarbomoyl | [2ECO] | TATG |
| **77.** |  | [4ECO] | TATT |
| **78.** |  | [7ECO] | TCGA |
| **79.** |  | [8ECO] | TCGC |
| **80.** |  | [?ECO] | TCGG |
| **81.** | Phosphocholine | [2PC] | TCGT |
| **82.** |  | [4PC] | TGAA |
| **83.** |  | [7PC] | TGAC |
| **84.** |  | [8PC] | TGAG |
| **85.** |  | [?PC] | TGAT |
| **86.** | Phosphoethnlamine | [2PE] | TGCA |
| **87.** |  | [4PE] | TGCC |
| **88.** |  | [7PE] | TGCG |
| **89.** |  | [8PE] | TGCT |
| **90.** |  | [?PE] | TGGA |
| **91.** | Methyl | [2ME] | TGGC |
| **92.** |  | [3ME] | TGGG |
| **93.** |  | [4ME] | TGGT |
| **94.** |  | [7ME] | TTGA |
| **95.** |  | [8ME] | TTGC |
| **96.** |  | [?ME] | TTGG |
| **97.** | Sulphate | [3S] | TTGT |

**Core Dictionary (Core Dict):**

**Table S7.** Core structure and their codes (Library D)

| S.No | Sugar blocks for Secondary naming | CFG 2D structure | Long Nucleotide | Length | Assigned short DNA code |
| --- | --- | --- | --- | --- | --- |
| **1.** | *Man* b1-4 *GlcNAc* b1-4 (*Fuc* a1-6) *GlcNAc* | 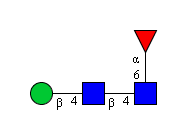 | AATATGTAAGCTGTTTTAGAAACAAAAAAGC | 31 | GTACA |
| **2.** | *Man* a1-6*Man* b1-4 *GlcNAc* b1-4 *GlcNAc* | 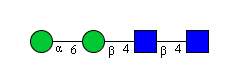 | AATAACAAATATGTAAGCTGTAAGC | 25 | GTACT |
| **3.** | *Gal* b1-4 *GlcNAc* b1-4 | 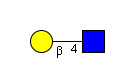 | ACGATGTAAGCTGT | 14 | GTCCT |
| ***4.*** | *Gal* a1-3 *Gal* b1-4 *GlcNAc* b1-2 | 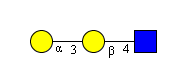 | ACGAAACACGATGTAAGCTCA | 21 | GTACC |
| ***5.*** | *Gal* a1-3 *Gal* b1-4 *GlcNAc* b1-3 | 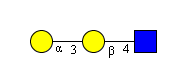 | ACGAAACACGATGTAAGCTCT | 21 | GTTAT |
| ***6.*** | *Gal* b1-3 *GlcNAc* b1-4 | 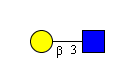 | ACGATCTAAGCTGT | 14 | GTTCA |
| **7.** | *Man* a1-3 *Man* a1-6 | 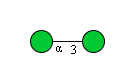 | AATAAACAATAACA | 14 | GTTTG |
| ***8.*** | *GalNAc* b1-4 *GlcNAc* b1-2 | 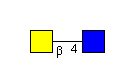 | ACGCTGTAAGCTCA | 14 | GTTTC |
| ***9.*** | *Fuc* a1-3 *GlcNAc* b1-2 | 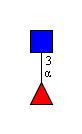 | AGAAAACAAGCTCA | 14 | GTTGT |
| ***10.*** | *Fuc* a1-2 *Gal* b1-4 (*Fuc* a1-3) *GlcNAc* b1-2 | 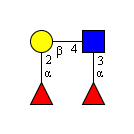 | AGAAAAGACGATGTTTTAGAAAACAAAAAGCTCA | 34 | GGATT |
| **11.** | *Fuc* a1-2 *Gal* b1-4 *GlcNAc* b1-2 | 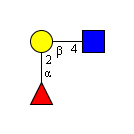 | AGAAAAGACGATGTAAGCTCA | 21 | GTTGC |
| ***12.*** | *Gal* b1-4 (*Fuc* a1-3) *GlcNAc* b1-4 | 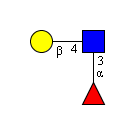 | ACGATGTTTTAGAAAACAAAAAGCTGT | 27 | GTGAG |
| ***13.*** | *GalNAc* b1-4 *GlcNAc* b1-6 | 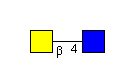 | ACGCTGTAAGCTTA | 14 | GTGTC |
| **14.** | *Gal* b1-4 *GlcNAc* b1-6 | 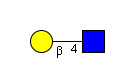 | ACGATGTAAGCTTA | 14 | GTGCG |
| ***15.*** | *NeuAc* a2-3 *Gal* b1-4 *GlcNAc* b1-4 | 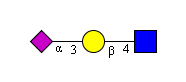 | AGGCGGTACGATGTAAGCTGT | 21 | GTGGA |
| ***16.*** | *NeuAc* a2-3 *Gal* b1-4 *GlcNAc* b1-6 | 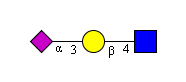 | AGGCGGTACGATGTAAGCTTA | 21 | GTGGT |

**Coding Examples:**

**Table S8.** Examples of DNA encoding of glycans using the program

| Structure | Long Nucleotide | Length | Short Nucleotide | Length |
| --- | --- | --- | --- | --- |
| 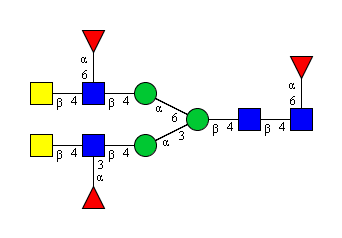 | acgctgttttagaaaagaaaaagctgtaataaactttacgctgttttagaaaagaaaaagctgtaataacaaaaaatatgtaagctgttttagaaacaaaaaagc | 105 | acgctgttttagaaaagaaaaagctgtaataaactttacgctgttttagaaaagaaaaagctgtaataacaaaagtaca | 79 |
| 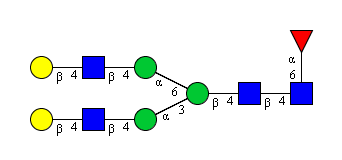 | acgatgtaagctgtaataaactttacgatgtaagctgtaataacaaaaaatatgtaagctgttttagaaacaaaaaagc | 79 | gtacgaataaactttacgatgtaagctgtaataacaaaagtaca | 44 |
| 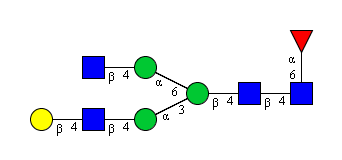 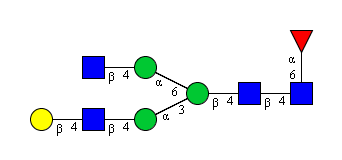 | acgatgtaagctgtaataaactttaagctgtaataacaaaaaatatgtaagctgttttagaaacaaaaaagc | 72 | gtacgaataaactttaagctgtaataacaaaagtaca | 37 |
| 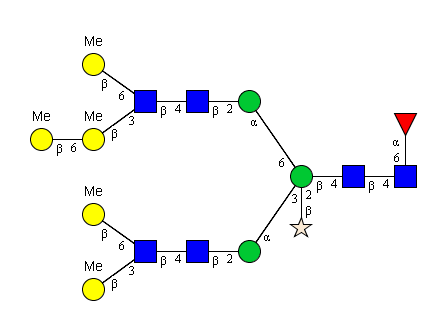 | acgatggttcttttacgatgggttaaaaaagctgtaagctcaaataaactttacgatgggttaacgatgggtcttttacgatgggttaaaaaagctgtaagctcaaataacaaaatttatcatcaaaaaatatgtaagctgttttagaaacaaaaaagc | 159 | acgatggttcttttacgatgggttaaaaaagctgtaagctcaaataaactttacgatgggttaacgatgggtcttttacgatgggttaaaaaagctgtaagctcaaataacaaaatttatcatcaaaagtaca | 133 |
| 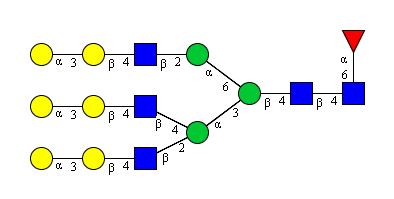 | acgaaacacgatgtaagctcatttacgaaacacgatgtaagctgtaaaaataaactttacgaaacacgatgtaagctcaaataacaaaaaatatgtaagctgttttagaaacaaaaaagc | 120 | acgaaacacgatgtaagctcatttacgaaacgtacgaaaaataaactttgtaccaataacaaaagtaca | 69 |
| 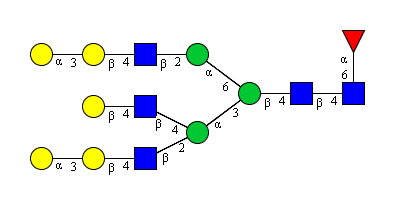 | acgaaacacgatgtaagctcatttacgatgtaagctgtaaaaataaactttacgaaacacgatgtaagctcaaataacaaaaaatatgtaagctgttttagaaacaaaaaagc | 113 | acgaaacacgatgtaagctcatttgtacgaaaaataaactttgtaccaataacaaaagtaca | 62 |
| 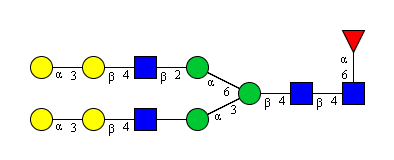 | acgaaacacgatgtaagctcaaataaactttacgaaacacgatgtaagctcaaataacaaaaaatatgtaagctgttttagaaacaaaaaagc | 93 | gtaccaataaactttacgaaacacgatgtaagctcaaataacaaaagtaca | 51 |
| 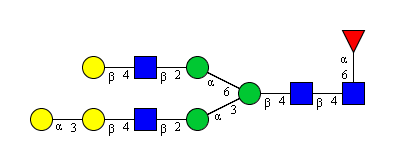 | acgaaacacgatgtaagctcaaataaactttacgatgtaagctcaaataacaaaaaatatgtaagctgttttagaaacaaaaaagc | 86 | gtaccaataaactttacgatgtaagctcaaataacaaaagtaca | 44 |
| 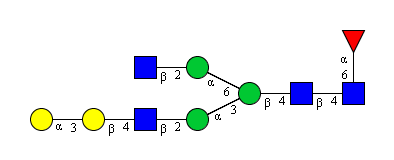 | acgaaacacgatgtaagctcaaataaactttaagctcaaataacaaaaaatatgtaagctgttttagaaacaaaaaagc | 79 | gtaccaataaactttaagctcaaataacaaaagtaca | 37 |
| 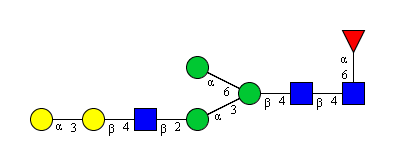 | acgaaacacgatgtaagctcaaataaactttaataacaaaaaatatgtaagctgttttagaaacaaaaaagc | 72 | gtaccaataaactttaataacaaaagtaca | 30 |
| 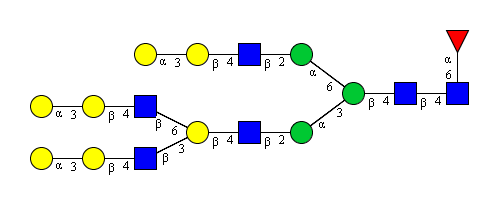 | acgaaacacgatgtaagctcttttacgaaacacgatgtaagcttaaaaacgatgtaagctcaaataaactttacgaaacacgatgtaagctcaaataacaaaaaatatgtaagctgttttagaaacaaaaaagc | 134 | gttattttacgaaacacgatgtaagcttaaaaacgatgtaagctcaaataaactttgtaccaataacaaaagtaca | 76 |
| 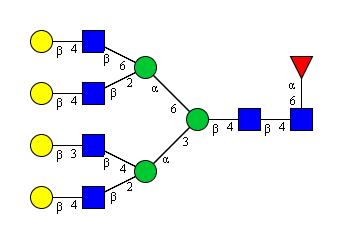 | acgatgtaagctcatttacgatctaagctgtaaaaataaactttacgatgtaagctcatttacgatgtaagcttaaaaaataacaaaaaatatgtaagctgttttagaaacaaaaaagc | 119 | acgatgtaagctcatttgttcaaaaaataaactttacgatgtaagctcatttacgatgtaagcttaaaaaataacaaaagtaca | 84 |
| 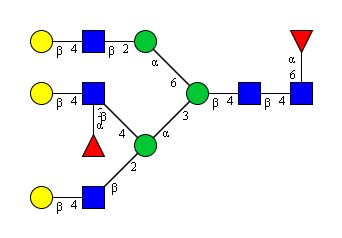 | acgatgtaagctcatttacgatgttttagaaaacaaaaagctgtaaaaataaactttacgatgtaagctcaaataacaaaaaatatgtaagctgttttagaaacaaaaaagc | 112 | acgatgtaagctcatttgtgagaaaaataaactttacgatgtaagctcaaataacaaaagtaca | 64 |
| 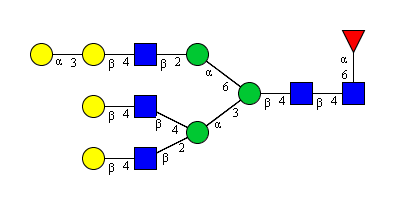 | acgatgtaagctcatttacgatgtaagctgtaaaaataaactttacgaaacacgatgtaagctcaaataacaaaaaatatgtaagctgttttagaaacaaaaaagc | 106 | acgatgtaagctcatttgtacgaaaaataaactttgtaccaataacaaaagtaca | 55 |
| 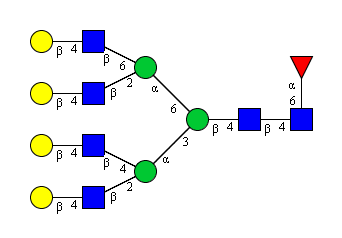 | acgatgtaagctcatttacgatgtaagctgtaaaaataaactttacgatgtaagctcatttacgatgtaagcttaaaaaataacaaaaaatatgtaagctgttttagaaacaaaaaagc | 119 | acgatgtaagctcatttgtacgaaaaataaactttacgatgtaagctcatttacgatgtaagcttaaaaaataacaaaagtaca | 84 |
| 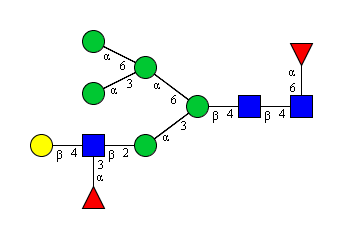 | acgatgttttagaaaacaaaaagctcaaataaactttaataaactttaataacaaaaaataacaaaaaatatgtaagctgttttagaaacaaaaaagc | 98 | acgatgttttagaaaacaaaaagctcaaataaactttaataaactttaataacaaaaaataacaaaagtaca | 72 |
| 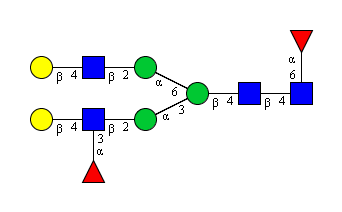 | acgatgttttagaaaacaaaaagctcaaataaactttacgatgtaagctcaaataacaaaaaatatgtaagctgttttagaaacaaaaaagc | 92 | acgatgttttagaaaacaaaaagctcaaataaactttacgatgtaagctcaaataacaaaagtaca | 66 |
| 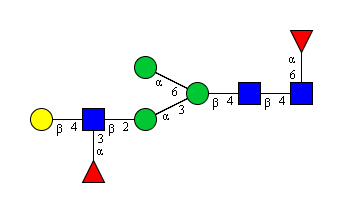 | acgatgttttagaaaacaaaaagctcaaataaactttaataacaaaaaatatgtaagctgttttagaaacaaaaaagc | 78 | acgatgttttagaaaacaaaaagctcaaataaactttaataacaaaagtaca | 52 |
| 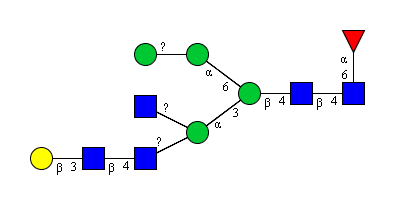 | acgatctaagctgtaagcagttttaagcagtaaaaataaactttaataagtaataaacaaaaatatgtaagctgttttagaaacaaaaaagc | 92 | gttcaaagcagttttaagcagtaaaaataaactttaataagtaataaacaaagtaca | 57 |
| 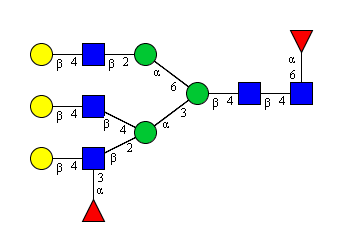 | acgatgttttagaaaacaaaaagctcatttacgatgtaagctgtaaaaataaactttacgatgtaagctcaaataacaaaaaatatgtaagctgttttagaaacaaaaaagc | 112 | acgatgttttagaaaacaaaaagctcatttgtacgaaaaataaactttacgatgtaagctcaaataacaaaagtaca | 77 |
| 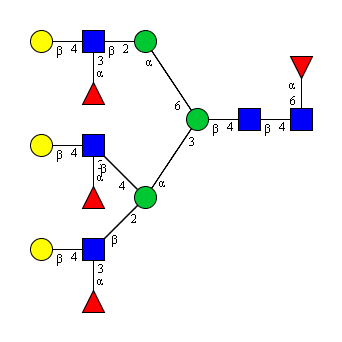 | acgatgttttagaaaacaaaaagctcatttacgatgttttagaaaacaaaaagctgtaaaaataaactttacgatgttttagaaaacaaaaagctcaaataacaaaaaatatgtaagctgttttagaaacaaaaaagc | 138 | acgatgttttagaaaacaaaaagctcatttgtgagaaaaataaactttacgatgttttagaaaacaaaaagctcaaataacaaaagtaca | 90 |
| 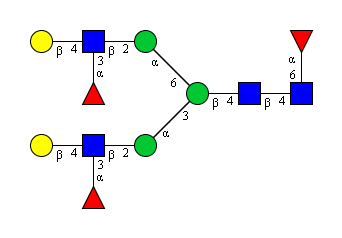 | acgatgttttagaaaacaaaaagctcaaataaactttacgatgttttagaaaacaaaaagctcaaataacaaaaaatatgtaagctgttttagaaacaaaaaagc | 105 | acgatgttttagaaaacaaaaagctcaaataaactttacgatgttttagaaaacaaaaagctcaaataacaaaagtaca | 79 |
| 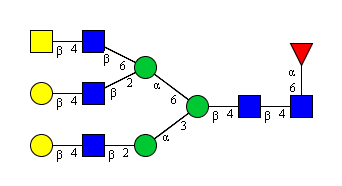 | acgatgtaagctcaaataaactttacgatgtaagctcatttacgctgtaagcttaaaaaataacaaaaaatatgtaagctgttttagaaacaaaaaagc | 99 | acgatgtaagctcaaataaactttacgatgtaagctcatttgtgtcaaaaataacaaaagtaca | 64 |
| 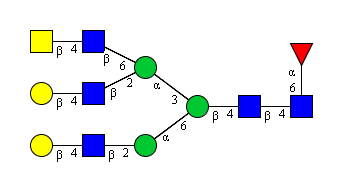 | acgatgtaagctcaaataacatttacgatgtaagctcatttacgctgtaagcttaaaaaataaacaaaaatatgtaagctgttttagaaacaaaaaagc | 99 | acgatgtaagctcaaataacatttacgatgtaagctcatttgtgtcaaaaataaacaaagtaca | 64 |
| 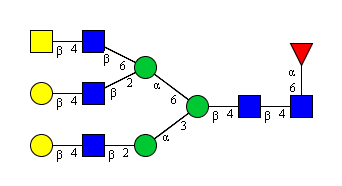 | acgatgtaagctcaaataaactttacgctgtaagctcatttacgatgtaagcttaaaaaataacaaaaaatatgtaagctgttttagaaacaaaaaagc | 99 | acgatgtaagctcaaataaactttgtttctttacgatgtaagcttaaaaaataacaaaagtaca | 64 |
| 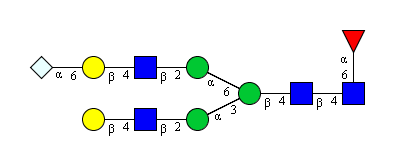 | acgatgtaagctcaaataaactttaggggttacgatgtaagctcaaataacaaaaaatatgtaagctgttttagaaacaaaaaagc | 86 | acgatgtaagctcaaataaactttaggggttacgatgtaagctcaaataacaaaagtaca | 60 |
| 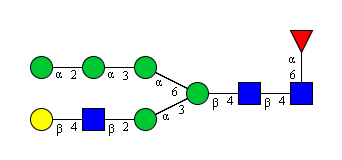 | acgatgtaagctcaaataaactttaataaagaataaacaataacaaaaaatatgtaagctgttttagaaacaaaaaagc | 79 | acgatgtaagctcaaataaactttaataaagaataaacaataacaaaagtaca | 53 |
| 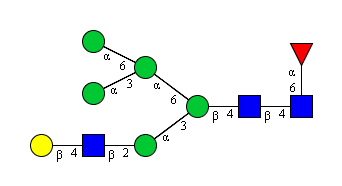 | acgatgtaagctcaaataaactttaataaactttaataacaaaaaataacaaaaaatatgtaagctgttttagaaacaaaaaagc | 85 | acgatgtaagctcaaataaactttaataaactttaataacaaaaaataacaaaagtaca | 59 |
| 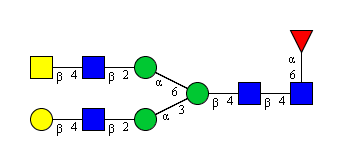 | acgatgtaagctcaaataaactttacgctgtaagctcaaataacaaaaaatatgtaagctgttttagaaacaaaaaagc | 79 | acgatgtaagctcaaataaactttgtttcaataacaaaagtaca | 44 |
| 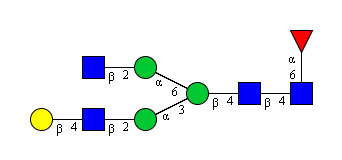 | acgatgtaagctcaaataaactttaagctcaaataacaaaaaatatgtaagctgttttagaaacaaaaaagc | 72 | acgatgtaagctcaaataaactttaagctcaaataacaaaagtaca | 46 |
| 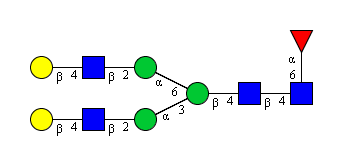 | acgatgtaagctcaaataaactttacgatgtaagctcaaataacaaaaaatatgtaagctgttttagaaacaaaaaagc | 79 | acgatgtaagctcaaataaactttacgatgtaagctcaaataacaaaagtaca | 53 |
| 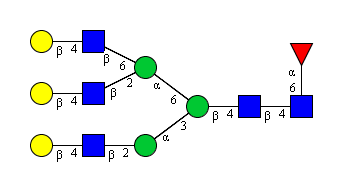 | acgatgtaagctcaaataaactttacgatgtaagctcatttacgatgtaagcttaaaaaataacaaaaaatatgtaagctgttttagaaacaaaaaagc | 99 | acgatgtaagctcaaataaactttacgatgtaagctcatttgtgcgaaaaataacaaaagtaca | 64 |
| 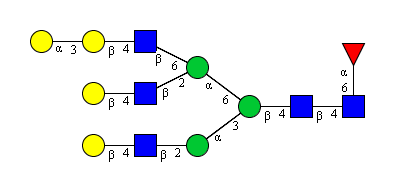 | acgatgtaagctcaaataaactttacgatgtaagctcatttacgaaacacgatgtaagcttaaaaaataacaaaaaatatgtaagctgttttagaaacaaaaaagc | 106 | acgatgtaagctcaaataaactttacgatgtaagctcatttacgaaacgtgcgaaaaataacaaaagtaca | 71 |
| 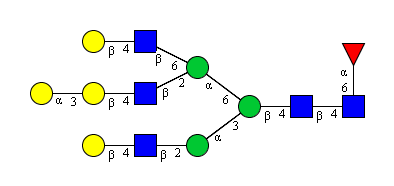 | acgatgtaagctcaaataaactttacgaaacacgatgtaagctcatttacgatgtaagcttaaaaaataacaaaaaatatgtaagctgttttagaaacaaaaaagc | 106 | acgatgtaagctcaaataaactttacgaaacacgatgtaagctcatttgtgcgaaaaataacaaaagtaca | 71 |
| 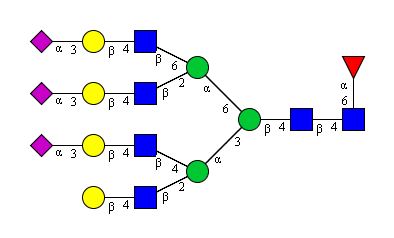 | acgatgtaagctcatttaggcggtacgatgtaagctgtaaaaataaactttaggcggtacgatgtaagctcatttaggcggtacgatgtaagcttaaaaaataacaaaaaatatgtaagctgttttagaaacaaaaaagc | 140 | acgatgtaagctcatttaggcggtacgatgtaagctgtaaaaataaactttaggcggtacgatgtaagctcatttgtggtaaaaataacaaaagtaca | 98 |
| 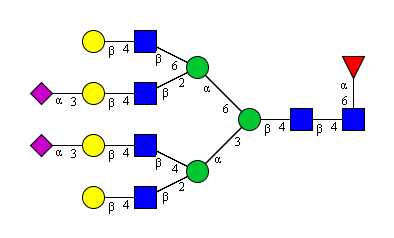 | acgatgtaagctcatttaggcggtacgatgtaagctgtaaaaataaactttaggcggtacgatgtaagctcatttacgatgtaagcttaaaaaataacaaaaaatatgtaagctgttttagaaacaaaaaagc | 133 | acgatgtaagctcatttgtggaaaaaataaactttaggcggtacgatgtaagctcatttgtgcgaaaaataacaaaagtaca | 82 |
| 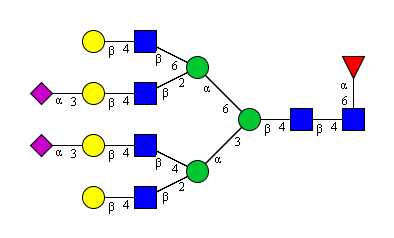 | acgatgtaagctcatttaggcggtacgatgtaagctgtaaaaataaactttaggcggtacgatgtaagctcatttacgatgtaagcttaaaaaataacaaaaaatatgtaagctgttttagaaacaaaaaagc | 133 | acgatgtaagctcatttgtggaaaaaataaactttaggcggtacgatgtaagctcatttgtgcgaaaaataacaaaagtaca | 82 |
| 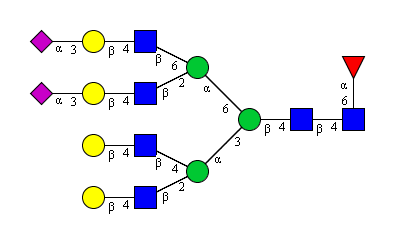 | acgatgtaagctcatttacgatgtaagctgtaaaaataaactttaggcggtacgatgtaagctcatttaggcggtacgatgtaagcttaaaaaataacaaaaaatatgtaagctgttttagaaacaaaaaagc | 133 | acgatgtaagctcatttacgatgtaagctgtaaaaataaactttaggcggtacgatgtaagctcatttgtggtaaaaataacaaaagtaca | 91 |
| 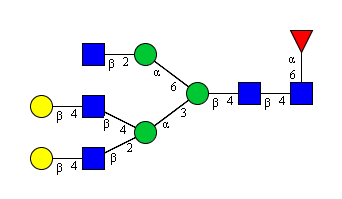 | acgatgtaagctcatttacgatgtaagctgtaaaaataaactttaagctcaaataacaaaaaatatgtaagctgttttagaaacaaaaaagc | 92 | acgatgtaagctcatttgtacgaaaaataaactttaagctcaaataacaaaagtaca | 57 |
| 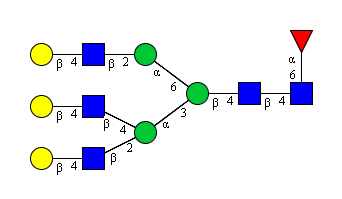 | acgatgtaagctcatttacgatgtaagctgtaaaaataaactttacgatgtaagctcaaataacaaaaaatatgtaagctgttttagaaacaaaaaagc | 99 | acgatgtaagctcatttgtacgaaaaataaactttacgatgtaagctcaaataacaaaagtaca | 64 |
| 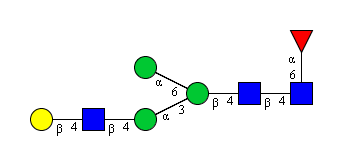 | acgatgtaagctgtaataaactttaataacaaaaaatatgtaagctgttttagaaacaaaaaagc | 65 | gtacgaataaactttaataacaaaagtaca | 30 |
| 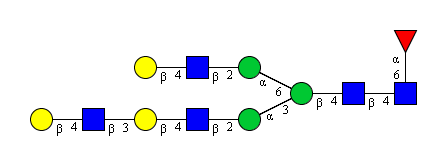 | acgatgtaagctctacgatgtaagctcaaataaactttacgatgtaagctcaaataacaaaaaatatgtaagctgttttagaaacaaaaaagc | 93 | acgatgtaagctctacgatgtaagctcaaataaactttacgatgtaagctcaaataacaaaagtaca | 67 |
| 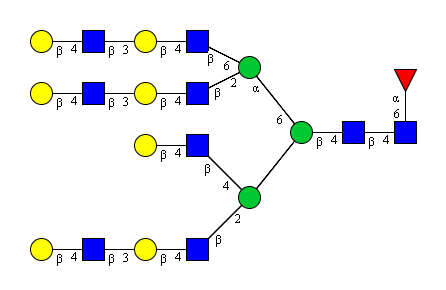 | acgatgtaagctctacgatgtaagctcatttacgatgtaagctgtaaaaataaactttacgatgtaagctctacgatgtaagctcatttacgatgtaagctctacgatgtaagcttaaaaaataacaaaaaatatgtaagctgttttagaaacaaaaaagc | 161 | acgatgtaagctctacgatgtaagctcatttacgatgtaagctgtaaaaataaactttacgatgtaagctctacgatgtaagctcatttacgatgtaagctctgtgcgaaaaataacaaaagtaca | 126 |
| 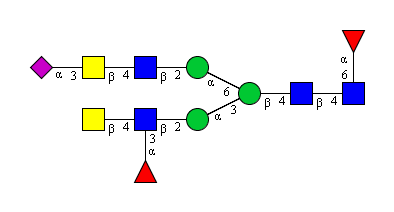 | acgctgttttagaaaacaaaaagctcaaataaactttaggcggtacgctgtaagctcaaataacaaaaaatatgtaagctgttttagaaacaaaaaagc | 99 | acgctgttttagaaaacaaaaagctcaaataaactttaggcggtgtttcaataacaaaagtaca | 64 |
| 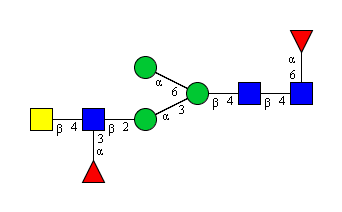 | acgctgttttagaaaacaaaaagctcaaataaactttaataacaaaaaatatgtaagctgttttagaaacaaaaaagc | 78 | acgctgttttagaaaacaaaaagctcaaataaactttaataacaaaagtaca | 52 |
| 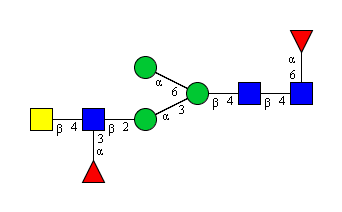 | acgctgttttagaaaacaaaaagctcaaataaactttaataacaaaaaatatgtaagctgttttagaaacaaaaaagc | 78 | acgctgttttagaaaacaaaaagctcaaataaactttaataacaaaagtaca | 52 |
| 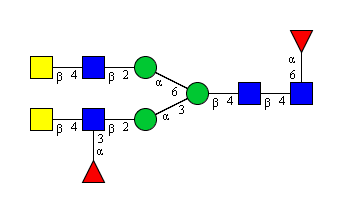 | acgctgttttagaaaacaaaaagctcaaataaactttacgctgtaagctcaaataacaaaaaatatgtaagctgttttagaaacaaaaaagc | 92 | acgctgttttagaaaacaaaaagctcaaataaactttgtttcaataacaaaagtaca | 57 |
| 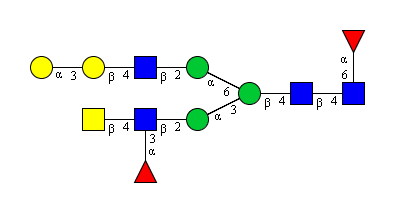 | acgctgttttagaaaacaaaaagctcaaataaactttacgaaacacgatgtaagctcaaataacaaaaaatatgtaagctgttttagaaacaaaaaagc | 99 | acgctgttttagaaaacaaaaagctcaaataaactttgtaccaataacaaaagtaca | 57 |
| 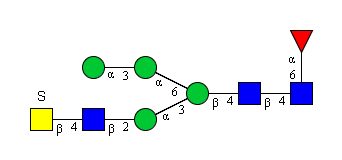 | acgccatttgtaagctcaaataaactttaataaacaataacaaaaaatatgtaagctgttttagaaacaaaaaagc | 76 | acgccatttgtaagctcaaataaactttaataaacaataacaaaagtaca | 50 |
| 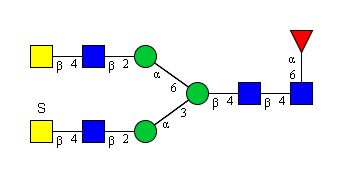 | acgccatttgtaagctcaaataaactttacgctgtaagctcaaataacaaaaaatatgtaagctgttttagaaacaaaaaagc | 83 | acgccatttgtaagctcaaataaactttgtttcaataacaaaagtaca | 48 |
| 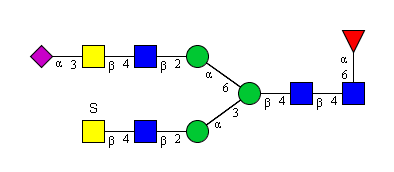 | acgccatttgtaagctcaaataaactttaggcggtacgatgtaagctcaaataacaaaaaatatgtaagctgttttagaaacaaaaaagc | 90 | acgccatttgtaagctcaaataaactttaggcggtacgatgtaagctcaaataacaaaagtaca | 64 |
| 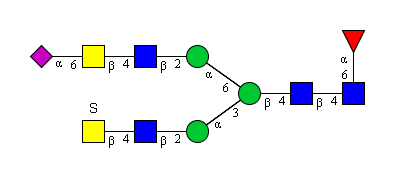 | acgccatttgtaagctcaaataaactttaggcgttacgatgtaagctcaaataacaaaaaatatgtaagctgttttagaaacaaaaaagc | 90 | acgccatttgtaagctcaaataaactttaggcgttacgatgtaagctcaaataacaaaagtaca | 64 |
| 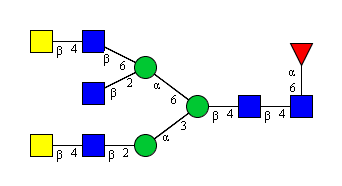 | acgctgtaagctcaaataaactttaagctcatttacgctgtaagcttaaaaaataacaaaaaatatgtaagctgttttagaaacaaaaaagc | 92 | acgctgtaagctcaaataaactttaagctcatttgtgtcaaaaataacaaaagtaca | 57 |
| 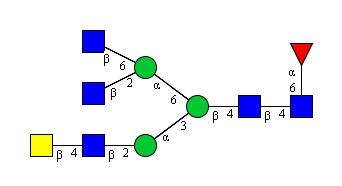 | acgctgtaagctcaaataaactttaagctcatttaagcttaaaaaataacaaaaaatatgtaagctgttttagaaacaaaaaagc | 85 | gtttcaataaactttaagctcatttaagcttaaaaaataacaaaagtaca | 50 |
| 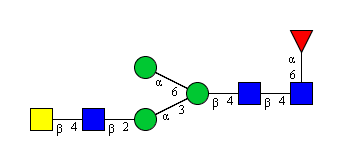 | acgctgtaagctcaaataaactttaataacaaaaaatatgtaagctgttttagaaacaaaaaagc | 65 | gtttcaataaactttaataacaaaagtaca | 30 |
| 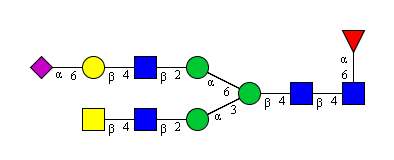 | acgctgtaagctcaaataaactttaggcgttacgatgtaagctcaaataacaaaaaatatgtaagctgttttagaaacaaaaaagc | 86 | gtttcaataaactttaggcgttacgatgtaagctcaaataacaaaagtaca | 51 |
| 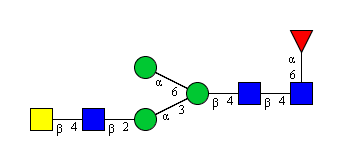 | acgctgtaagctcaaataaactttaataacaaaaaatatgtaagctgttttagaaacaaaaaagc | 65 | gtttcaataaactttaataacaaaagtaca | 30 |
| 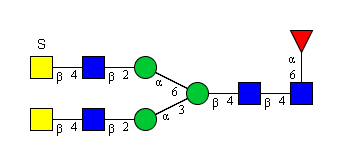 | acgctgtaagctcaaataaactttacgccatttgtaagctcaaataacaaaaaatatgtaagctgttttagaaacaaaaaagc | 83 | gtttcaataaactttacgccatttgtaagctcaaataacaaaagtaca | 48 |
| 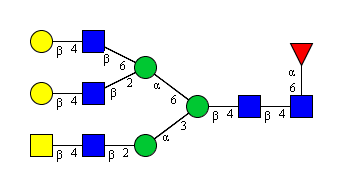 | acgctgtaagctcaaataaactttacgatgtaagctcatttacgatgtaagcttaaaaaataacaaaaaatatgtaagctgttttagaaacaaaaaagc | 99 | gtttcaataaactttacgatgtaagctcatttgtgcgaaaaataacaaaagtaca | 55 |
| 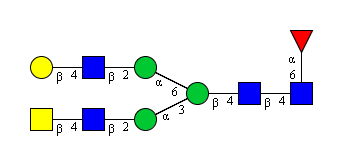 | acgctgtaagctcaaataaactttacgatgtaagctcaaataacaaaaaatatgtaagctgttttagaaacaaaaaagc | 79 | gtttcaataaactttacgatgtaagctcaaataacaaaagtaca | 44 |
| 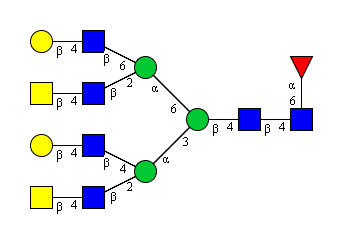 | acgctgtaagctcatttacgatgtaagctgtaaaaataaactttacgctgtaagctcatttacgatgtaagcttaaaaaataacaaaaaatatgtaagctgttttagaaacaaaaaagc | 119 | gtttctttacgatgtaagctgtaaaaataaactttacgctgtaagctcatttgtgcgaaaaataacaaaagtaca | 75 |
| 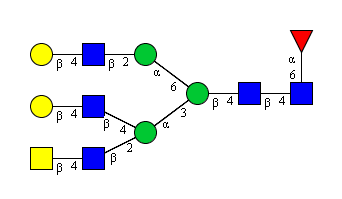 | acgctgtaagctcatttacgatgtaagctgtaaaaataaactttacgatgtaagctcaaataacaaaaaatatgtaagctgttttagaaacaaaaaagc | 99 | gtttctttgtcctaaaaataaactttacgatgtaagctcaaataacaaaagtaca | 55 |
| 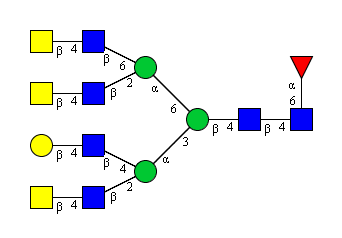 | acgctgtaagctcatttacgatgtaagctgtaaaaataaactttacgctgtaagctcatttacgctgtaagcttaaaaaataacaaaaaatatgtaagctgttttagaaacaaaaaagc | 119 | acgctgtaagctcatttacgatgtaagctgtaaaaataaactttacgctgtaagctcatttgtgtcaaaaataacaaaagtaca | 84 |
|  | acgctgtaagctcatttaagctgtaaaaataaactttaagctcatttaagcttaaaaaataacaaaaaatatgtaagctgttttagaaacaaaaaagc | 98 | gttgaaataaactttaagctcatttaagcttaaaaaataacaaaagtaca | 50 |
|  | acgctgtaagctcatttaagctgtaaaaataaactttacgctgtaagctcatttaagcttaaaaaataacaaaaaatatgtaagctgttttagaaacaaaaaagc | 105 | gtttctttaagctgtaaaaataaactttacgctgtaagctcatttaagcttaaaaaataacaaaagtaca | 70 |
|  | acgctgtaagctcatttaagctgtaaaaataaactttaagctcaaataacaaaaaatatgtaagctgttttagaaacaaaaaagc | 85 | gtttctttaagctgtaaaaataaactttaagctcaaataacaaaagtaca | 50 |
|  | acgctgtaagctcaaataaactttacgatgtaagctcatttacgatgtaagcttaaaaaataacaaaaaatatgtaagctgttttagaaacaaaaaagc | 99 | gtttcaataaactttacgatgtaagctcatttgtgcgaaaaataacaaaagtaca | 55 |
|  | acgctgtaagctcaaataaactttacgatgtaagctcaaataacaaaaaatatgtaagctgttttagaaacaaaaaagc | 79 | gtttcaataaactttacgatgtaagctcaaataacaaaagtaca | 44 |
|  | acgctgtaagctcaaataaactttacgctgtaagctcaaataacaaaaaatatgtaagctgttttagaaacaaaaaagc | 79 | gtttcaataaactttacgctgtaagctcaaataacaaaagtaca | 44 |
|  | acgctgtaagctcaaataaactttacgctgtaagctcaaataacaaaaaatatgtaagctgttttagaaacaaaaaagc | 79 | gtttcaataaactttacgctgtaagctcaaataacaaaagtaca | 44 |
|  | agaaaacaagctcaaataaactttaataacaaaaaatatgtaagctgttttagaaacaaaaaagc | 65 | gttgtaataaactttaataacaaaagtaca | 30 |
|  | agaaaagacgatgttttagaaaacaaaaagctcaaataaactttagaaaagacgatgtaagctcaaataacaaaaaatatgtaagctgttttagaaacaaaaaagc | 106 | gttggaataaactttgttgcaataacaaaagtaca | 35 |
|  | agaaaagacgatgttttagaaaacaaaaagctcaaataaactttagaaaagacgatgttttagaaaacaaaaagctcaaataacaaaaaatatgtaagctgttttagaaacaaaaaagc | 119 | gttggaataaactttagaaaagacgatgttttagaaaacaaaaagctcaaataacaaaagtaca | 64 |
|  | agaaaagacgatgttttagaaaacaaaaagctcaaataaactttaataaactttaataacaaaaaataacaaaaaatatgtaagctgttttagaaacaaaaaagc | 105 | gttggaataaactttaataaactttaataacaaaaaataacaaaagtaca | 50 |
|  | agaaaagacgatgttttagaaaacaaaaagctcaaataaactttaatagccaataacaaaaaatatgtaagctgttttagaaacaaaaaagc | 92 | gttggaataaactttaatagccaataacaaaagtaca | 37 |
|  | agaaaagacgatgttttagaaaacaaaaagctcaaataaactttacgatgttttagaaaacaaaaagctcaaataacaaaaaatatgtaagctgttttagaaacaaaaaagc | 112 | gttggaataaactttacgatgttttagaaaacaaaaagctcaaataacaaaagtaca | 57 |
|  | agaaaagacgatgttttagaaaacaaaaagctcatttagaaaagacgatgttttagaaaacaaaaagctgtaaaaataaactttagaaaagacgatgttttagaaaacaaaaagctcaaataacaaaaaatatgtaagctgttttagaaacaaaaaagc | 159 | gttggtttagaaaaggtgagaaaaataaactttagaaaagacgatgttttagaaaacaaaaagctcaaataacaaaagtaca | 82 |
|  | agaaaagacgatgtaagctcaaataaactttagaaaagacgatgtaagctcaaataacaaaaaatatgtaagctgttttagaaacaaaaaagc | 93 | gttgcaataaactttagaaaagacgatgtaagctcaaataacaaaagtaca | 51 |
|  | agaaaagacgatgtaagctcaaataaactttaatagccaataacaaaaaatatgtaagctgttttagaaacaaaaaagc | 79 | gttgcaataaactttaatagccaataacaaaagtaca | 37 |
|  | agaaaagacgatgttttagaaaacaaaaagctcaaataaactttaataacaaaaaatatgtaagctgttttagaaacaaaaaagc | 85 | gttggaataaactttaataacaaaagtaca | 30 |
|  | agaaaagacgatgtaagctcaaataaactttacgatgtaagctcaaataacaaaaaatatgtaagctgttttagaaacaaaaaagc | 86 | gttgcaataaactttacgatgtaagctcaaataacaaaagtaca | 44 |
|  | aagctgttttacgatgttttagaaaacaaaaagctcaaataaacaaatttacgatgtaagctcaaataacaaaaaatatgtaagctgttttagaaacaaaaaagc | 105 | aagctgttttacgatgttttagaaaacaaaaagctcaaataaacaaatttacgatgtaagctcaaataacaaaagtaca | 79 |
|  | aagctgttttacgatgttttagaaaacaaaaagctcaaataaacaaatttacgatgtaagctcaaataacaaaaaatatgtaagctgttttagaaacaaaaaagc | 105 | aagctgttttacgatgttttagaaaacaaaaagctcaaataaacaaatttacgatgtaagctcaaataacaaaagtaca | 79 |
|  | aagctgttttacgatgtaagctcaaataaacaaatttaataaactttaataacaaaaaataacaaaaaatatgtaagctgttttagaaacaaaaaagc | 98 | aagctgttttacgatgtaagctcaaataaacaaatttaataaactttaataacaaaaaataacaaaagtaca | 72 |
|  | aagctgttttacgatgtaagctcaaataaacaaatttacgatgttttagaaaacaaaaagctcaaataacaaaaaatatgtaagctgttttagaaacaaaaaagc | 105 | aagctgttttacgatgtaagctcaaataaacaaatttacgatgttttagaaaacaaaaagctcaaataacaaaagtaca | 79 |
|  | aagctgttttaggtaacacgatgtaagctcaaataaacaaatttaggtaacacgatgtaagctcaaataacaaaaaatatgtaagctgttttagaaacaaaaaagc | 106 | aagctgttttaggtaacacgatgtaagctcaaataaacaaatttaggtaacacgatgtaagctcaaataacaaaagtaca | 80 |
|  | aagctgttttaataaacaaatttaagctcaaataacaaaaaatatgtaagctgttttagaaacaaaaaagc | 71 | aagctgttttaataaacaaatttaagctcaaataacaaaagtaca | 45 |
|  | aagctgttttaataaacaaatttaataacaaaaaatatgtaagctgttttagaaacaaaaaagc | 64 | aagctgttttaataaacaaatttaataacaaaagtaca | 38 |
|  | aagctgttttaagctcaaataaacaaatttacgatgttttagaaaacaaaaagctcaaataacaaaaaatatgtaagctgttttagaaacaaaaaagc | 98 | aagctgttttaagctcaaataaacaaatttacgatgttttagaaaacaaaaagctcaaataacaaaagtaca | 72 |
|  | aagctgttttacgctgtaagctcaaataaacaaatttacgccatttgtaagctcaaataacaaaaaatatgtaagctgttttagaaacaaaaaagc | 96 | aagctgttttgtttcaataaacaaatttacgccatttgtaagctcaaataacaaaagtaca | 61 |
|  | aagctgttttaagctcaaataaacaaatttaataacaaaaaatatgtaagctgttttagaaacaaaaaagc | 71 | aagctgttttaagctcaaataaacaaatttaataacaaaagtaca | 45 |
|  | aagctcaaataaactttaagctcaaataacaaaaaatatgtaagctgttttagaaacaaaaaagc | 65 | aagctcaaataaactttaagctcaaataacaaaagtaca | 39 |
|  | aagctcaaataaactttaataaacaataacaaaaaatatgtaagctgttttagaaacaaaaaagc | 65 | aagctcaaataaactttaataaacaataacaaaagtaca | 39 |
|  | aagcacaacgatgtaagctcaaataaactttaagcacaacgatgtaagctcaaataacaaaaaatatgtaagctgttttagaaacaaaaaagc | 93 | aagcacaacgatgtaagctcaaataaactttaagcacaacgatgtaagctcaaataacaaaagtaca | 67 |
|  | aagctcatttaagctgtaaaaataaactttaagctcaaataacaaaaaatatgtaagctgttttagaaacaaaaaagc | 78 | aagctcatttaagctgtaaaaataaactttaagctcaaataacaaaagtaca | 52 |
|  | aagctcatttaagctgtaaaaataaactttaagctcatttaagcttaaaaaataacaaaaaatatgtaagctgttttagaaacaaaaaagc | 91 | aagctcatttaagctgtaaaaataaactttaagctcatttaagcttaaaaaataacaaaagtaca | 65 |
|  | aagctcaaataaactttacgaaacacgatgtaagctcaaataacaaaaaatatgtaagctgttttagaaacaaaaaagc | 79 | aagctcaaataaactttgtaccaataacaaaagtaca | 37 |
|  | aagctcaaataaactttaataacaaaaaatatgtaagctgttttagaaacaaaaaagc | 58 | aagctcaaataaactttaataacaaaagtaca | 32 |
|  | aagctcaaataaactttaataacaaaaaatatgtaagctgttttagaaacaaaaaagc | 58 | aagctcaaataaactttaataacaaaagtaca | 32 |
|  | aagctcaaataaactttacgccatttgtaagctcaaataacaaaaaatatgtaagctgttttagaaacaaaaaagc | 76 | aagctcaaataaactttacgccatttgtaagctcaaataacaaaagtaca | 50 |
|  | agaaaagacgatctaagctctacgatgtaaga | 32 |  |  |
|  | acgcaactttagaaaagaaaacgatctaagctctacgatgtaaga | 45 |  |  |
|  | acgaaactttagaaaagaaaacgatctaagctctacgatgtaaga | 45 |  |  |
|  | agaaaagacgatctacgctctacgaataacgatgtaaga | 39 |  |  |
|  | acgatctacgctctacgaataacgatgtaaga | 32 |  |  |
|  | acgctctacgaataacgatgtaaga | 25 |  |  |
|  | acgaataacgatgtaagaccccagtcaggcctaacgta | 38 |  |  |
|  | acgatgtaagaccccagtcaggcctaacgta | 31 |  |  |
|  | agaaaagacgatctacgctctacga | 25 |  |  |
|  | agaaaagacgatctacgcccccagtcaggcctaacgta | 38 |  |  |
|  | agaaaagacgaccccagtcaggcctaacgta | 31 |  |  |
