## Supplementary material for "DNA Encoded Glycan Libraries as a next-generation tool for the study of glycan-protein interactions": GLYCODER: carb_nuc.py

```
# To get dictionary containing monosaccharide, linkages, modifiactions, core glycan structures and their nucelotide codes.
def get_dict_for_key_val(keys,values):
    new_dict = {}
    key_index = 0
    max_possible = len(keys)
    for value in values:
        if key_index < max_possible:
            new_dict[keys[key_index]] = value
            key_index += 1
    return new_dict

carb_keys_list = ['All','AllNAc','AllN','AllA','Alt','AltNAc','AltN','AltA','Glc','GlcNAc','GlcN','GlcA','Man','ManNAc','ManN','ManA','Gul','GulNAc','GulN','GulA','Ido','IdoNAc','IdoN','IdoA','Gal','GalNAc','GalN','GalA','Tal','TalNAc','TalN','TalA','Fuc','FucNAc','FucN','Qui','QuiNAc','QuiN','Rha','RhaNAc','RhaN','NeuAc','NeuGc','Kdn','Kdo','Hep','Fru','Sor','Psi','Tag','Rib','Ara','Xyl','Lyx','R']
carb_val_list = ['AAAA','AAAC','AAAG','AAAT','AACA','AACC','AACG','AACT','AAGA','AAGC','AAGG','AAGT','AATA','AATC','AATG','AATT','ACAA','ACAC','ACAG','ACAT','ACCA','ACCC','ACCG','ACCT','ACGA','ACGC','ACGG','ACGT','ACTA','ACTC','ACTG','ACTT','AGAA','AGAC','AGAG','AGAT','ATAA','ATAC','ATAG','ATAT','ATCA','AGGC','AGGG','AGGT','ATCC','ATCG','ATGA','ATGC','ATAA','ATAC','GGGG','ATAT','ATCA','ATCT','CCCCAGTCAGGCCTAACGTA','ATCG','ATCT','ATGG','ATGT','ATTA','ATTC','ATTG','ATTT','CAAA','CAAC','CAAG','CAAT','CACA','CACC','CACG','CACT','CCAA','CCAC','CCAG','CCAT','CCCC','CCCG','CCCT','CCGA','CCGC','CCGG','CCGT','CCTA','CCTC','CCTG','CCTT','CGCA','CGCC','CGCG','CGCT','CTCA','CTCC','CTCG','CTCT','GAAA','GAAC','GAAG','GAAT','GACA','GACC','GACG','GACT']
carb_dict = get_dict_for_key_val(carb_keys_list,carb_val_list)
#print carb_dict

link_keys_list = ['a1-1','a1-2','a1-3','a1-4','a1-5','a1-6','a1-7','a1-8','a2-1','a2-2','a2-3','a2-4','a2-5','a2-6','a2-7','a2-8','b1-1','b1-2','b1-3','b1-4','b1-5','b1-6','b1-7','b1-8','a?','b?','?1','?2','?3','?4','?5','?6','?7','?8','??',')','(']
link_val_list=['AAT','AAG','AAC','ATA','AGA','ACA','ATT','ATC','GTA','GTC','GGT','GGC','GGA','GTT','GAC','GAT','ATG','TCA','TCT','TGT','TCC','TTA','TTC','GAA','GCC','CAA','CAC','CTC','CCA','GGG','CCT','AGG','CCG','CGC','AGT','AAA','TTT']
link_dict=get_dict_for_key_val(link_keys_list, link_val_list)
#print link_dict

mod_keys_list = ['[2Y]','[4Y]','[7Y]','[8Y]','[?Y]','[2OH]','[4OH]','[7OH]','[8OH]','[?OH]','[2V]','[4V]','[7V]','[8V]','[?V]','[2S]','[4S]','[7S]','[8S]','[?S]','[2P]','[4P]','[7P]','[8P]','[?P]','[2J]','[4J]','[7J]','[8J]','[?J]','[2NAc]','[4NAc]','[7NAc]','[8NAc]','[?NAc]','[2Ac]','[4Ac]','[7Ac]','[8Ac]','[?Ac]','[2COOH]','[4COOH]','[7COOH]','[8COOH]','[?COOH]','[2IN]','[4IN]','[7IN]','[8IN]','[?IN]','[2EE]','[4EE]','[7EE]','[8EE]','[?EE]','[2EH]','[4EH]','[7EH]','[8EH]','[?EH]','[2Q]','[4Q]','[7Q]','[8Q]','[?Q]','[2QS]','[4QS]','[7QS]','[8QS]','[?QS]','[2PYR]','[4PYR]','[7PYR]','[8PYR]','[?PYR]','[2ECO]','[4ECO]','[7ECO]','[8ECO]','[?ECO]','[2PC]','[4PC]','[7PC]','[8PC]','[?PC]','[2PE]','[4PE]','[7PE]','[8PE]','[?PE]','[2ME]','[3ME]','[4ME]','[7ME]','[8ME]','[?ME]','[3S]']
mod_val_list = ['AGCA','AGCC','AGCG','AGCT','CCCA','AGTA','AGTC','AGTG','AGTT','CAGA','CAGC','CAGG','CAGT','CATA','CATC','CATG','CATT','CGAA','CGAC','CGAG','CGAT','CGGA','CGGC','CGGG','CGGT','CGTA','CGTC','CGTG','CGTT','CTAA','CTAC','CTAG','CTAT','CTGA','CTGC','CTGG','CTGT','CTTA','CTTC','CTTG','CTTT','GAGA','GAGC','GAGG','GAGT','GCAA','GCAC','GCAG','GCAT','GCGA','GCGC','GCGG','GCGT','GCTA','GCTC','GCTG','GCTT','GTGA','GTGC','GTGG','GTGT','TAAA','TAAC','TAAG','TAAT','TACA','TACC','TACG','TACT','TAGA','TAGC','TAGG','TAGT','TATA','TATC','TATG','TATT','TCGA','TCGC','TCGG','TCGT','TGAA','TGAC','TGAG','TGAT','TGCA','TGCC','TGCG','TGCT','TGGA','TGGC','TGGG','TGGT','TTGA','TTGC','TTGG','TTGT']
mod_dict = get_dict_for_key_val(mod_keys_list,mod_val_list)
#print mod_dict

core_keys_list = ['AATATGTAAGCTGTTTTAGAAACAAAAAAGC','AATAACAAATATGTAAGCTGTAAGC','ACGATGTAAGCTGT','ACGAAACACGATGTAAGCTCA','ACGAAACACGATGTAAGCTCT','ACGATCTAAGCTGT','AATAAACAATAACA','ACGCTGTAAGCTCA','AGAAAACAAGCTCA','AGAAAAGACGATGTTTTAGAAAACAAAAAGCTCA','AGAAAAGACGATGTAAGCTCA','ACGATGTTTTAGAAAACAAAAAGCTGT','ACGCTGTAAGCTTA','ACGATGTAAGCTTA','AGGCGGTACGATGTAAGCTGT','AGGCGGTACGATGTAAGCTTA']
core_val_list = ['GTACA','GTACT','GTCCT','GTACC','GTTAT','GTTCA','GGATT','GTTTC','GTTGT','GTTGG','GTTGC','GTGAG','GTGTC','GTGCG','GTGGA','GTGGT']
core_dict = get_dict_for_key_val(core_keys_list, core_val_list)
#print core_dict, len(core_dict)

# To get reverse dictionary from the above dictionary.
def get_rev_dict(dict):
    dict_2 ={}
    for key in dict:
        dict_2[dict[key]] = key
    return dict_2

carb_dict_2 = get_rev_dict(carb_dict)
#print carb_dict_2

link_dict_2 = get_rev_dict(link_dict)
#print link_dict_2

mod_dict_2 = get_rev_dict(mod_dict)
#print mod_dict_2

core_dict_2 = get_rev_dict(core_dict)
#print core_dict_2


# To get a list of nucleotide code from the given glycan.
def get_nuc(carb):
    carb_list = carb.split(' ')
    seq =[]
    nuc = ''
    for i in carb_list:
        if carb_dict.has_key(i):
            seq.append(carb_dict.get(i))
        #print seq
        if link_dict.has_key(i):
            seq.append(link_dict.get(i))
        if mod_dict.has_key(i):
            seq.append(mod_dict.get(i))
        #print seq
    nuc = ''.join(seq)
    return nuc

carb = ("Rha")
nuc = get_nuc(carb)
print get_nuc(carb)

# To increase the length of short glycan
def get_mono(nuc):
    if len(nuc)< 20:
        new_nuc_str = ''
        nuc_new = [nuc]
        #print '+++', nuc_new
        R = 'CCCCAGTCAGGCCTAACGTA'
        nuc_new.append(R)
        new_nuc_str = ''.join(nuc_new)
        return new_nuc_str

print get_mono(nuc), len(get_mono(nuc))

# To get glycan from when the nucleotide is given.
def get_carb(nuc):
    if len(nuc)> 38:
        seq_1 = []
        i = 0
        #print "###",i+4,len(nuc_str)
        while i+4 <= len(nuc):
            a = nuc[i:i+4]
            seq_1.append(carb_dict_2.get(a))
            i = i+4
            #print "----->",a, carb_dict_2.get(a)
            if i+3 < len(nuc):
                a = nuc[i:i+3]
                if link_dict_2.has_key(a):
                    seq_1.append(link_dict_2.get(a))
                    #print "!!!!",a,link_dict_2.get(a)
                elif i+4 < len(nuc):
                     a = nuc[i:i+4]
                     seq_1.append(mod_dict_2.get(a))
                     #print "(((",a,mod_dict_2.get(a)
                     i = i+4
                     a = nuc[i:i+3]
                     seq_1.append(link_dict_2.get(a))
                     #print ")))",a,link_dict_2.get(a)
            i = i+3
            a = nuc[i:i+3]
            #print "+=+=",a
            if a == 'AAA' or a == 'TTT':
                seq_1.append(link_dict_2.get(a))
                #print  "@@@>" , a,link_dict_2.get(a)
                i = i+3
                a = nuc[i:i+3]
                if a == 'AAA' or a =='TTT':
                    seq_1.append(link_dict_2.get(a))
                    #print "%%%%",a,link_dict_2.get(a)
            i = i+3
    elif len(nuc)< 38:
        seq_1 = []
        i = 0
        while i+4 <= len(nuc):
            a = nuc[i:i+4]
            seq_1.append(carb_dict_2.get(a))
            i = i+4
            #print "----->",a, carb_dict_2.get(a)
            if i+3 < len(nuc):
                a = nuc[i:i+3]
                print '+++',a
                if a == 'CCC':
                    print 'CCC'
                    seq_1.append('R')
                    i = len(nuc)
                elif link_dict_2.has_key(a):
                    seq_1.append(link_dict_2.get(a))
                    i = i+3

        return seq_1
                    
print get_carb(nuc)

# To get short sequence from long sequence
def get_short_nuc(nuc, core_dict):
    short_nuc = ""
    long_rev_sorted_keys=sorted(core_dict.keys(),reverse=True)
    #print long_rev_sorted_keys, len(long_rev_sorted_keys)
    rem_str = nuc
    no_more_to_replace = False
    #print "+++" +rem_str
    while ( len(rem_str) > 0 and not no_more_to_replace):
        i=0
        for long_carb in long_rev_sorted_keys:
            i =i+1
            index= rem_str.find(long_carb)
            #print rem_str.find(long_carb)
            if index >=0:
                #print index
                split_index = index+len(long_carb)
                #print split_index
                temp=rem_str[0:split_index]
                #print "+++" +temp
                rem_str = rem_str[split_index:]
                #print "+++" + temp.replace(long_carb,core_dict[long_carb])
                short_nuc = short_nuc + temp.replace(long_carb,core_dict[long_carb])
                #print "#### "+short_nuc
                #print "+++" +nuc_str
        if i == len(long_rev_sorted_keys) :
            no_more_to_replace = True
                
    return short_nuc
my_short_nuc = get_short_nuc(nuc, core_dict)
print my_short_nuc, len(my_short_nuc)

# To get long sequence from short sequence
def get_long_nuc_from_short(short_nuc, core_dict_2):
    long_nuc = ""
    long_rev_act_sorted_keys=sorted(core_dict_2.keys(),reverse=True)
    #print long_rev_act_sorted_keys, len(long_rev_act_sorted_keys)
    rem_str = short_nuc
    no_more_to_replace = False
    while ( len(rem_str) > 0 and not no_more_to_replace):
        i=0
        for short_carb in long_rev_act_sorted_keys:
            i = i+1
            index = rem_str.find(short_carb)
            if index >= 0:
                 #print index
                 split_index = index + len(short_carb)
                 #print split_index
                 temp = rem_str[0:split_index]
                 #print "+++" +temp
                 rem_str = rem_str[split_index:]
                 #print "+++" + temp.replace(short_carb,core_dict_2[short_carb])
                 long_nuc = long_nuc + temp.replace(short_carb, core_dict_2[short_carb])
                 #print "#### "+short_nuc
                 #print "+++" +nuc_str
        if i == len(long_rev_act_sorted_keys):
            no_more_to_replace = True

    return long_nuc

my_long_nuc= get_long_nuc_from_short(my_short_nuc, core_dict_2)
print my_long_nuc, len(my_long_nuc)
```
