## Supplementary material for "DNA Encoded Glycan Libraries as a next-generation tool for the study of glycan-protein interactions": GLYCODER: NGS_data_summary (4).ipynb

```
# %% [code]
# Import the libraries in python

import pandas as pd
import sys
import os
from collections import OrderedDict

# %% [code]
# Define the sequences to analyze

keys = ['AATGATACGGCGACCACCGAAGAAAAGACGATCTACGCTCTACGAATAACGATGTAAGACACGTCTGAACTCCAGTCAC',
             'GTGACTGGAGTTCAGACGTGTCTTACATCGTTATTCGTAGAGCGTAGATCGTCTTTTCTTCGGTGGTCGCCGTATCATT',
             'AATGATACGGCGACCACCGAACGATCTACGCTCTACGAATAACGATGTAAGACACGTCTGAACTCCAGTCAC',
             'GTGACTGGAGTTCAGACGTGTCTTACATCGTTATTCGTAGAGCGTAGATCGTTCGGTGGTCGCCGTATCATT',
            'AATGATACGGCGACCACCGAACGCTCTACGAATAACGATGTAAGACACGTCTGAACTCCAGTCAC',
            'GTGACTGGAGTTCAGACGTGTCTTACATCGTTATTCGTAGAGCGTTCGGTGGTCGCCGTATCATT',
            'AATGATACGGCGACCACCGAACGAATAACGATGTAAGACCCCAGTCAGGCCTAACGTACACGTCTGAACTCCAGTCAC',
            'GTGACTGGAGTTCAGACGTGTACGTTAGGCCTGACTGGGGTCTTACATCGTTATTCGTTCGGTGGTCGCCGTATCATT',
            'AATGATACGGCGACCACCGAACGATGTAAGACCCCAGTCAGGCCTAACGTACACGTCTGAACTCCAGTCAC',
             'GTGACTGGAGTTCAGACGTGTACGTTAGGCCTGACTGGGGTCTTACATCGTTCGGTGGTCGCCGTATCATT',
            'AATGATACGGCGACCACCGAAGAAAAGACGATCTACGCTCTACGACACGTCTGAACTCCAGTCAC',
            'GTGACTGGAGTTCAGACGTGTCGTAGAGCGTAGATCGTCTTTTCTTCGGTGGTCGCCGTATCATT',
            'AATGATACGGCGACCACCGAAGAAAAGACGATCTACGCCCCCAGTCAGGCCTAACGTACACGTCTGAACTCCAGTCAC',
            'GTGACTGGAGTTCAGACGTGTACGTTAGGCCTGACTGGGGGCGTAGATCGTCTTTTCTTCGGTGGTCGCCGTATCATT',
            'AATGATACGGCGACCACCGAAGAAAAGACGACCCCAGTCAGGCCTAACGTACACGTCTGAACTCCAGTCAC',
            'GTGACTGGAGTTCAGACGTGTACGTTAGGCCTGACTGGGGTCGTCTTTTCTTCGGTGGTCGCCGTATCATT',
            'AATGATACGGCGACCACCGAACGCAACTTTAGAAAAGAAAACGATCTAAGCTCTACGATGTAAGACACGTCTGAACTCCAGTCAC',
            'GTGACTGGAGTTCAGACGTGTCTTACATCGTAGAGCTTAGATCGTTTTCTTTTCTAAAGTTGCGTTCGGTGGTCGCCGTATCATT',
            'AATGATACGGCGACCACCGAACGAAACTTTAGAAAAGAAAACGATCTAAGCTCTACGATGTAAGACACGTCTGAACTCCAGTCAC',
            'GTGACTGGAGTTCAGACGTGTCTTACATCGTAGAGCTTAGATCGTTTTCTTTTCTAAAGTTTCGTTCGGTGGTCGCCGTATCATT',
            'AATGATACGGCGACCACCGAAGAAAAGACGATCTAAGCTCTACGATGTAAGACACGTCTGAACTCCAGTCAC',
            'GTGACTGGAGTTCAGACGTGTCTTACATCGTAGAGCTTAGATCGTCTTTTCTTCGGTGGTCGCCGTATCATT',
       'AATGATACGGCGACCACCGAACGCTCTACGAATAACGATGTAAGACCCCAGTACCGTAATGCTGACTGCACTGGACTTGACACGTCTGAACTCCAGTCAC',
       'GTGACTGGAGTTCAGACGTGTCAAGTCCAGTGCAGTCAGCATTACGGTACTGGGGTCTTACATCGTTATTCGTAGAGCGTTCGGTGGTCGCCGTATCATT']
words = OrderedDict.fromkeys(keys,0)

# %% [code]
DF = pd.DataFrame()
finDF = pd.DataFrame()

# %% [code]
fullpath = '/Users/h264632/Desktop/1234' # the path to the directory containing fastaq files
for filename in os.listdir(fullpath):
    if filename.endswith('.fastaq'): continue
    fullname = os.path.join(fullpath, filename)
    with open(fullname) as f:
        data = f.read()
        for key,val in words.items():
            ct = data.count(key)
            words[key] = ct
            DF = pd.DataFrame(words,index=[filename]).transpose()
    finDF = pd.concat([finDF,DF], axis=1)

# %% [code]
# Write the final analysis to a csv file named NGS_data_summary_without_sorting_24_Sequences_new.csv

finDF.to_csv('/Users/h264632/Desktop/1234/NGS_data_summary_without_sorting_24_Sequences_new.csv')
```
