## Supplementary material for "DNA Encoded Glycan Libraries as a next-generation tool for the study of glycan-protein interactions": Raw data

### Run Summary

| 428 M | 92 |
|---|---|
| Total Bases | Key Signal |

**86%**
ISP Loading

##### ISP Density

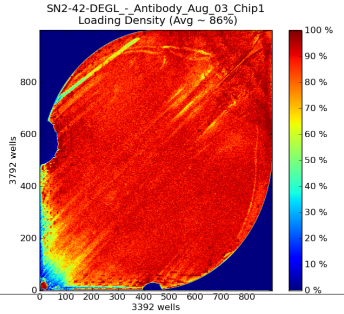

**6,231,567**
Total Reads

**64%**
Usable Reads

##### ISP Summary

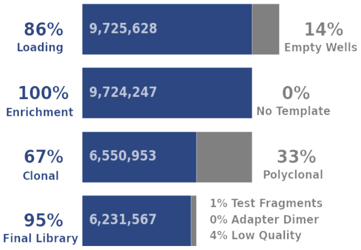

| 69 bp | 65 bp | 64 bp |
|---|---|---|
| Mean | Median | Mode |

##### Read Length

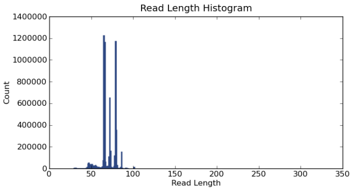

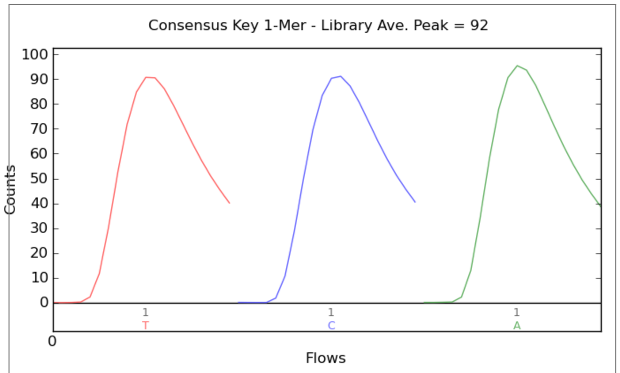

| **Addressable Wells** | **11,287,275** | |
|---|---|---|
| With ISPs | 9,725,628 | 86.2% |
| Live | 9,724,247 | 100.0% |
| Test Fragment | 41,071 | 00.4% |
| Library | 9,683,176 | 99.6% |

| **Library ISPs** | **9,683,176** | |
|---|---|---|
| Filtered: Polyclonal | 3,173,294 | 32.8% |
| Filtered: Low Quality | 278,199 | 02.9% |
| Filtered: Adapter Dimer | 116 | 00.0% |
| **Final Library ISPs** | **6,231,567** | **64.4%** |

| Barcode Name | Sample | Bases | $\geq Q20$ | Reads | Mean Read Length |
|---|---|---|---|---|---|
| No barcode | none | 14,323,561 | 11,630,454 | 191,461 | 75 bp |
| IonXpress_001 | Sample 1 | 18,328,993 | 17,792,934 | 270,224 | 68 bp |
| IonXpress_002 | Sample 2 | 19,315,245 | 18,763,609 | 284,877 | 68 bp |
| IonXpress_003 | Sample 3 | 21,081,191 | 20,449,423 | 309,350 | 68 bp |
| IonXpress_004 | Sample 4 | 18,689,630 | 18,125,494 | 274,749 | 68 bp |
| IonXpress_005 | Sample 5 | 18,234,845 | 17,698,836 | 266,593 | 68 bp |
| IonXpress_006 | Sample 6 | 17,853,084 | 17,203,782 | 263,020 | 68 bp |
| IonXpress_007 | Sample 7 | 18,889,993 | 18,117,430 | 289,432 | 65 bp |
| IonXpress_008 | Sample 8 | 17,666,792 | 16,941,592 | 270,576 | 65 bp |
| IonXpress_009 | Sample 9 | 15,117,761 | 14,583,530 | 230,380 | 66 bp |
| IonXpress_010 | Sample 10 | 18,736,168 | 18,025,091 | 285,620 | 66 bp |
| IonXpress_011 | Sample 11 | 18,965,517 | 18,222,621 | 289,713 | 65 bp |

iontorrent by Thermo Fisher Scientific

| | | | | | |
|---|---|---|---|---|---|
| IonXpress_012 | Sample 12 | 19,211,895 | 18,510,258 | 293,970 | 65 bp |
| IonXpress_013 | Sample 13 | 15,797,462 | 14,994,772 | 223,312 | 71 bp |
| IonXpress_014 | Sample 14 | 15,667,558 | 14,956,962 | 219,574 | 71 bp |
| IonXpress_015 | Sample 15 | 15,059,418 | 14,298,519 | 220,949 | 68 bp |
| IonXpress_016 | Sample 16 | 15,809,932 | 14,985,257 | 233,543 | 68 bp |
| IonXpress_017 | Sample 17 | 15,813,730 | 14,972,184 | 234,220 | 68 bp |
| IonXpress_018 | Sample 18 | 15,127,749 | 14,412,197 | 222,060 | 68 bp |
| IonXpress_019 | Sample 19 | 16,420,716 | 15,639,191 | 228,037 | 72 bp |
| IonXpress_020 | Sample 20 | 14,353,816 | 13,734,241 | 198,627 | 72 bp |
| IonXpress_021 | Sample 21 | 16,098,742 | 15,320,078 | 223,281 | 72 bp |
| IonXpress_022 | Sample 22 | 17,362,095 | 16,506,639 | 240,528 | 72 bp |
| IonXpress_023 | Sample 23 | 15,669,188 | 14,977,098 | 215,610 | 73 bp |
| IonXpress_024 | Sample 24 | 18,232,820 | 17,381,379 | 251,857 | 72 bp |

| Test Fragment | Reads | Percent 50AQ17 | Read Length Histogram |
|---|---|---|---|
| **TF_C** | **8,625** | **94%** | 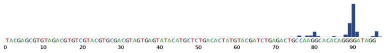 |
| **TF_1** | **30,132** | **96%** | 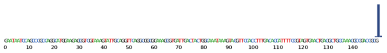 |

iontorrent
by Thermo Fisher Scientific

### Alignment Summary *(aligned to )*

| **0** | |
|---|---|
| Total Alignment Bases | Average Coverage Depth of Reference |

alignment_rate_plot.png

| | Count | % |
|---|---|---|
| **Total Reads** | 0 | – |
| **Aligned Reads** | 0 | % |

**%**

Mean Raw Accuracy 1x

base_error_plot.png

**Alignment Quality**

| | AQ17 | AQ20 | Perfect |
|---|---|---|---|
| **Total Number of Bases [Mbp]** | 0 | 0 | 0 |
| **Mean Length [bp]** | 0 | 0 | 0 |
| **Longest Alignment [bp]** | 0 | 0 | 0 |
| **Mean Coverage Depth** | | | |

Filtered_Alignments_Q10.png

Filtered_Alignments_Q17.png

Filtered_Alignments_Q20.png

Filtered_Alignments_Q47.png

iontorrent
by Thermo Fisher Scientific

#### Analysis Details

| | |
|---|---|
| **Run Name** | R_2015_01_30_21_43_02_user_SN2-42-DEGL_-_Antibody_Aug_03_Chip1 |
| **Run Date** | Jan. 30, 2015, 9:43 p.m. |
| **Run Flows** | 500 |
| **Projects** | DEGL |
| **Sample** | Sample_18, Sample_19, Sample_14, Sample_15, Sample_16, Sample_17, Sample_10, Sample_11, Sample_12, Sample_13, Sample_24, Sample_21, Sample_20, Sample_23, Sample_22, Sample_8, Sample_9, Sample_6, Sample_7, Sample_4, Sample_5, Sample_2, Sample_3, Sample_1 |
| **Reference** | |
| **Instrument** | sn274670958 |
| **Flow Order** | TACGTACGTCTGAGCATCGATCGATGTACAGC |
| **Library Key** | TCAG |
| **TF Key** | ATCG |
| **Chip ID** | AB0084403 |
| **Chip Check** | Passed |
| **Chip Type** | 318C |
| **Chip Data** | single |
| **Barcode Set** | IonXpress |
| **Analysis Name** | Auto_user_SN2-42-DEGL_-_Antibody_Aug_03_Chip1_105 |
| **Analysis Date** | Aug. 3, 2018, 4:39 p.m. |
| **Analysis Flows** | 500 |
| **runID** | JS6JY |
| **BeadFind Args** | justBeadFind |
| **Analysis Args** | Analysis –from-beadfind –use-alternative-etbR-equation –mixed-first-flow 12 –mixed-last-flow 120 |
| **Pre-BaseCaller Args for calibration** | BaseCaller –barcode-filter 0.01 –barcode-filter-minreads 20 |
| **Calibration Args** | Calibration |
| **BaseCaller Args** | BaseCaller –barcode-filter 0.01 –barcode-filter-minreads 20 –phred-table-file /opt/ion/config/phredTable.314.B5.h5 |
| **Alignment Args** | tmap mapall ... stage1 map4 |
| **IonStats Args** | ionstats alignment |
| **Analysis Parameters** | custom |

#### Chef Summary

| |
|---|
| **Ion Chef was not used for this run** |

#### Software Version

| | |
|---|---|
| **Torrent_Suite** | 5.0.5 |
| **host** | 5R0VV12 |
| **ion-analysis** | 5.0.13-1 |
| **ion-chefupdates** | 5.0.7 |
| **ion-dbreports** | 5.0.34-1 |
| **ion-gpu** | 5.0.0-1 |
| **ion-pipeline** | 5.0.17-1 |
| **ion-plugins** | 5.0.28-1 |
| **ion-torrentr** | 5.0.0-1 |
| **Script** | 22.0.0 |
| **LiveView** | 643 |
| **DataCollect** | 488 |
| **OS** | 21 |
| **Graphics** | 36 |

iontorrent
by Thermo Fisher Scientific
