## Supplementary material for "DNA Encoded Glycan Libraries as a next-generation tool for the study of glycan-protein interactions": Raw data

### Run Summary

**347 M** Total Bases | **94** Key Signal

**84%** ISP Loading

**ISP Density**

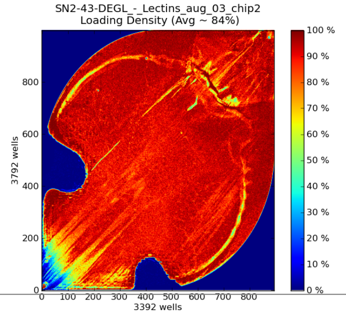

**4,766,527** Total Reads

**50%** Usable Reads

**ISP Summary**

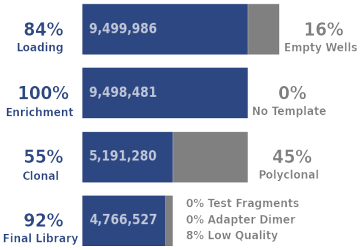

**73 bp** Mean | **71 bp** Median | **85 bp** Mode

**Read Length**

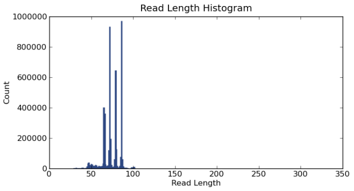

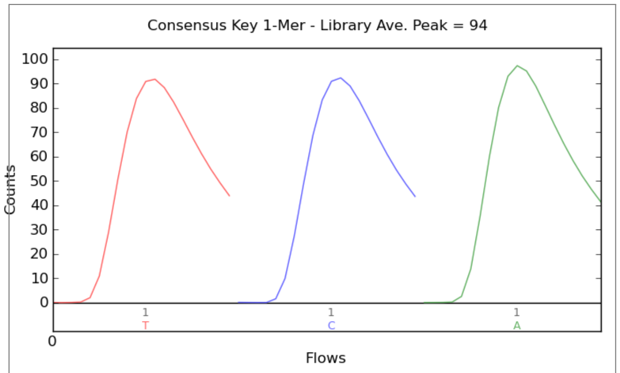

| **Addressable Wells** | **11,287,275** | |
|---|---|---|
| With ISPs | 9,499,986 | 84.2% |
| Live | 9,498,481 | 100.0% |
| Test Fragment | 25,135 | 00.3% |
| Library | 9,473,346 | 99.7% |

| **Library ISPs** | **9,473,346** | |
|---|---|---|
| Filtered: Polyclonal | 4,307,201 | 45.5% |
| Filtered: Low Quality | 399,609 | 04.2% |
| Filtered: Adapter Dimer | 9 | 00.0% |
| **Final Library ISPs** | **4,766,527** | **50.3%** |

| Barcode Name | Sample | Bases | $\geq Q20$ | Reads | Mean Read Length |
|---|---|---|---|---|---|
| No barcode | none | 15,322,695 | 12,065,693 | 198,761 | 77 bp |
| IonXpress_001 | Sample 1 | 15,900,069 | 15,326,710 | 189,831 | 84 bp |
| IonXpress_002 | Sample 2 | 15,218,780 | 14,756,312 | 181,590 | 84 bp |
| IonXpress_003 | Sample 3 | 14,729,846 | 14,245,889 | 176,792 | 83 bp |
| IonXpress_004 | Sample 4 | 16,513,616 | 15,940,916 | 198,115 | 83 bp |
| IonXpress_005 | Sample 5 | 17,939,125 | 17,297,692 | 215,419 | 83 bp |
| IonXpress_006 | Sample 6 | 15,975,531 | 15,382,644 | 192,514 | 83 bp |
| IonXpress_007 | Sample 7 | 11,809,356 | 11,105,589 | 175,688 | 67 bp |
| IonXpress_008 | Sample 8 | 14,201,380 | 13,358,887 | 210,489 | 67 bp |
| IonXpress_009 | Sample 9 | 13,692,246 | 12,951,061 | 195,156 | 70 bp |
| IonXpress_010 | Sample 10 | 12,006,436 | 11,361,019 | 172,567 | 70 bp |
| IonXpress_011 | Sample 11 | 14,317,593 | 13,502,313 | 208,435 | 69 bp |

iontorrent by Thermo Fisher Scientific

| | | | | | |
|---|---|---|---|---|---|
| IonXpress_012 | Sample 12 | 12,288,759 | 11,640,951 | 179,389 | 69 bp |
| IonXpress_013 | Sample 13 | 12,625,160 | 11,883,283 | 176,334 | 72 bp |
| IonXpress_014 | Sample 14 | 15,718,041 | 14,814,143 | 220,403 | 71 bp |
| IonXpress_015 | Sample 15 | 14,956,422 | 14,156,030 | 210,820 | 71 bp |
| IonXpress_016 | Sample 16 | 13,967,089 | 13,193,484 | 198,215 | 70 bp |
| IonXpress_017 | Sample 17 | 13,777,307 | 12,962,946 | 190,906 | 72 bp |
| IonXpress_018 | Sample 18 | 14,357,659 | 13,622,259 | 198,922 | 72 bp |
| IonXpress_019 | Sample 19 | 12,522,110 | 11,971,882 | 187,954 | 67 bp |
| IonXpress_020 | Sample 20 | 10,623,645 | 10,192,555 | 158,679 | 67 bp |
| IonXpress_021 | Sample 21 | 10,720,786 | 10,267,408 | 161,301 | 66 bp |
| IonXpress_022 | Sample 22 | 14,270,232 | 13,590,883 | 214,623 | 66 bp |
| IonXpress_023 | Sample 23 | 11,154,735 | 10,680,788 | 167,538 | 67 bp |
| IonXpress_024 | Sample 24 | 12,448,281 | 11,941,107 | 186,079 | 67 bp |

| Test Fragment | Reads | Percent 50AQ17 | Read Length Histogram |
|---|---|---|---|
| **TF_C** | **4,576** | **93%** | 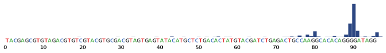 |
| **TF_1** | **18,414** | **94%** | 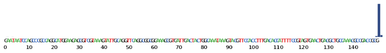 |

### Alignment Summary *(aligned to )*

| 0 | |
|---|---|
| Total Alignment Bases | Average Coverage Depth of Reference |

alignment_rate_plot.png

| | Count | % |
|---|---|---|
| Total Reads | 0 | – |
| Aligned Reads | 0 | % |

| % |
|---|
| Mean Raw Accuracy 1x |

base_error_plot.png

**Alignment Quality**

| | AQ17 | AQ20 | Perfect |
|---|---|---|---|
| **Total Number of Bases [Mbp]** | 0 | 0 | 0 |
| **Mean Length [bp]** | 0 | 0 | 0 |
| **Longest Alignment [bp]** | 0 | 0 | 0 |
| **Mean Coverage Depth** | | | |

iontorrent by Thermo Fisher Scientific

Filtered_Alignments_Q10.png

Filtered_Alignments_Q17.png

Filtered_Alignments_Q20.png

Filtered_Alignments_Q47.png

### Analysis Details

| | |
|---|---|
| **Run Name** | R_2015_01_31_02_17_26_user_SN2-43-DEGL_-_Lectins_aug_03_chip2 |
| **Run Date** | Jan. 31, 2015, 2:17 a.m. |
| **Run Flows** | 500 |
| **Projects** | DEGL |
| **Sample** | Sample_18, Sample_19, Sample_14, Sample_15, Sample_16, Sample_17, Sample_10, Sample_11, Sample_12, Sample_13, Sample_24, Sample_21, Sample_20, Sample_23, Sample_22, Sample_8, Sample_9, Sample_6, Sample_7, Sample_4, Sample_5, Sample_2, Sample_3, Sample_1 |
| **Reference** | |
| **Instrument** | sn274670958 |
| **Flow Order** | TACGTACGTCTGAGCATCGATCGATGTACAGC |
| **Library Key** | TCAG |
| **TF Key** | ATCG |
| **Chip ID** | AB0084458 |
| **Chip Check** | Passed |
| **Chip Type** | 318C |
| **Chip Data** | single |
| **Barcode Set** | IonXpress |
| **Analysis Name** | Auto_user_SN2-43-DEGL_-_Lectins_aug_03_chip2_106 |
| **Analysis Date** | Aug. 3, 2018, 9:15 p.m. |
| **Analysis Flows** | 500 |
| **runID** | R189T |
| **BeadFind Args** | justBeadFind |
| **Analysis Args** | Analysis –from-beadfind –use-alternative-etbR-equation –mixed-first-flow 12 –mixed-last-flow 120 |
| **Pre-BaseCaller Args for calibration** | BaseCaller –barcode-filter 0.01 –barcode-filter-minreads 20 |
| **Calibration Args** | Calibration |
| **BaseCaller Args** | BaseCaller –barcode-filter 0.01 –barcode-filter-minreads 20 –phred-table-file /opt/ion/config/phredTable.314.B5.h5 |
| **Alignment Args** | tmap mapall ... stage1 map4 |
| **IonStats Args** | ionstats alignment |
| **Analysis Parameters** | custom |

### Chef Summary

| |
|---|
| **Ion Chef was not used for this run** |

iontorrent
by Thermo Fisher Scientific

#### Software Version

| | |
|---|---|
| **Torrent_Suite** | 5.0.5 |
| **host** | 5R0VV12 |
| **ion-analysis** | 5.0.13-1 |
| **ion-chefupdates** | 5.0.7 |
| **ion-dbreports** | 5.0.34-1 |
| **ion-gpu** | 5.0.0-1 |
| **ion-pipeline** | 5.0.17-1 |
| **ion-plugins** | 5.0.28-1 |
| **ion-torrentr** | 5.0.0-1 |
| **Script** | 22.0.0 |
| **LiveView** | 643 |
| **DataCollect** | 488 |
| **OS** | 21 |
| **Graphics** | 36 |

iontorrent by Thermo Fisher Scientific
